# An Agentic AI Framework for Ingestion and Standardization of Single-Cell RNA-seq Data Analysis

**DOI:** 10.1101/2025.07.31.667880

**Authors:** Nima Nouri, Ronen Artzi, Virginia Savova

## Abstract

The proliferation of publicly available single-cell RNA sequencing (scRNA-seq) data has created significant opportunities in biomedical research. However, the reuse of these resources is constrained by a series of preparatory steps, including metadata extraction from primary literature, retrieval of datasets from corresponding repositories, and the subsequent manual execution of standardized downstream analysis. These tasks often require manual scripting and rely on fragmented workflows, limiting accessibility and increasing turnaround time. To address these challenges, we designed a two-component system consisting of an artificial intelligence (AI) agent coordinating an automated analysis pipeline. CellAtria (Agentic Triage of Regulated single-cell data Ingestion and Analysis) is an agentic AI framework that enables dialogue-driven, document-to-analysis automation through a chatbot interface. Built on a graph-based, multi-actor architecture, CellAtria integrates a large language model (LLM) with tool-execution capabilities to orchestrate the full lifecycle of data reuse. To support downstream analysis, CellAtria incorporates CellExpress, a co-developed pipeline that applies state-of-the-art scRNA-seq processing steps to transform raw count matrices into analysis-ready single-cell profiles. Thus, CellAtria provides computational skill-agnostic and time-efficient access to standardized single-cell data ingestion and analysis.

## Main

Single-cell RNA sequencing (scRNA-seq) has emerged as a central technique in biomedical research^1, 2, 3, 4^, enabling high-resolution profiling of cellular heterogeneity across tissues, disease states, and therapeutic interventions^5, 6^. Its adoption continues to grow across the pharmaceutical landscape^7, 8^, where it supports applications in target discovery, biomarker stratification, and mechanism-of-action studies^9, 10, 11, 12, 13^. In parallel, public repositories have accumulated an unprecedented volume and diversity of scRNA-seq datasets, spanning multiple species, tissues, and experimental conditions^14, 15, 16^. This rapid expansion presents a unique opportunity to transform fragmented single-cell datasets into structured, decision-support workflows that enhance biological interpretation and accelerate translational research^17, 18^.

Despite the availability of these datasets, integrating published studies into institutional and enterprise-level analysis workflows remains a slow and resource-intensive process (**Figure 1a**). Data ingestion typically begins with manual interpretation of the source publication to establish biological context, followed by the extraction of key metadata - such as tissue type, species, disease condition, and accession identifiers - and culminates in the manual retrieval of datasets from corresponding public repositories. The process then continues with dataset processing and analysis, which often require adaptation to institutional conventions and custom scripting practices. These interpretation- and scripting-based tasks are often delegated to specialists with both domain knowledge and computational proficiency, resulting in high personnel costs and increased susceptibility to analyst-driven errors and inconsistent throughput due to user-dependent variability in single-cell data analysis. Given the repetitive and procedural nature of these tasks, shifting them left - from specialist bioinformaticians to bench scientists - would not only enable faster, more consistent execution of data workflows and reduce dependence on bespoke computational support, but also free up expert capacity for more exploratory or high-impact scientific efforts.

**Figure 1.**
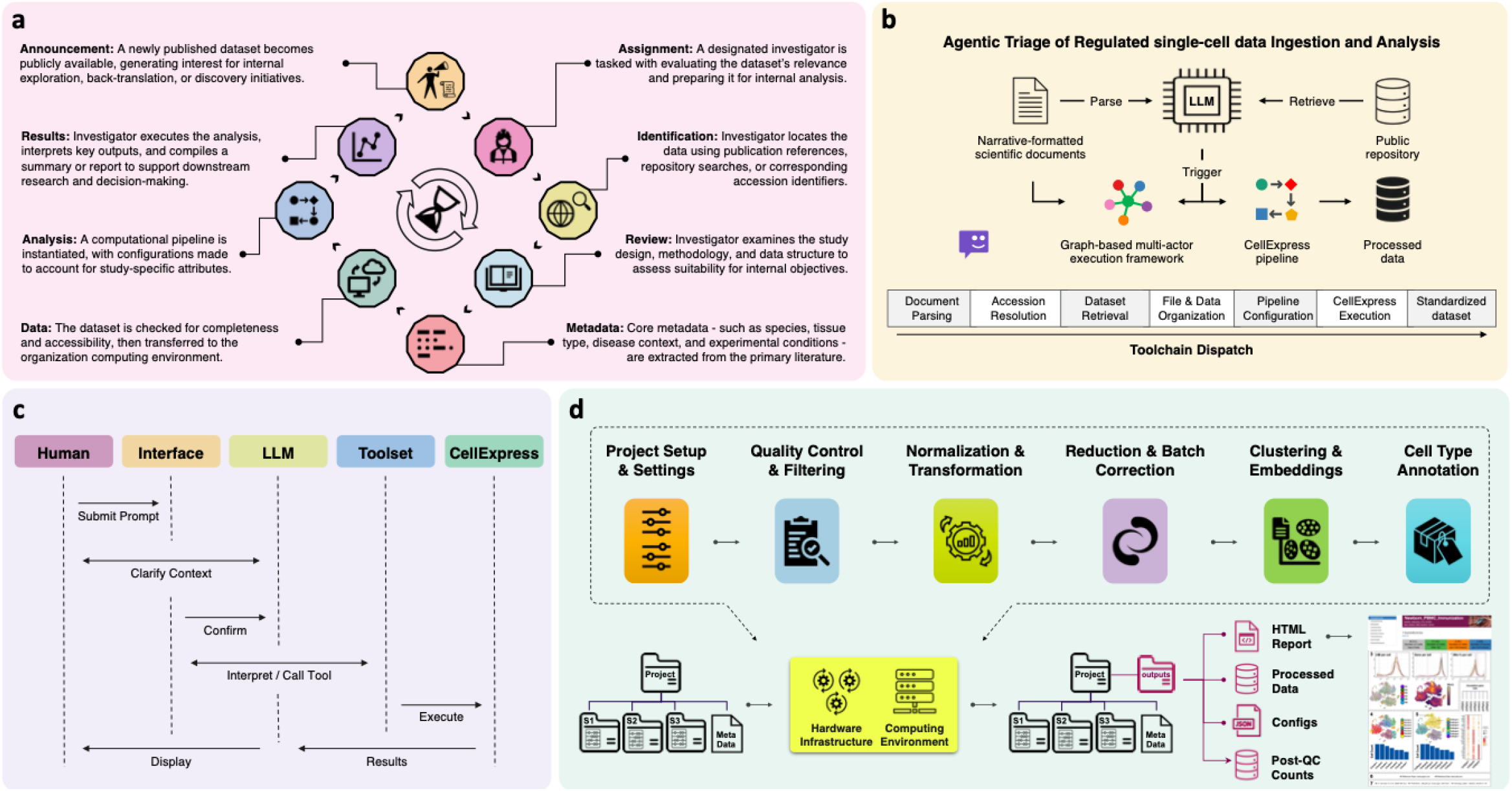
Overview of the CellAtria agentic framework. **a)** Manual data onboarding cycle for public single-cell datasets. This panel illustrates the conventional workflow followed by investigators when integrating newly published datasets into internal research environments. The process spans from dataset announcement and assignment through metadata extraction, data validation, pipeline configuration, and analysis execution. **b)** Architecture of agentic triage and execution. CellAtria employs a large language model (LLM) to mediate task dispatch across a graph-based, multi-actor execution framework, which is integrated with the co-developed CellExpress pipeline. High-level actions such as document parsing, metadata structuring, dataset retrieval, and file organization are coordinated through the LLM-mediated interface and mapped to appropriate backend tools for execution, ultimately producing analysis-ready outputs. **c)** Language model-mediated orchestration of toolchains. Upon receiving a user prompt, the CellAtria interface transfers the request to the LLM agent, which interprets the user’s intent and autonomously invokes relevant tools. Outputs are then returned through the interface, completing a full cycle of context-aware execution. **d)** Structure of the CellExpress pipeline. CellExpress implements a standardized single-cell RNA-seq workflow, encompassing project setup, quality control (QC), normalization, batch correction, dimensionality reduction, clustering, and cell type annotation. This pipeline is fully customizable and seamlessly integrated with the CellAtria orchestration layer, enabling automated execution from raw count matrices to interpretable results. It produces a comprehensive analytical package, including: (1) An interactive HTML report summarizing the workflow with key QC metrics and visualizations; (2) A finalized, fully annotated AnnData object for downstream analyses; (3) A machine-readable configuration export for full auditability; and (4) A QC-filtered AnnData object for alternative usages.

Agentic artificial intelligence (AI) systems^19, 20^ offer a structured approach to automating complex biomedical workflows by coupling large language models (LLMs)^21, 22^ with domain-specific computational toolchains^23^. These systems operate through modular, executable architectures in which predefined computational functions are dynamically composed in response to user prompts and contextual cues. The LLM serves as a semantic interface layer, interpreting natural language prompts and dispatching appropriate computational actions, thereby enabling dialogue-driven interaction with underlying infrastructure. Consequently, agentic AI systems, through their adaptive control flow, autonomous decision-making, and operational self-sufficiency, offer a scalable framework for minimizing manual intervention and democratizing access to bioinformatics infrastructures.

AI agents span a spectrum of capability and autonomy, ranging from basic tools to highly sophisticated systems^24, 25,26^. This range includes reactive agents, which respond to predefined inputs with scripted logic; context-aware agents, which adjust responses based on memory and contextual understanding; and goal-oriented agents, capable of planning and executing tasks to meet specific objectives. Further along this spectrum are adaptive agents, that learn dynamically to refine strategies based on experience, culminating in autonomous agents, characterized by general intelligence, self-learning, and independent decision-making across complex scenarios. This progression reflects the increasing sophistication of agents towards more versatile and adaptable systems.

While some contemporary perspectives propose leveraging LLMs for dynamic, ad-hoc analytical code generation at runtime^27, 28, 29, 30^, such strategies encounter obstacles in practice. The primary challenge with direct code generation for complex single-cell bioinformatics tasks lies in ensuring consistent output and scientific validity^31^. Such methods inherently struggle with reproducibility, as LLM outputs can vary based on model versions and prompting nuances^32^. More critically, the generated code carries a substantial risk of hallucinations^33, 34^, logical errors^35, 36^, incompatibility with state-of-the-art methods or software (due to the LLM’s training data temporal cutoff)^37, 38^, and the misapplication of domain-specific parameters^39^. Any of these issues can lead to scientifically unsound analyses without extensive human oversight^40^ and debugging^41^. Most importantly, in clinical or highly regulated biomedical settings, the lack of protocol standardization inherent in ad-hoc code generation risks violating established quality control and analysis guidelines^42^.

Moving beyond the risks in ad-hoc code-generation strategy, we adopted a critical architectural decision: an LLM-mediated tool-centric paradigm that balances flexibility with the imperative for scientific rigor and reproducibility. Here, the LLM’s role shifts from writing raw code to intelligently orchestrating a library of pre-vetted, robust analytical tools. We developed CellAtria, an agentic AI system that enables end-to-end, document-to-analysis automation in single-cell research (**Figure 1b**). By combining natural language interaction with a graph-based, multi-actor architecture execution framework (**Figure 1c**), CellAtria links tasks ranging from literature parsing, metadata extraction, and dataset retrieval to scRNA-seq processing via its co-developed companion pipeline, CellExpress, which applies state-of-the-art processing steps to transform raw count matrices into analysis-ready single-cell profiles (**Figure 1d**). Utilizing CellAtria user interface (**Supplementary Fig. S1**), researchers interact with a language model that orchestrates pre-validated analytical tools (**Supplementary Fig. S2**), eliminating the need for manual scripting while ensuring standardized, reproducible analyses and accelerating the reuse of public single-cell resources.

## Results

A comprehensive description of the agentic system, CellAtria, including its architectural design and implementation details is provided in the Methods section. To illustrate CellAtria’s capabilities, we implemented three prototypical use cases, each representing a common scenario in single-cell research.

### CellAtria Extracts Metadata from Article URLs and Retrieves Study-Level Datasets

In translational and early discovery research, investigators often encounter newly published studies whose associated datasets are highly relevant to institutional priorities.

To demonstrate CellAtria’s capability for literature-driven data acquisition and analysis execution, we selected a publicly available longitudinal transcriptomic study profiling immune responses in 2-month-old infants following routine vaccination^43^. Upon receiving the article URL, the agent conducts a multi-turn dialogue to parse the manuscript directly from the journal webpage, extract key structured metadata - including sample annotations and accession identifiers - and coordinate dataset retrieval from corresponding public repositories using GSE-level (GEO study-wide) accessioning (**Supplementary Figs. S3-S10**). Following user validation and direction, the agent proceeds to triage these data, executing essential organizational tasks and ensuring strict naming compatibility for seamless integration with downstream analytical functions (**Supplementary Figs. S10-S11**).

We note that this multi-turn interaction established a dynamic feedback loop, wherein the agent proactively surfaced context-aware downstream options, enabling real-time user validation and iterative adaptation (anticipate needs and suggest actions) aligned with the evolving agent strategic objective. This form of interactive and goal-conditioned reasoning exemplifies core agentic capabilities, which surpass the constraints of traditional rule-based approaches (e.g., static dashboard systems) by effectively handling uncertainty, enabling workflow reconfiguration on demand, and supporting user-driven exploration in a dialogic manner.

Thus, CellAtria effectively bridges literature discovery and structured dataset acquisition, establishing a foundation for automated, goal-directed workflows in single-cell research.

### CellAtria Parses Scientific PDFs and Retrieves Sample-Level Datasets

Direct access to structured content on publisher websites can be restricted by technical barriers (e.g., dynamic page rendering, authentication walls) or constrained by licensing terms that limit programmatic scraping - even when institutional access rights are granted.

To address this limitation, the second scenario demonstrates metadata extraction from a locally stored PDF file in place of an article URL. CellAtria mitigates these access constraints by supporting direct parsing of PDF documents, enabling metadata retrieval in settings where web-based extraction is infeasible. We sought to demonstrate this capability by supplying CellAtria with an offline copy of a published study profiling T cell states across tumor, lymph node, and normal tissues in non-small cell lung cancer patients undergoing immune checkpoint blockade^44^. Upon uploading the PDF file, the agent engages in a multi-turn dialogue to extract structured metadata using a built-in document parser, enabling dataset retrieval even when the journal webpage scraping is infeasible (**Supplementary Figs. S12 and S13**). In contrast to the first scenario, where datasets are retrieved at the study level, data acquisition here is carried out at the GEO sample level (GSM) using a task-specific tool, enabling fine-grained retrieval. The agent carries out each stage in response to user instructions conveyed through conversational natural language (**Supplementary Figs. S14 and S15**).

We note that, to support verification and transparency, the real-time log viewer within the CellAtria interface continuously displays each issued prompt along with a status indicator confirming whether the associated tool invocation was successful. This persistent execution trace facilitates user comprehension and troubleshooting during live interactions (**Supplementary Figs. S3-S15**).

Thus, CellAtria enables flexible, fine-grained data acquisition even under restricted access conditions, while preserving full transparency and traceability through real-time execution logs.

### CellAtria Retrieves Pre-Identified Datasets from Public Databases and Manages File Integration

In applied research settings, analysts often begin with pre-identified datasets - discovered through public single-cell portals or cross-study meta-analyses - where the source publication is already known or metadata extraction has been performed independently. In such cases, CellAtria supports direct ingestion of datasets from public repositories using user-supplied download URLs, bypassing upstream literature parsing and metadata extraction steps. This scenario was demonstrated using two curated collections from the CZ Cell by Gene Discover platform^14^: one integrating scRNA-seq data across 13 single-cell studies from 8 tumor types and normal tissues to delineate myeloid-derived cell states^45^, and another compiling scRNA-seq data from 223 patients across 9 cancer types to investigate cancer cell-specific responses to immune checkpoint blockade^46^. Upon receiving the dataset locations, the agent initiates a sequence of interactions with the user, subsequently retrieves the files, integrates them into the working directory, and prepares the necessary configuration files for downstream analysis (**Supplementary Figs. S16-S18**).

This operational mode demonstrates the agent’s ability to coordinate data acquisition and file integration through natural language prompts, enabling flexible invocation of tools at non-linear entry points within the agent’s engineered execution narrative.

### CellAtria Enables Shell-Level Execution and File Navigation within Agent-Guided Workflows

Scientific agentic systems must balance automation with transparency and user oversight, particularly over file system operations, environment context, and task provenance. While language models can coordinate tool execution, these lower-level operations are more effectively managed through dedicated interface components that complement the conversational layer. To address this, CellAtria integrates a set of interactive panels (n=4) that expose critical system-level functionality during live agent sessions (**Supplementary Fig. S19**).

To demonstrate the backend interpretability and user oversight of CellAtria during natural language interaction, we performed a series of targeted tests. Submitting the very simple prompt “Article title?” (implicitly requesting extraction of the publication title from a given URL^43^), we observed the agent’s internal reasoning, tool invocation, and model output displayed step-by-step in the agent backend panel (**Supplementary Fig. S19a**). Despite the minimal input, the agent accurately inferred the intended task, invoked the appropriate tool, and returned only the relevant information - faithfully aligning its output with the user’s request. This live trace offered direct insight into how natural language queries are processed and aligned with the agent’s internal tool logic. To verify the agent’s workspace context and access to relevant data, we then utilized the embedded terminal panel to navigate the project directory. This interaction confirmed the agent’s correct positioning within the expected working path and its access to necessary input files and subdirectories (**Supplementary Fig. S19b**). Complementing the terminal view, the file browser panel allowed us to visually inspect the same directory structure interactively, further reinforcing the consistency between user-issued commands and the agent-managed file system (**Supplementary Fig. S19c**). Finally, to ensure complete task provenance, the chat export utility enabled downloading a structured transcript of the entire session, including prompts and system response traces (**Supplementary Fig. S19d** and **Supplementary Material S1**).

Collectively, these interactive components establish CellAtria as a transparent, traceable, and auditable agentic system. By embedding low-level controls within a high-level dialogue framework, the system balances automation with user-in-the-loop oversight - a critical feature for fostering trust in AI-driven scientific discovery pipelines.

### CellAtria Enables End-to-End Automation of Single-Cell RNA-seq Processing Through CellExpress

Having demonstrated the capabilities and task-specific functionalities of CellAtria, we postulated that by interacting with a fully automated downstream pipeline, CellAtria could further transform literature-based inputs into fully processed single-cell datasets - minimizing the need for hands-on user time and specialized analytic expertise by linking study discovery with standardized downstream analysis through a unified, dialogue-driven workflow.

To this end, we developed CellExpress, a companion computational pipeline that standardizes the processing of scRNA-seq data, transforming raw inputs into biologically interpretable, analysis-ready outputs (**Figure 1d)**. CellExpress builds on previously published and validated methods, ensuring that their application is carried out in a consistent, efficient, and unified workflow with minimal user intervention. A complete methodological description is provided in the Methods section. To facilitate integration with CellAtria, we also developed a suite of agent-triggered tools that support comprehensive pipeline configuration, execution control, and real-time monitoring - enabling the agent to coordinate the entire analytical workflow through natural language interaction (**Supplementary Fig. S2**).

To demonstrate CellExpress orchestration through CellAtria agent, we extended the previously initiated scenario using the peripheral blood scRNA-seq dataset from 2-month-old infants^43^. As with upstream tasks, the pipeline’s configuration, execution, and monitoring were conducted entirely through natural-language dialogue between the user and the agent, culminating in the successful operation of CellExpress pipeline (**Supplementary Figs. S20-S26**). Additionally, to evaluate the full analytical scope of CellExpress, we performed an autonomous pipeline execution with all modules enabled through complete argument specification. In this single-run execution, 18 samples were processed, yielding approximately 71,000 cells after quality control filtering. Stringent thresholds were applied, including a minimum of 750 UMIs per cell, 250 detected genes per cell, expression in at least 3 cells, and a maximum mitochondrial gene content of 15%. Doublets were identified and removed using Scrublet with a score cutoff of 0.25, and batch effects were corrected to account for sample-specific variation. Dimensionality reduction was conducted using both UMAP and t-SNE embeddings, followed by graph-based clustering. Automated cell type annotation showed high concordance between tissue-agnostic and tissue-specific models, highlighting the pipeline’s robustness in immune profiling contexts. Finally, cluster-level marker gene identification was performed to support downstream biological interpretation (**Supplementary Material S2**). All relevant artifacts - including the execution summary, sample metadata, and workflow configuration - were stored in a structured file to ensure auditability and reproducibility (**Supplementary Material S3**).

Thus, CellExpress addresses persistent challenges in single-cell transcriptomic analysis by delivering a fully integrated, end-to-end workflow that ensures transparency and reproducibility through the use of rigorously benchmarked, field-standard components.

### CellAtria Enables Full-Lifecycle Document-to-Analysis Execution Through Fully Autonomous Toolchain Orchestration

The hallmark of robust agentic systems lies in their ability to autonomously synthesize complex operational sequences from a single, high-level command - abstracting procedural complexity while preserving fidelity to the intended outcome. Such generalizable and autonomous orchestration reflects a carefully designed, context-aware system prompt and an explicit, unambiguous definition of tool input/output (I/O) behaviors - elements that collectively underpin the system’s capacity for full-scope task execution.

To evaluate the extent of autonomous task coordination in CellAtria, we tested its ability to execute a complete document-to-analysis workflow using a single instruction, thereby eliminating the need for iterative user-agent interaction. In this scenario, the agent was provided with the primary article URL reporting longitudinal scRNA-seq data from 2-month-old infants^43^, with the objective of executing the CellExpress pipeline using the associated single-cell data. In response - and without further user input - CellAtria autonomously carried out the full underlying predesigned agentic workflow, leading to the successful execution of the CellExpress pipeline (**Supplementary Fig. S27**). In particular, CellAtria performed several key steps autonomously, including parsing the primary article, extracting structured metadata, retrieving the associated dataset, configuring the CellExpress pipeline with context-aware parameters, and dispatching the execution - all without manual intervention (**Supplementary Fig. S28**). Notably, this autonomous run - excluding the CellExpress pipeline’s runtime - completed all preparatory steps in under 10 minutes. This performance stands in stark contrast to the approximately 15 cumulative hours of manual effort typically required by a bioinformatics analyst for equivalent tasks, as per our internal benchmarks.

Hence, this assessment demonstrates CellAtria’s capability for strategic problem-solving by translating high-level objectives into a coherent sequence of tool invocations - marking its shift from prompt-driven response to an AI agent capable of autonomous workflow orchestration with minimal intervention and time-efficient performance.

## Discussion

This study introduces CellAtria, an agentic system that enables dialogue-driven, document-to-analysis automation in single-cell research. The system integrates a natural language interface with modular computational toolchains - including task-specific utilities that trigger the CellExpress pipeline - to form a unified semantic layer that orchestrates data triage and analysis. Its composable architecture supports non-linear, context-aware interaction, allowing users to engage flexibly at different stages of data ingestion and preparation. This design enables researchers to process more studies with less effort, effectively improving their analytical capacity without sacrificing reproducibility or protocol adherence.

CellAtria’s strategy of orchestrating a pre-vetted, modular toolchain (including CellExpress pipeline) leverages the LLM’s strengths in intent interpretation and task delegation. This approach guarantees that all executed analytical steps (including complex single-cell data analysis) adhere to established best practices, are fully transparent, and maintain the high level of reliability and auditability essential for robust scientific discovery, all while leveraging the full orchestration potential of LLMs.

Agentic systems operate according to an underlying execution narrative - a structured sequence of modular actions that defines how tasks are interpreted and fulfilled. While this narrative is inherently flexible and permits user-initiated entry at arbitrary points in the workflow, it remains anchored in a coherent logic that guides the agent’s behavior and goals. This design departs from traditional rule-based automation by allowing unconstrained user interaction while still aligning those inputs with predefined tools and execution pathways. In the case of CellAtria, this narrative integrates a four-stage operational sequence: (1) metadata extraction, (2) dataset acquisition, (3) file organization, and (4) downstream analysis execution. Each module can be invoked independently; however, this study focuses on the optimized execution path, which reflects the intended canonical workflow.

CellAtria can be characterized as a goal-oriented agent: it interprets user intent, structures task sequences, and autonomously executes predefined workflows to accomplish specific analytical objectives. Distinct from reactive systems limited to scripted responses, CellAtria orchestrates multi-step operations through a unified execution layer that maps high-level prompts to appropriate computational tools. It also incorporates context-aware behavior and exhibits within-session adaptivity, dynamically adjusting its actions in response to evolving user input. Furthermore, its ability to deconstruct abstract objectives into sequenced tool invocations reflects domain-specific autonomous behavior within a well-defined operational scope.

Agentic AI frameworks establish a principled division of responsibilities: domain experts engage with high-level analytical tasks through natural language interfaces, while system developers ensure the integrity and robustness of the underlying toolchains. The core reasoning engine of these agents - LLMs - while effective at semantically interpreting dynamic user prompts and aligning them with task objectives, remains inherently prone to hallucination^33, 34^, often producing linguistically coherent yet semantically or factually incorrect outputs. Although such vulnerabilities can be mitigated in constrained execution environments through carefully engineered system prompts and tightly scoped tool schemas, they may still persist when the agent encounters ambiguous instructions or tasks beyond its operational scope. To safeguard analytical reliability and ensure interpretability, maintaining a human-in-the-loop paradigm - wherein a context-aware investigator actively evaluates, verifies, and contextualizes agent responses - remains indispensable.

CellAtria metadata extraction capabilities are optimized for scientific articles that conform to structured narrative conventions - such as standardized sectioning and consistent biomedical terminology. While the underlying language model possesses broad generalization capacity, the system may face inconsistency when applied to unstructured or idiosyncratic content, including informal supplementary notes lacking hierarchical organization or consistent labeling, or documents containing irregular phrasing such as nonstandard abbreviations, ad hoc formatting, or domain-specific shorthand.

A key design consideration for agentic systems is the LLM’s lack of direct internet access - a common constraint adopted to ensure security and maintain control over external interactions. Consequently, the system depends on deterministic, tool-mediated mechanisms for content retrieval and cannot independently browse or query online databases in real time. This reliance highlights the critical importance of standardized data management practices^47^: discrepancies between manuscript-reported metadata and repository-level annotations can hinder the reliability of agentic execution. We therefore advocate for closer harmonization between narrative metadata and structured repository schemas to better support scalable agentic applications.

The modular design of CellAtria, combined with its graph-based execution architecture, provides a foundation for extensibility and seamless integration of additional computational tools. This framework supports future expansion beyond the system’s current operational scope; the present implementation maintains a balance, leveraging human-in-the-loop control to preserve interpretability and task-level adaptability, even as forthcoming iterations may incorporate more autonomous execution capabilities - driven by predefined workflow presets - particularly in high-confidence settings, characterized by well-defined data inputs and established processing logic, where minimal user prompting suffices.

In conclusion, CellAtria demonstrates how domain-informed agentic AI systems can operationalize scientific research processes in a computationally skill-agnostic manner, thereby accelerating discovery and supporting the transition toward next-generation, AI-integrated research ecosystems.

## Methods

### CellAtria Interface and Architectural Framework

The CellAtria interface integrates seven principal components that provide fine-grained control over task execution and collectively facilitate rich agentic interaction (**Supplementary Fig. S1**): (1) Persistent Chatbot Window: Manages user–agent communication and maintains conversational continuity through an internal hidden state, supporting coherent, multi-turn exchanges. (2) User Input Panel: Accepts textual prompts and facilitates document uploads, with all inputs jointly processed through a unified execution handler. (3) Real-time Log Viewer: Displays user-agent transaction status. (4) Agent Backend Panel: Provides a live, step-by-step view of the agent’s internal reasoning, tool invocation sequence, and backend responses, directly supporting transparency and debugging. (5) Embedded Terminal Panel: Enables direct shell interaction within the agent’s runtime environment, facilitating robust system-level control without exiting the interface. (6) Interactive File Browser: Allows users to navigate project directories and inspect file contents within the active workspace, complementing terminal operations. (7) Chat Export Utility: Generates structured conversation transcripts in a machine-readable format to support downstream auditing and reproducibility. These diverse interface elements are orchestrated atop a LangGraph-based^48^ backbone that encodes the agent’s execution flow logic as a directed graph of modular, state-aware functions-ensuring robust and interpretable coordination across heterogeneous toolchains and complex computational workflows.

### CellAtria Modular Toolchain for Task Execution

To enable flexible and robust task execution, we developed a comprehensive suite of 32 interoperable tools that encapsulate core agent functionalities across four principal operational domains (**Supplementary Fig. S2**): (1) Metadata Parsing and Semantic Structuring: Handles the extraction and organized representation of relevant information from diverse sources. (2) Programmatic Data Retrieval and Hierarchical Organization: Manages the automated acquisition of datasets and their structured arrangement within the workspace. (3) Standardized File Handling and Pre-processing: Ensures consistent management and preparation of data files for downstream analysis. (4) Automated Workflow Configuration and Execution Orchestration: Facilitates the dynamic setup and control of complex computational analysis pipelines. Each tool is precisely implemented as an atomic function with rigorously defined input/output (I/O) behavior, a design choice that fundamentally enables the agent to compose dynamic and reliable task sequences in response to user prompts. These tools are inherently embedded as graph nodes within CellAtria’s architectural backbone and accessed through natural-language interfaces, thereby facilitating intelligent, context-aware orchestration. This modular architecture not only accommodates user-initiated entry at arbitrary points within the workflow but also establishes a crucial foundation for tool reuse, adaptation, and the scalable extension of execution flows.

### Standardized Single-Cell Data Analysis via CellExpress

A cornerstone of CellAtria’s full-scale capabilities is the co-developed CellExpress, a standardized single-cell analysis pipeline - fully automated and engineered to deliver robust scRNA-seq analysis, from raw count matrices through comprehensive processing and report generation (**Figure 1d**). Designed to lower bioinformatics barriers, CellExpress implements a comprehensive set of state-of-the-art, Scanpy-based^49^ processing stages, including: (1) quality control; performed globally or per sample, (2) data transformation; encompassing normalization, highly variable gene selection, and scaling, (3) dimensionality reduction; utilizing UMAP^50^ and t-SNE^51^, (4) graph-based clustering^52^, and (5) marker gene identification. Additional tools are seamlessly integrated to support advanced analysis tasks, such as doublet detection (Scrublet^53^), batch correction (Harmony^54^), and automated cell type annotation using both tissue-agnostic (SCimilarity^55^) and tissue-specific (CellTypist^56^) models. All analytical steps are executed sequentially under centralized control, with parameters fully configurable via a comprehensive input schema.

The pipeline natively supports a wide range of standard input formats including 10X Genomics Cell Ranger^57^ outputs (provided as the trio of a UMI count matrix, a cell barcode list, and a gene annotation file), as well as HDF5 (h5) and AnnData objects.

Upon execution, CellExpress processes a designated set of scRNA-seq samples and generates a comprehensive analytical package comprising four components: (1) a finalized, fully annotated AnnData object, directly suitable for downstream analysis; (2) a structured, publication-ready HTML report that captures a complete snapshot of the entire workflow, including key parameters, quality control metrics, dimensionality reduction, clustering results, and cell type annotations, presented through dynamics tables and graphical visualizations; (3) a complete export of all configuration settings in a machine-readable, standardized format to ensure auditability and full reproducibility; and (4) a quality control-filtered AnnData object generated for reuse in alternative or customized workflows.

Designed for flexible deployment, CellExpress operates as a fully standalone pipeline for comprehensive scRNA-seq data analysis. It can be orchestrated either through an agentic system - as incorporated into the CellAtria framework - or via direct command-line execution. Furthermore, its Pythonic foundation directly addresses scalability constraints commonly associated with R-based pipelines^58^, enabling more efficient handling of large single-cell datasets.

### CellAtria Containerized Environment and CellExpress Execution

The CellAtria runtime environment is fully containerized using Docker, enabling consistent and reproducible deployment across diverse computational infrastructures. This encapsulation strategy ensures environmental parity by isolating workflows from system-specific variability and mitigating software dependency conflicts, thereby facilitating seamless portability across local, cloud, and high-performance computing environments.

To preserve agent responsiveness during potentially long-running computations, CellAtria executes the CellExpress pipeline in a detached mode. Upon receiving a complete execution schema, the agent delegates the task to a background subprocess, decoupling it from the interactive session. Standard output and error streams are redirected to persistent log files for downstream inspection, and a unique process identifier is recorded to support real-time status tracking and diagnostics.

### Large Language Model Provider and Computational Environment

The CellAtria environment is designed as a multi-tenant system, capable of interfacing with a variety of large language model (LLM) providers, include Azure OpenAI, OpenAI, Anthropic, Google, as well as local deployments.

For the benchmarking experiments presented in this study, the LLM backend was provisioned through the Azure OpenAI service, specifically utilizing a managed deployment of gpt-4o accessed via Azure API endpoints. All associated downstream agent interactions, metadata parsing, and pipeline execution tasks were comprehensively carried out on an AWS EC2 r6i.32xlarge instance configured with 1,024 GiB of memory.

## Data availability

No new data were generated in this study. The datasets used for demonstration purposes in the use case scenarios are publicly available. Data for the first scenario are accessible through the NIH Gene Expression Omnibus (GEO) under accession number GSE204716, and for the second scenario under accession number GSE185206. The processed objects used in the third scenario are available via the CZ Cell by Gene Discover platform at the following links: https://cellxgene.cziscience.com/collections/3f7c572c-cd73-4b51-a313-207c7f20f188 and https://cellxgene.cziscience.com/collections/61e422dd-c9cd-460e-9b91-72d9517348ef.

## Code availability

The source code, documentation, and all materials required to execute the agent are publicly available on the GitHub at https://github.com/astrazeneca/cellatria.

## Acknowledgments

We would like to extend our gratitude to the developers and maintainers of the various software packages and tools that were instrumental to this research.

## Author contributions

N.N. and R.A.: Conceptualization. N.N.: Methodology, software development, and original manuscript drafting. N.N., R.A., and V.S.: Manuscript review, editing, and content refinement. V.S.: Supervision and strategic direction.

