## Supplementary material for "An Agentic AI Framework for Ingestion and Standardization of Single-Cell RNA-seq Data Analysis": Supplementary_Figures.pdf

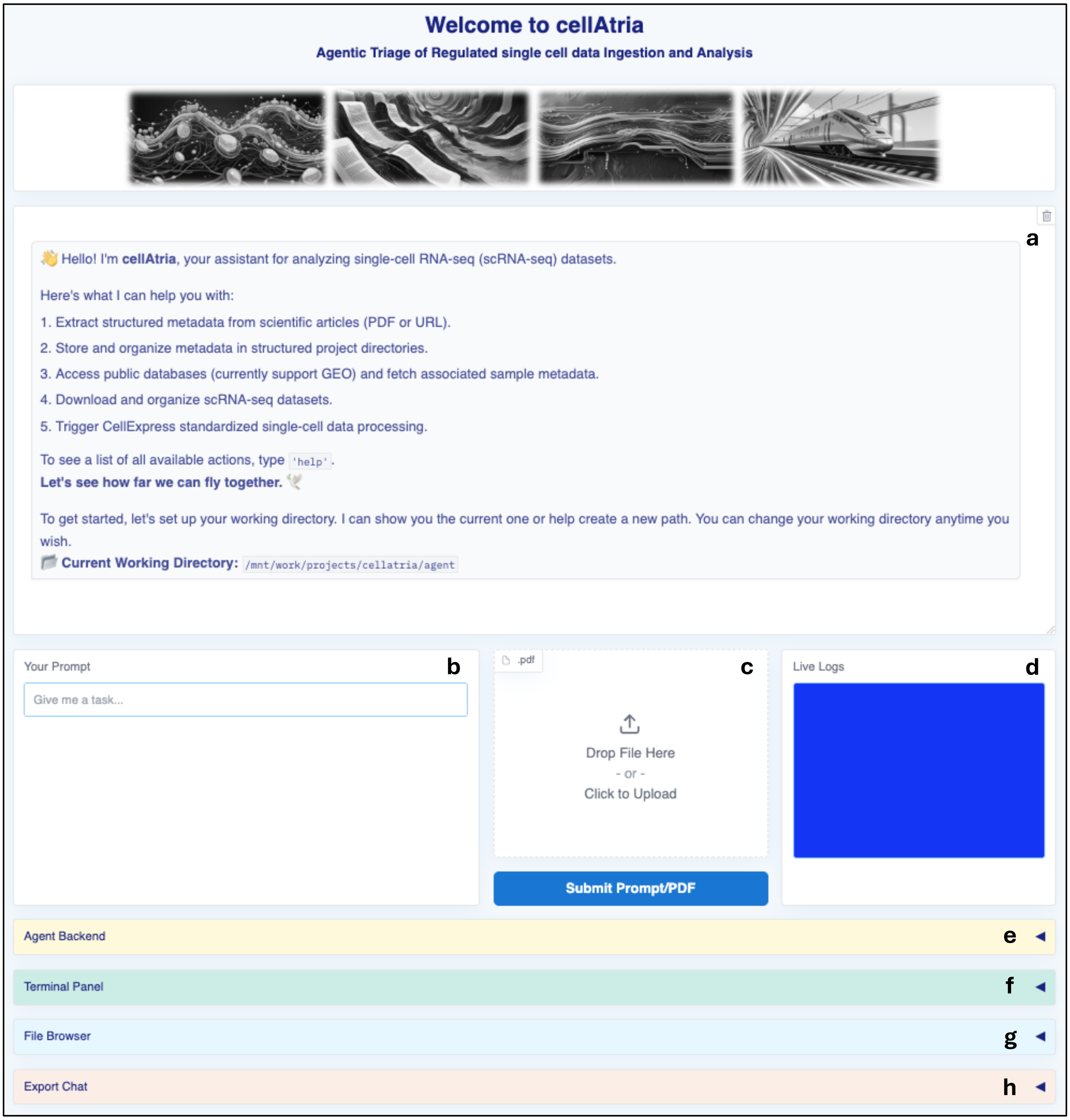


**Supplementary Figure S1. User interface of the CellAtria agentic system. a)** Persistent chatbot window that mediates user-agent interaction. **b)** Input panel for free-text prompts. **c)** PDF upload widget enabling ingestion of documents and submit button that triggers the unified execution handler, dispatching tasks based on input context. **d)** Live log viewer providing real-time telemetry on agent execution status. **e)** The agent backend panel displays a live, step-by-step stream of the agent's internal reasoning, tool calls, and backend responses. **f)** Embedded terminal interface for issuing UNIX-style shell commands within the agent context. **g)** Interactive file browser for navigating directory structures, inspecting files. **h)** Export utility for downloading conversation transcripts in machine-readable format. Note: the system-level panels (terminal, file browser, and chat export) are designed as collapsible accordion sections.

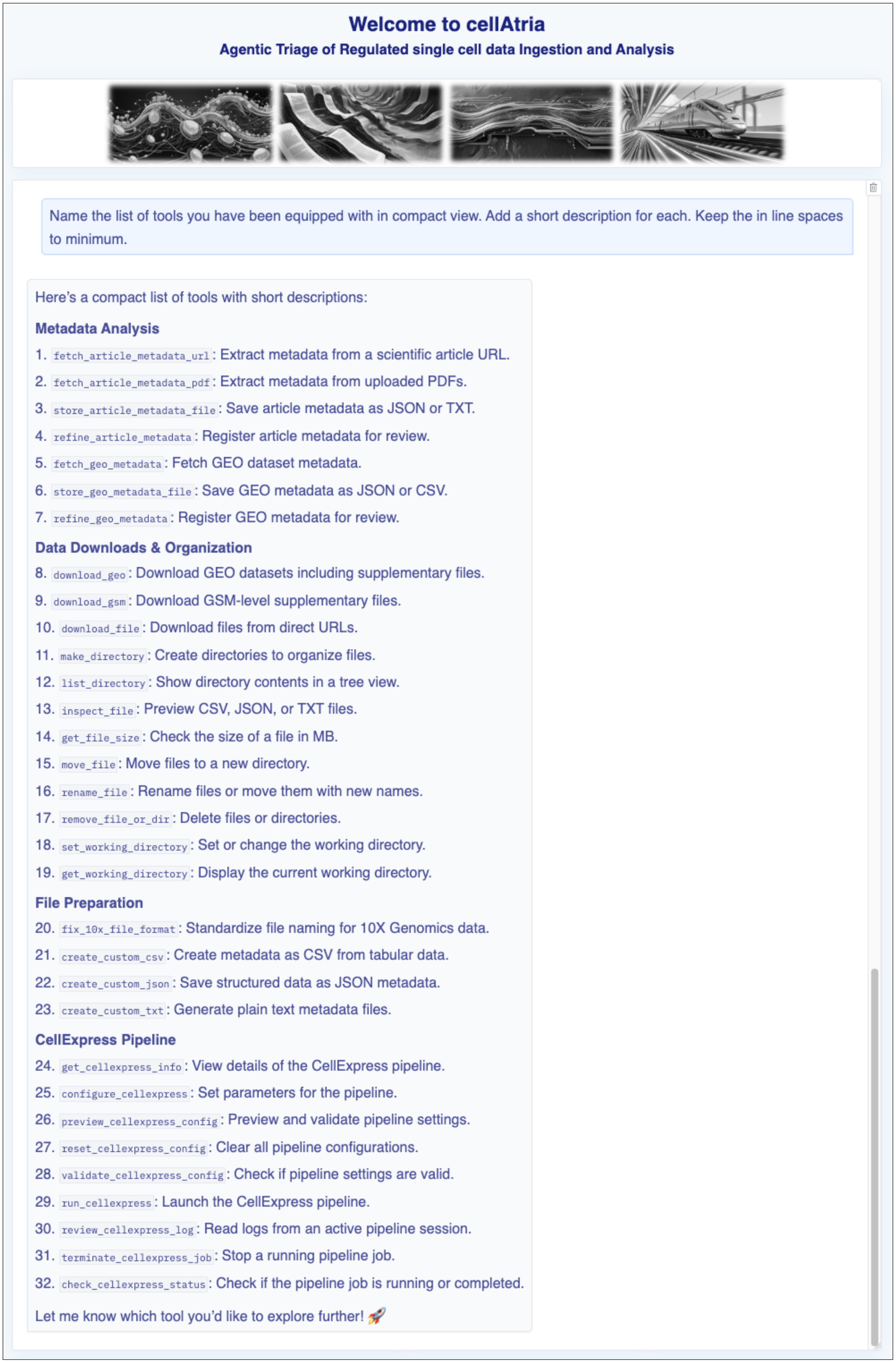


**Supplementary Figure S2. Overview of the CellAtria toolchain in response to a prompt querying its functional capabilities.** The agent enumerates its internal toolkit, which includes modules for article metadata extraction, dataset access, file system management, metadata structuring, and CellExpress configuration. Each tool is exposed as a callable function with defined scope and I/O specifications. Note: the system-level panels (backend, terminal, file browser, and chat export) have been cropped for improved visibility.

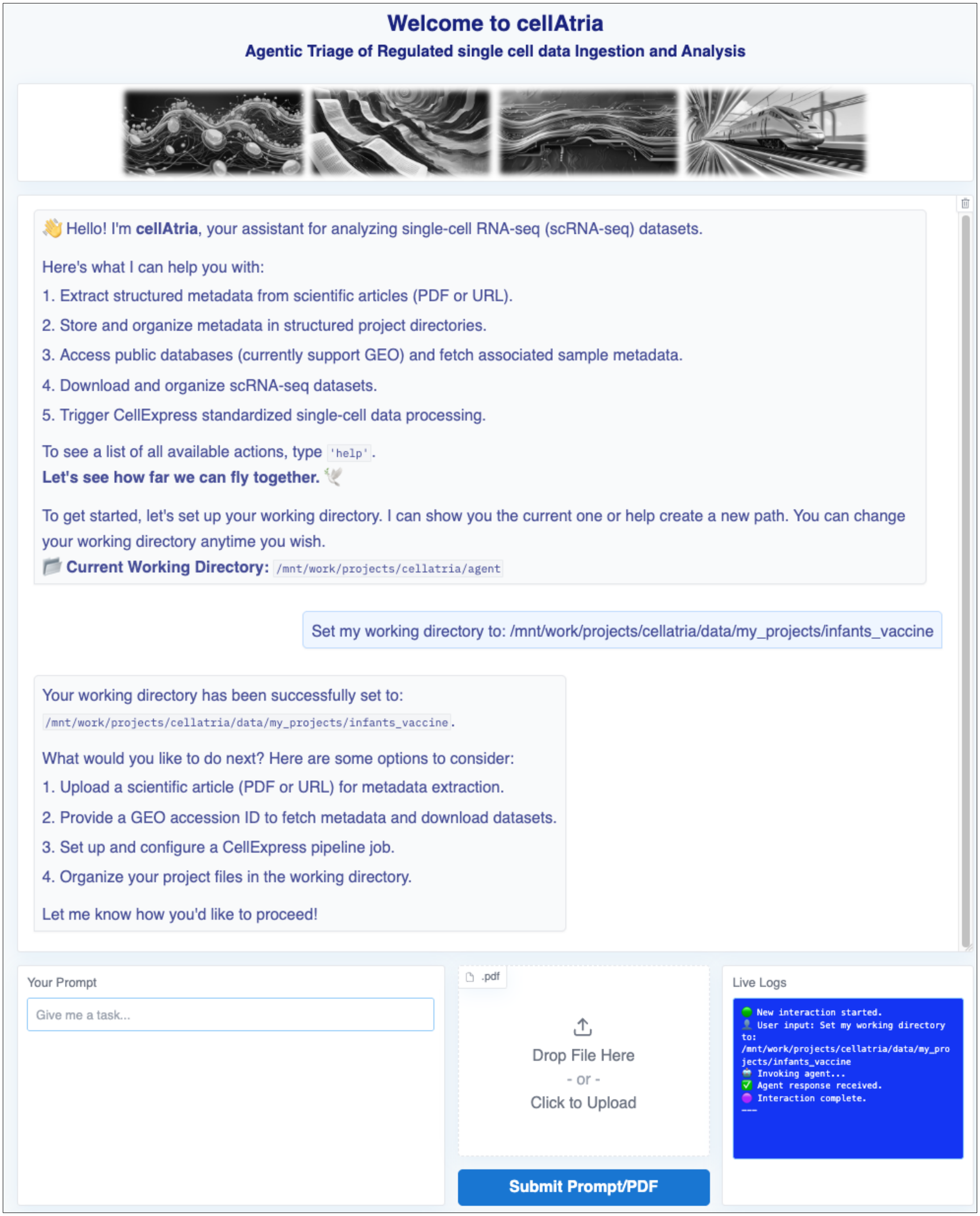


**Supplementary Figure S3. Interactive initialization of a working directory within the CellAtria agentic interface.** In response to a user prompt, CellAtria updates the working directory and offers context-aware follow-up actions. The integrated live log viewer provides real-time feedback on execution status and confirms successful completion. Note: the system-level panels (backend, terminal, file browser, and chat export) have been cropped for improved visibility.

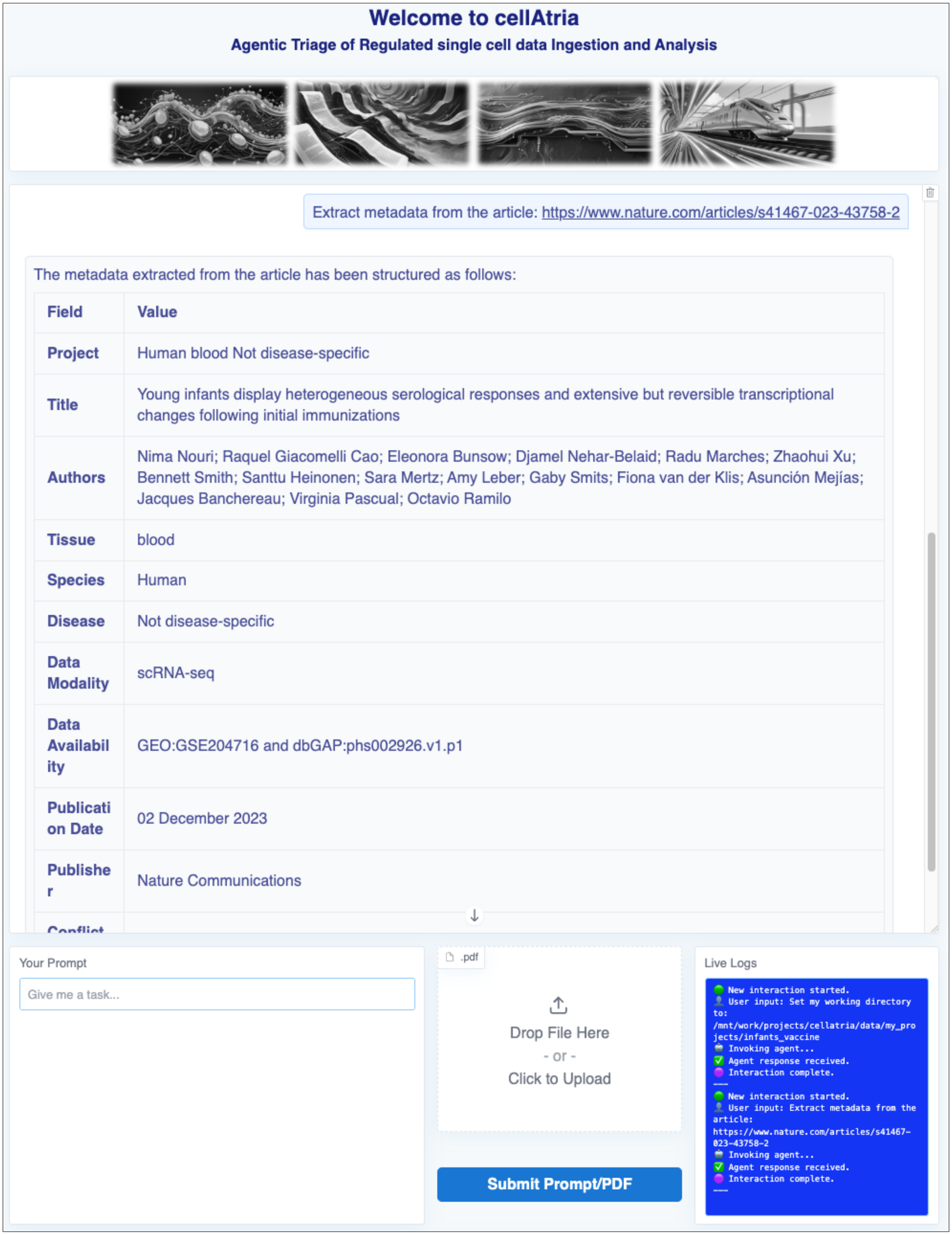


**Supplementary Figure S4. Metadata extraction from a DOI-based URL using the CellAtria agentic interface.** In response to a user-provided article URL, the agent retrieves and parses the full-text document, converting key study descriptors into a structured tabular format. The integrated live log viewer provides real-time feedback on execution status and confirms successful completion. Note: the system-level panels (backend, terminal, file browser, and chat export) have been cropped for improved visibility.

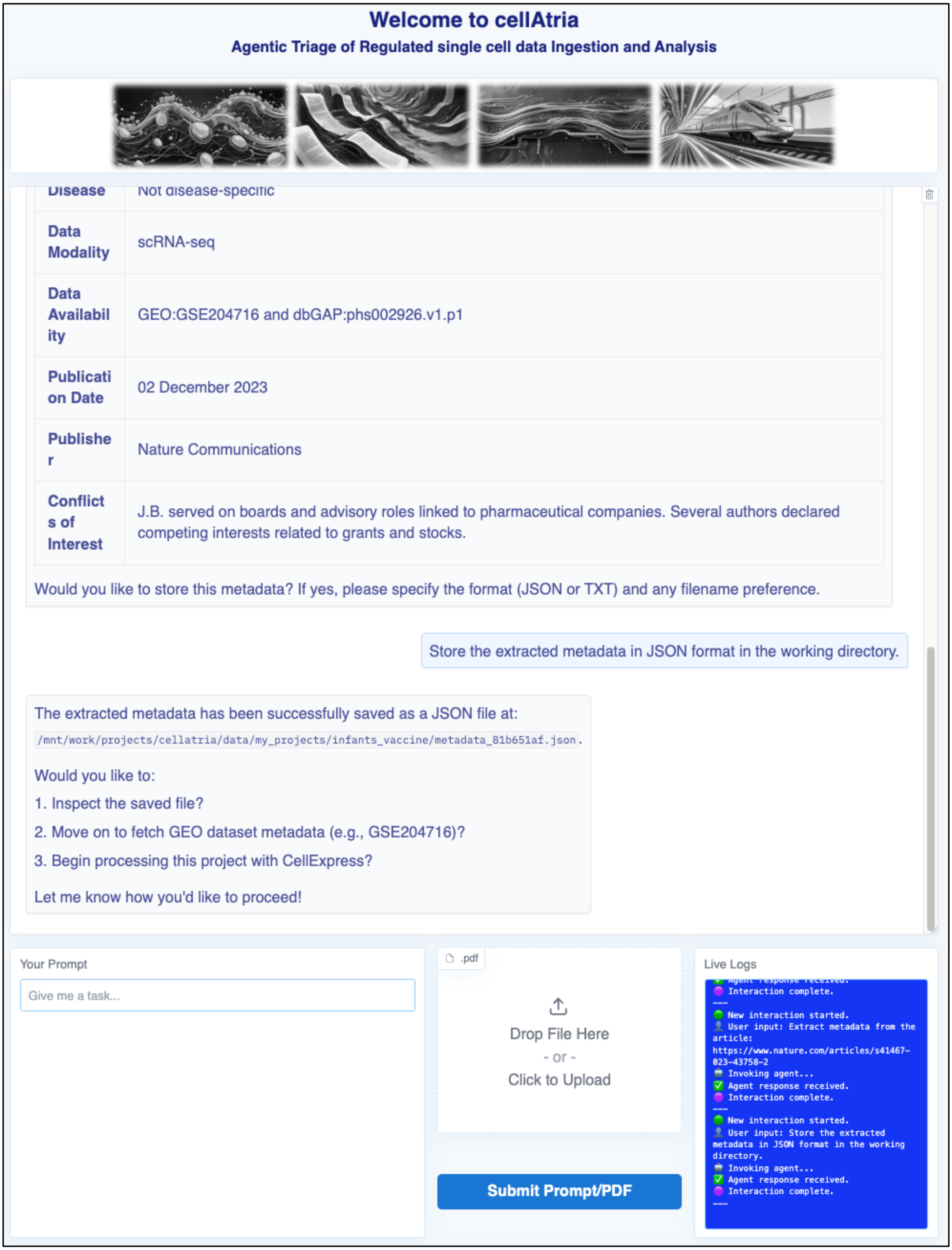


**Supplementary Figure S5. Structured metadata export to file following article parsing in CellAtria.** Upon user request, the metadata is saved as a JSON file in the active working directory. The interface confirms the file path and proposes context-aware next steps. The integrated live log viewer provides real-time feedback on execution status and confirms successful completion. Note: the system-level panels (backend, terminal, file browser, and chat export) have been cropped for improved visibility.

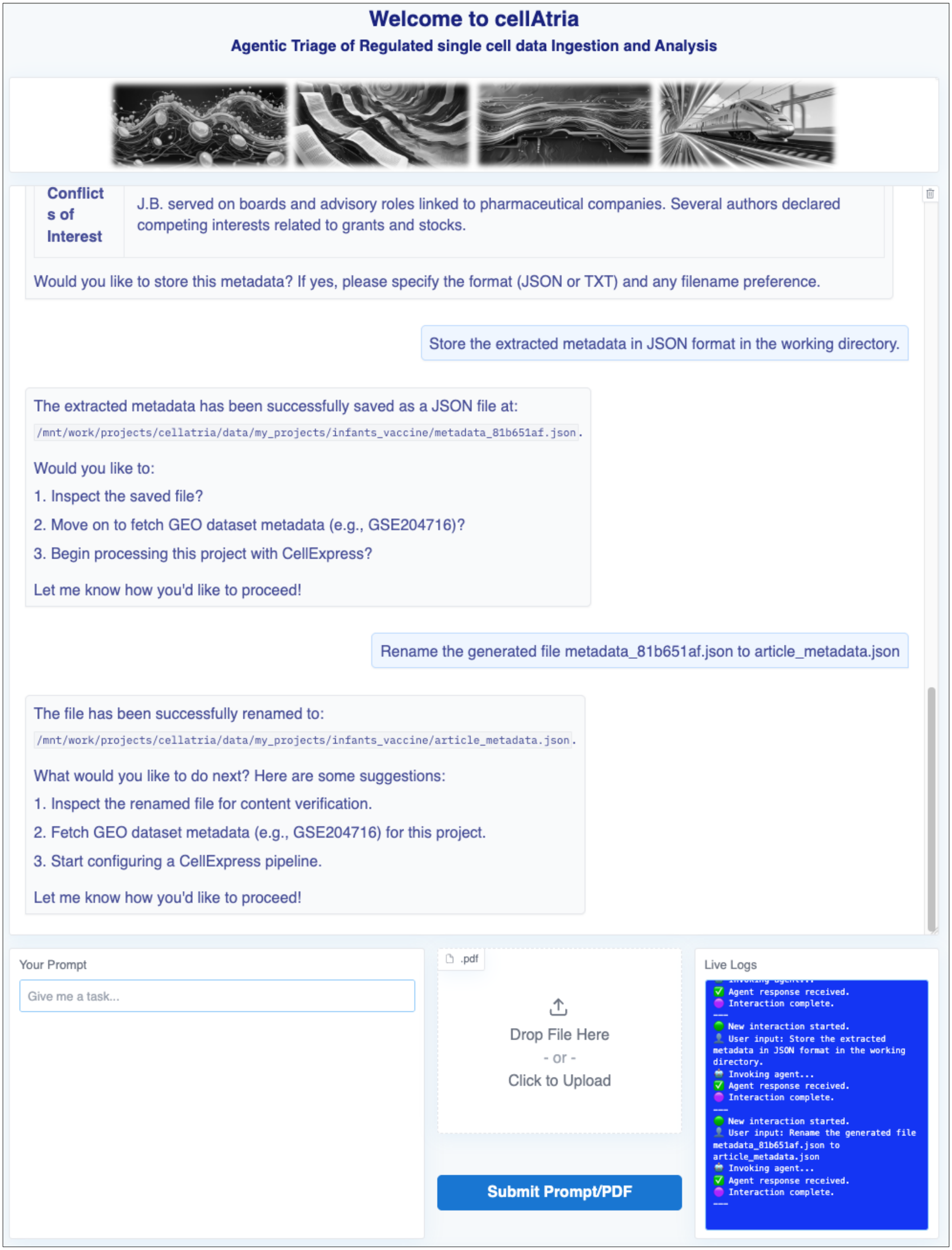


**Supplementary Figure S6. File management automation for structured metadata workflows.** CellAtria autonomously renames the output file based on a user-specified name within the active working directory. The task includes confirmation prompts to validate the action. The integrated live log viewer provides real-time feedback on execution status and confirms successful completion. Note: the system-level panels (backend, terminal, file browser, and chat export) have been cropped for improved visibility.

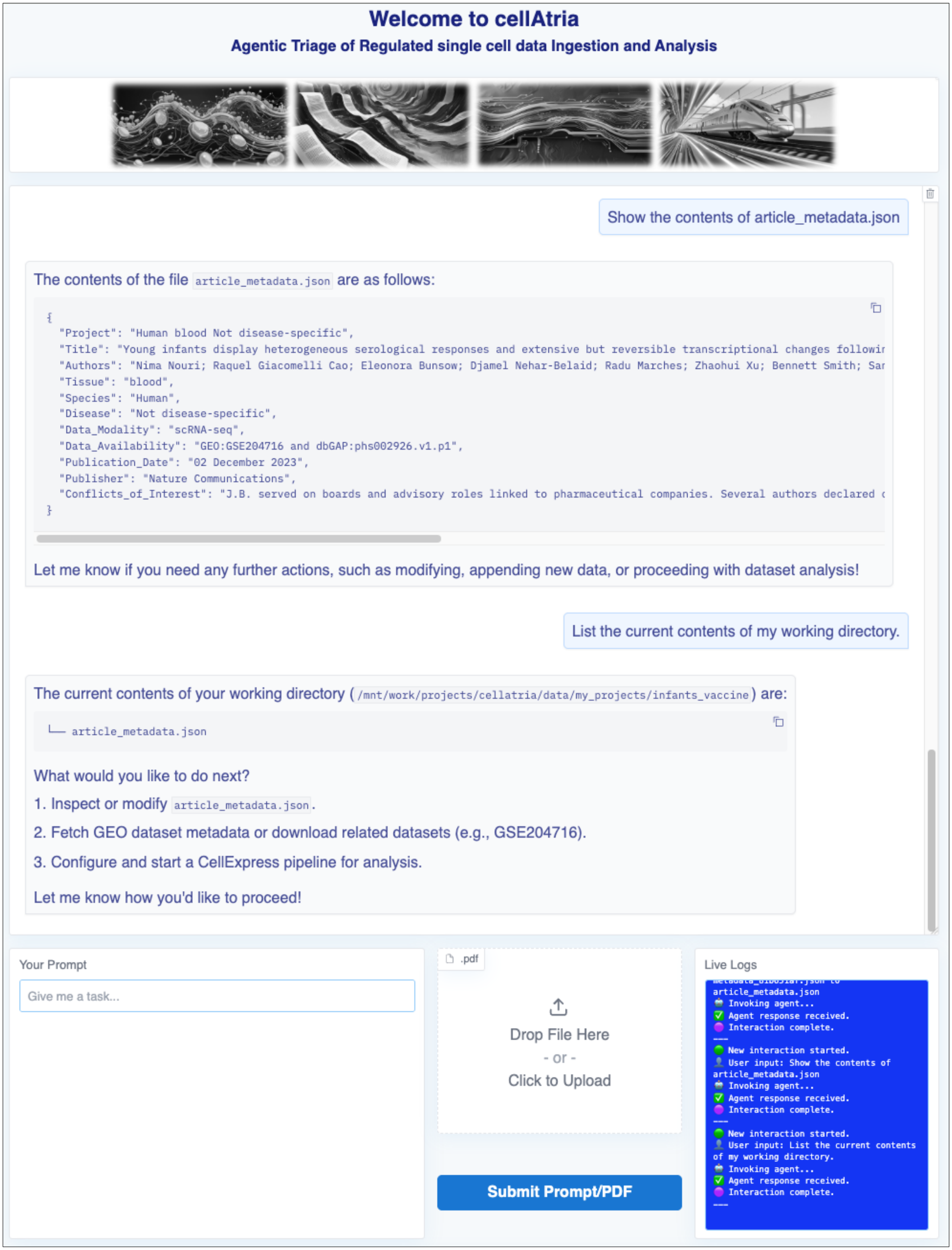


**Supplementary Figure S7. Structured file inspection and metadata verification.** Upon user request, CellAtria displays the contents of the stored metadata file and inspects the current working directory. The integrated live log viewer provides real-time feedback on execution status and confirms successful completion. Note: the system-level panels (backend, terminal, file browser, and chat export) have been cropped for improved visibility.

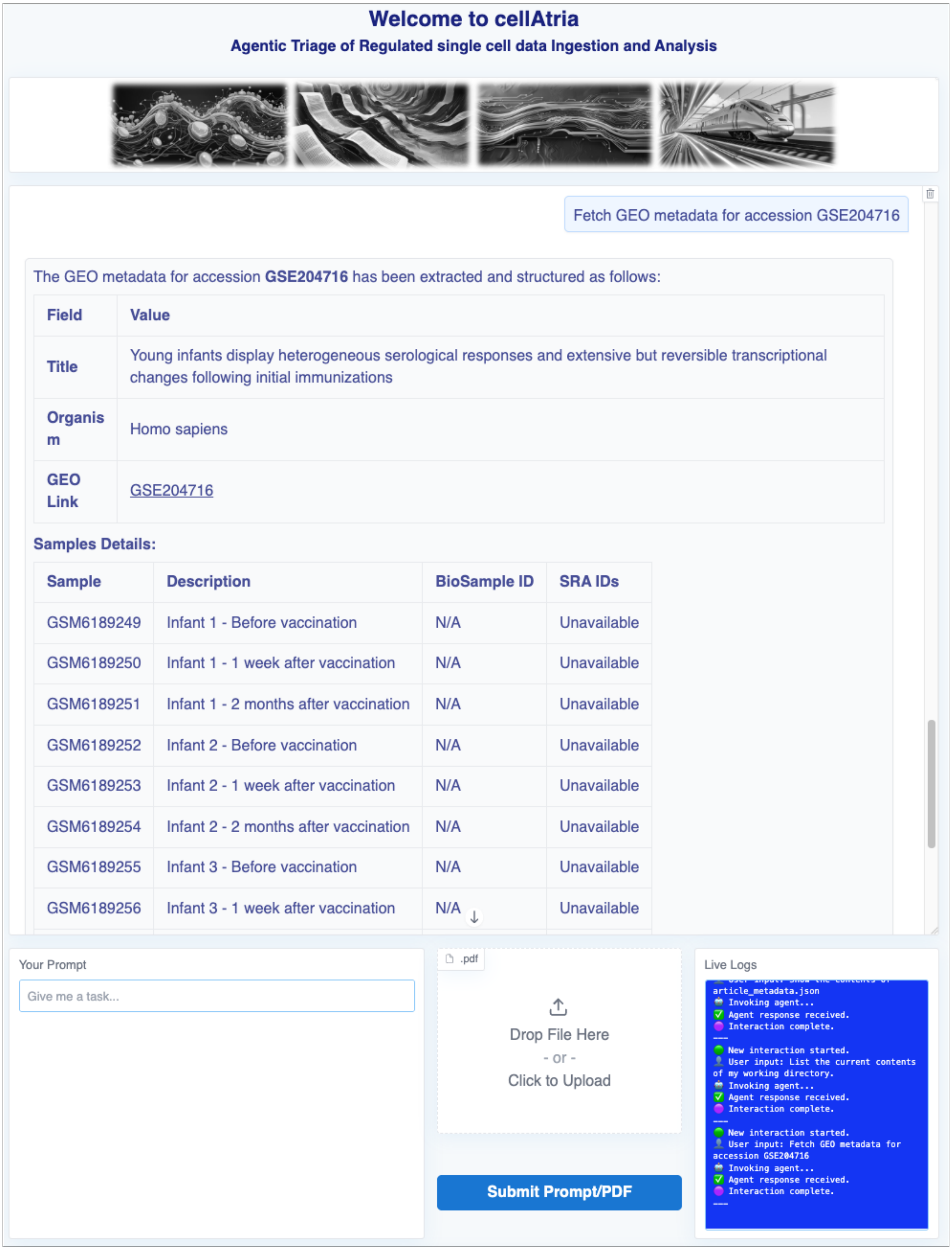


**Supplementary Figure S8. Agent-driven GEO metadata extraction and sample summary.** CellAtria retrieves and parses GEO study metadata, displaying structured sample-level information including descriptions and accession statuses. The integrated live log viewer provides real-time feedback on execution status and confirms successful completion. Note: the system-level panels (backend, terminal, file browser, and chat export) have been cropped for improved visibility.

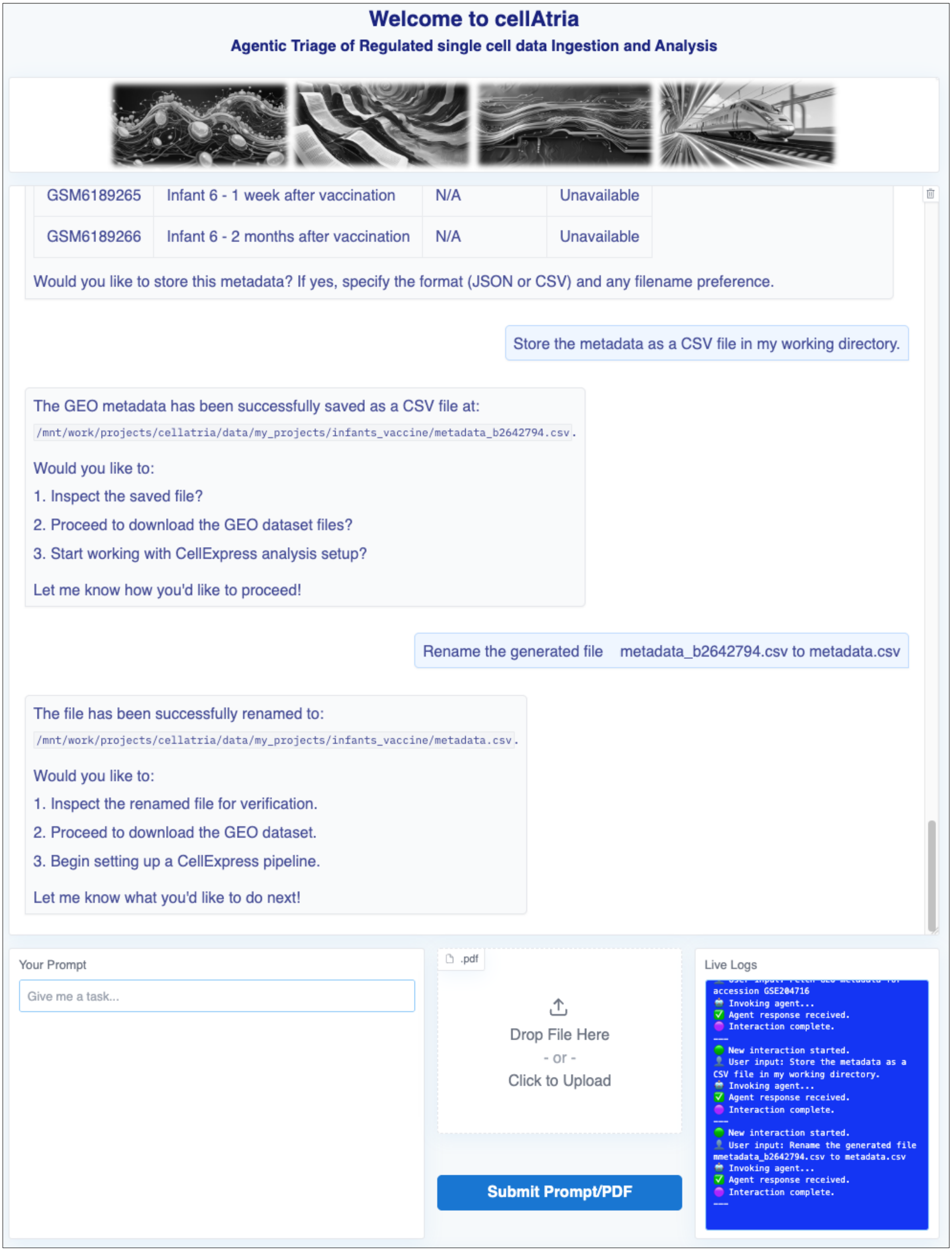


**Supplementary Figure S9. Structured storage and renaming of extracted GEO metadata.** CellAtria stores the parsed metadata as a CSV file in the working directory and facilitates user-guided file renaming. The interface confirms path updates and suggests next actions. The integrated live log viewer provides real-time feedback on execution status and confirms successful completion. Note: the system-level panels (backend, terminal, file browser, and chat export) have been cropped for improved visibility.

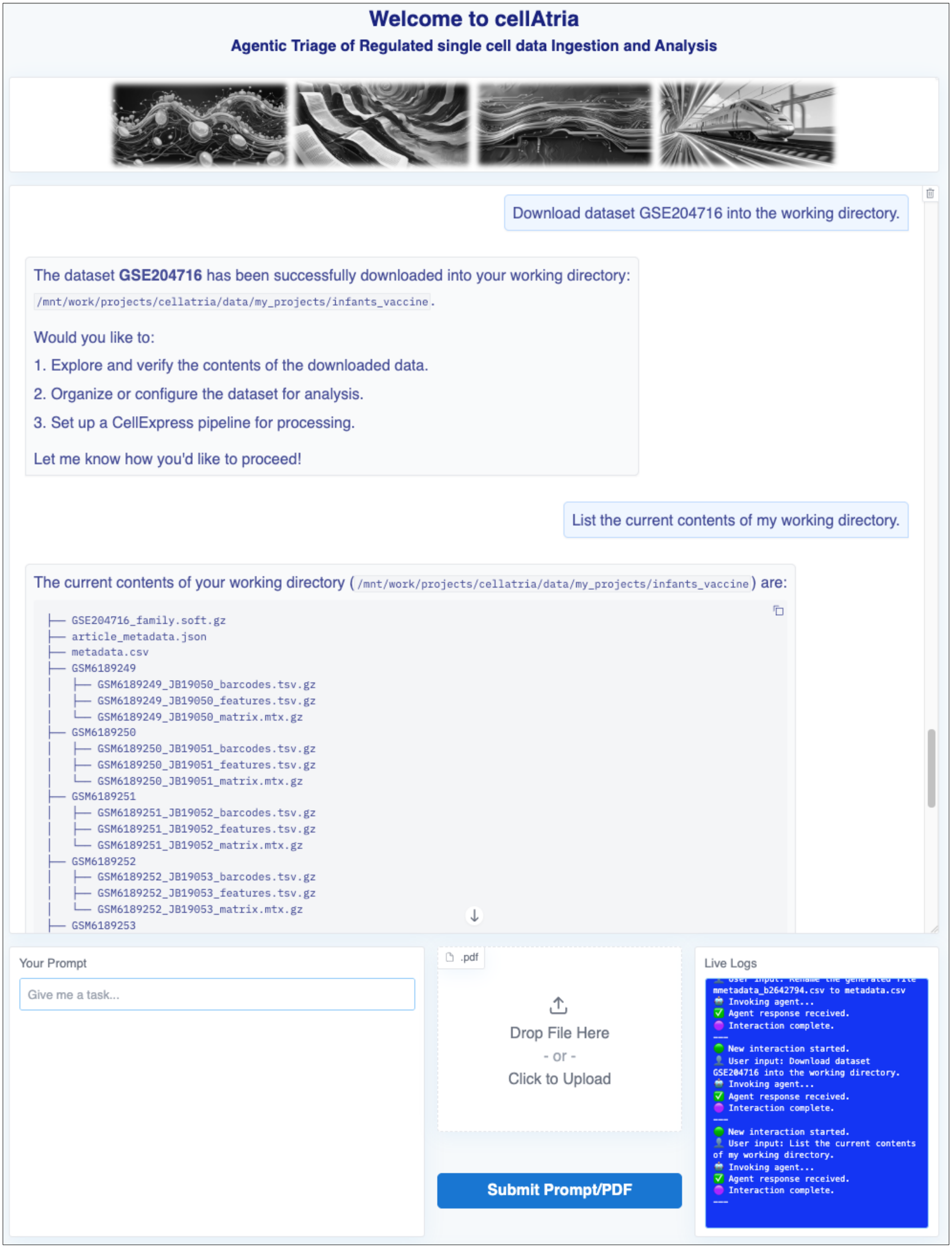


**Supplementary Figure S10. Automated dataset retrieval.** Following a user request, CellAtria downloads all supplementary files for the specified GSE dataset and stores them in the working directory. The agent then provides a complete file listing upon user request. The integrated live log viewer provides real-time feedback on execution status and confirms successful completion. Note: the system-level panels (backend, terminal, file browser, and chat export) have been cropped for improved visibility.

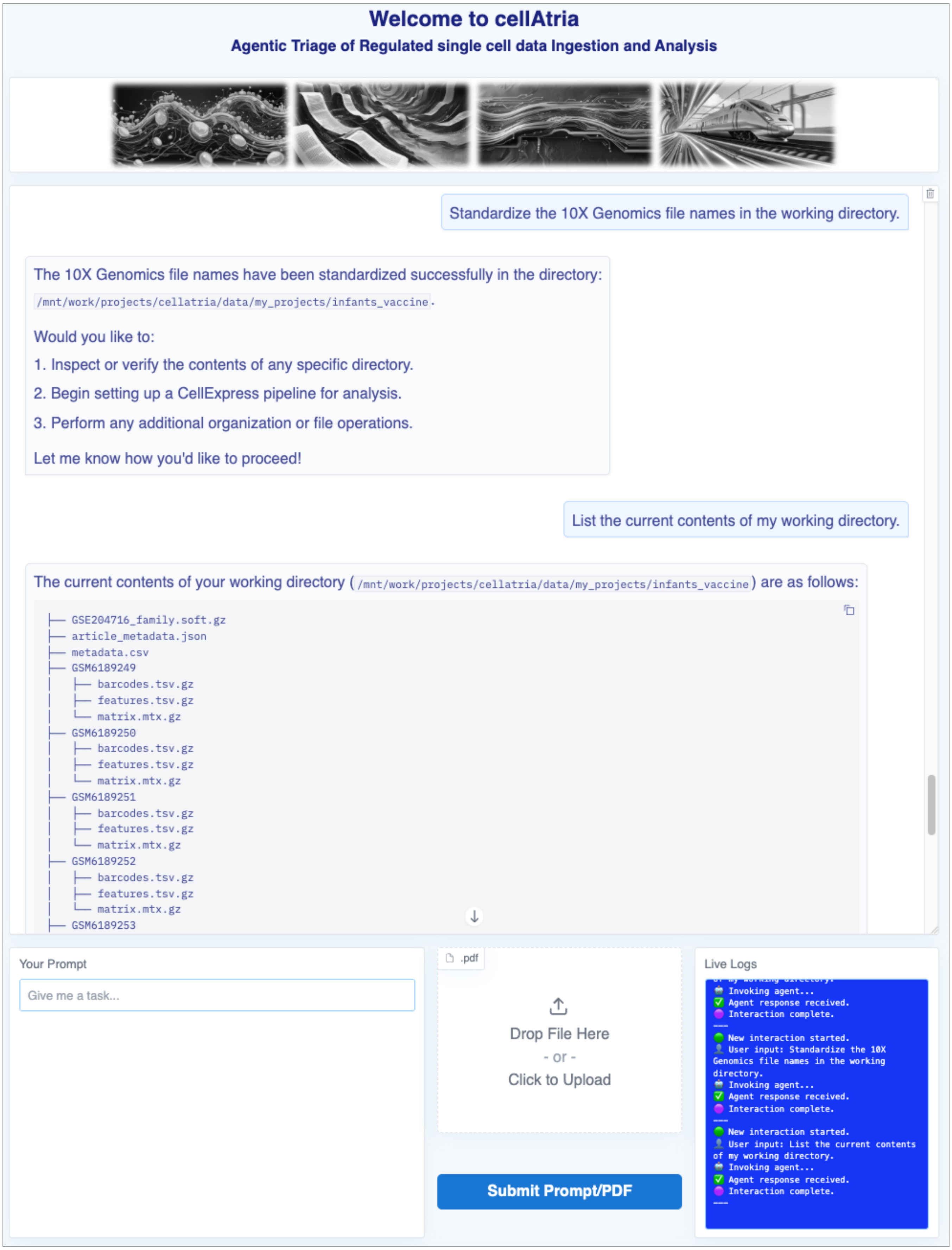


**Supplementary Figure S11. Standardized file renaming for downstream compatibility.** CellAtria programmatically renames all files to conform with 10X Genomics conventions, ensuring compatibility with downstream pipelines. A directory listing is presented upon request. The integrated live log viewer provides real-time feedback on execution status and confirms successful completion. Note: the system-level panels (backend, terminal, file browser, and chat export) have been cropped for improved visibility.

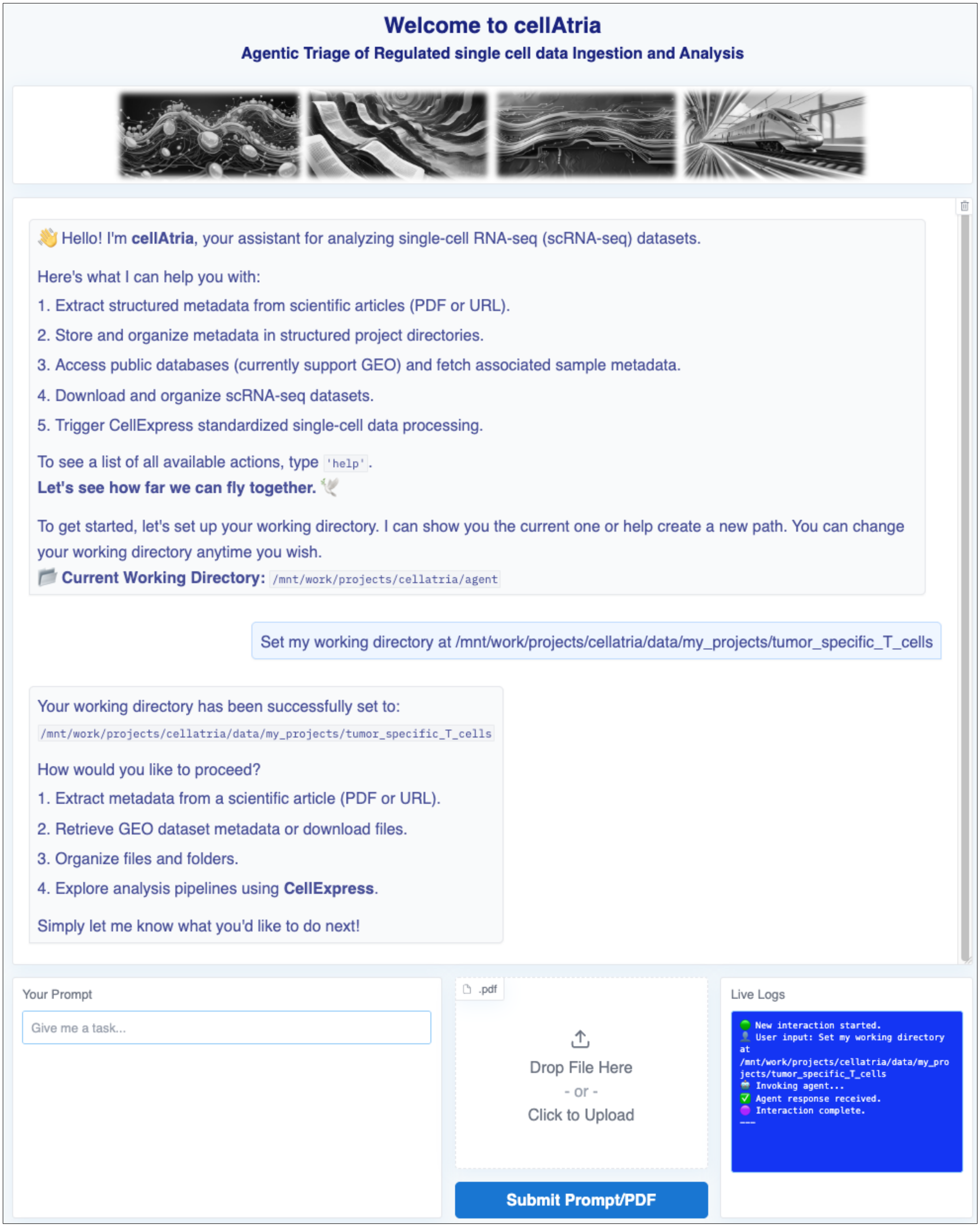


**Supplementary Figure S12. Interactive initialization of a working directory within the CellAtria agentic interface.** In response to a user prompt, CellAtria updates the working directory and offers context-aware follow-up actions. The integrated live log viewer provides real-time feedback on execution status and confirms successful completion. Note: the system-level panels (backend, terminal, file browser, and chat export) have been cropped for improved visibility.

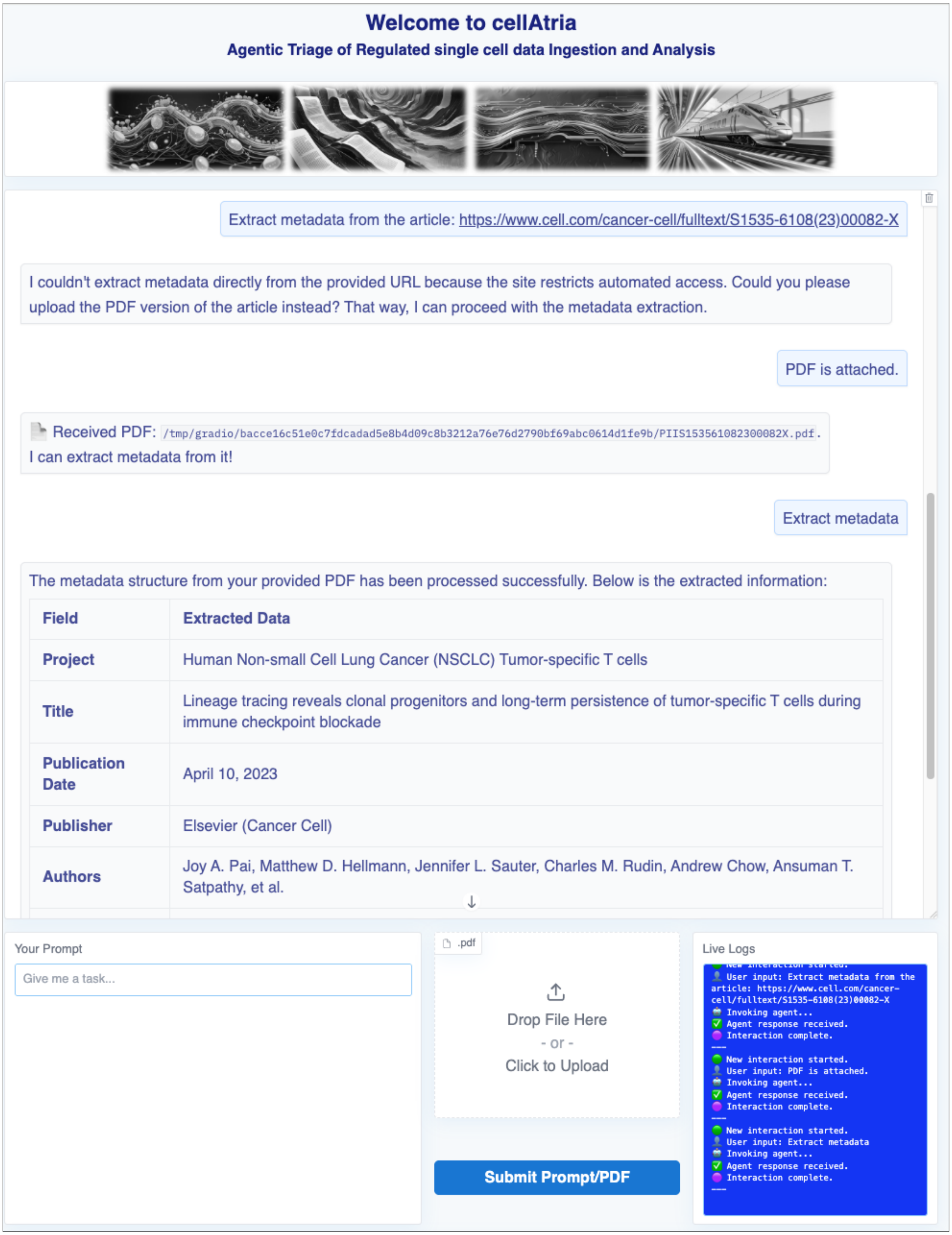


**Supplementary Figure S13. PDF-driven metadata extraction workflow in CellAtria.** When automated access to an article URL is restricted, the user uploads a locally stored PDF. The agent parses the document and extracts structured metadata which is then presented in a tabulated format. The integrated live log viewer provides real-time feedback on execution status and confirms successful completion. Note: the system-level panels (backend, terminal, file browser, and chat export) have been cropped for improved visibility.

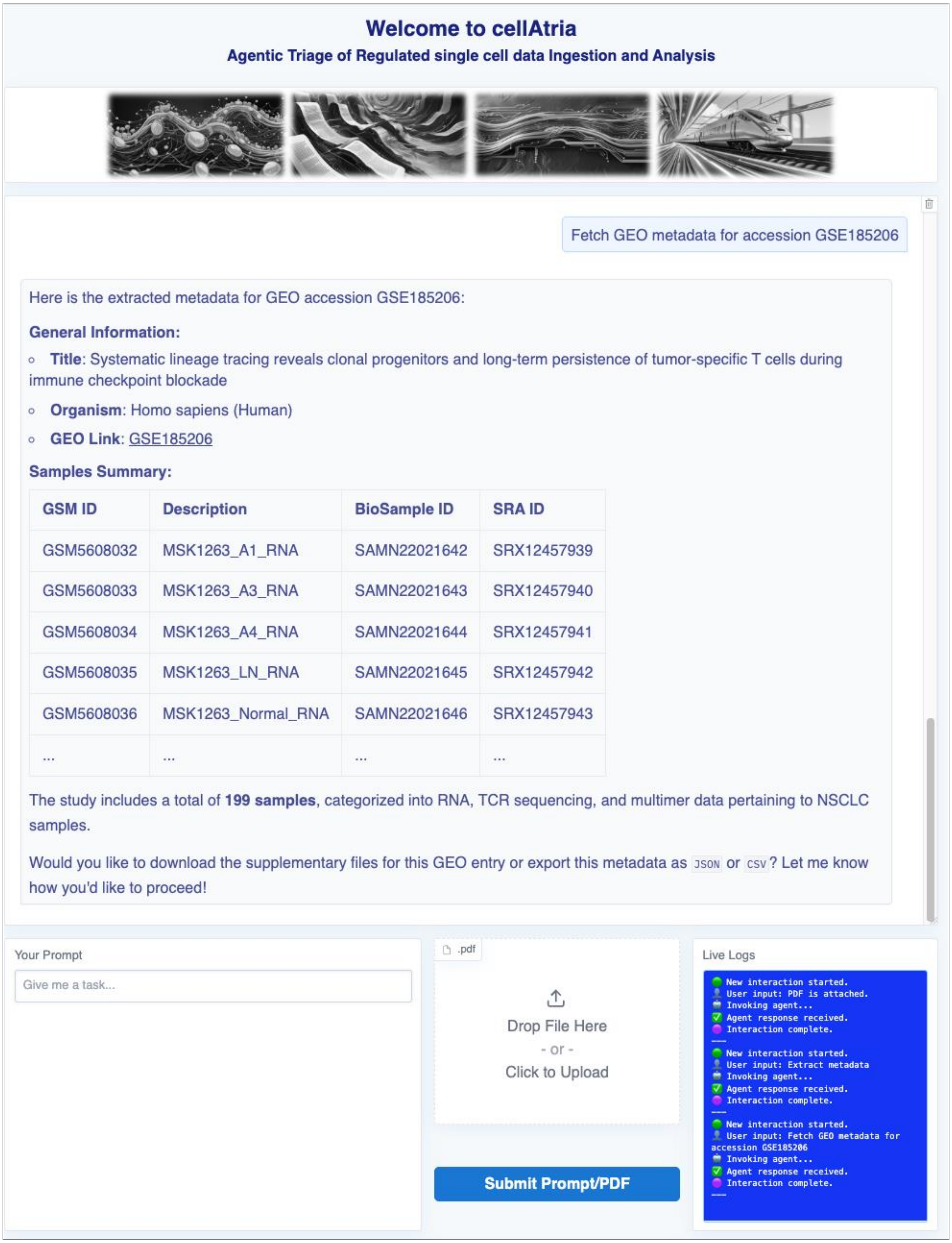


**Supplementary Figure S14. GEO metadata extraction in CellAtria following PDF-based article parsing.** Upon user request, the agent retrieves and displays study metadata associated with the referenced GEO accession. The interface summarizes sample-level information and prompts the user to proceed with file download or metadata export. The integrated live log viewer provides real-time feedback on execution status and confirms successful completion. Note: the system-level panels (backend, terminal, file browser, and chat export) have been cropped for improved visibility.

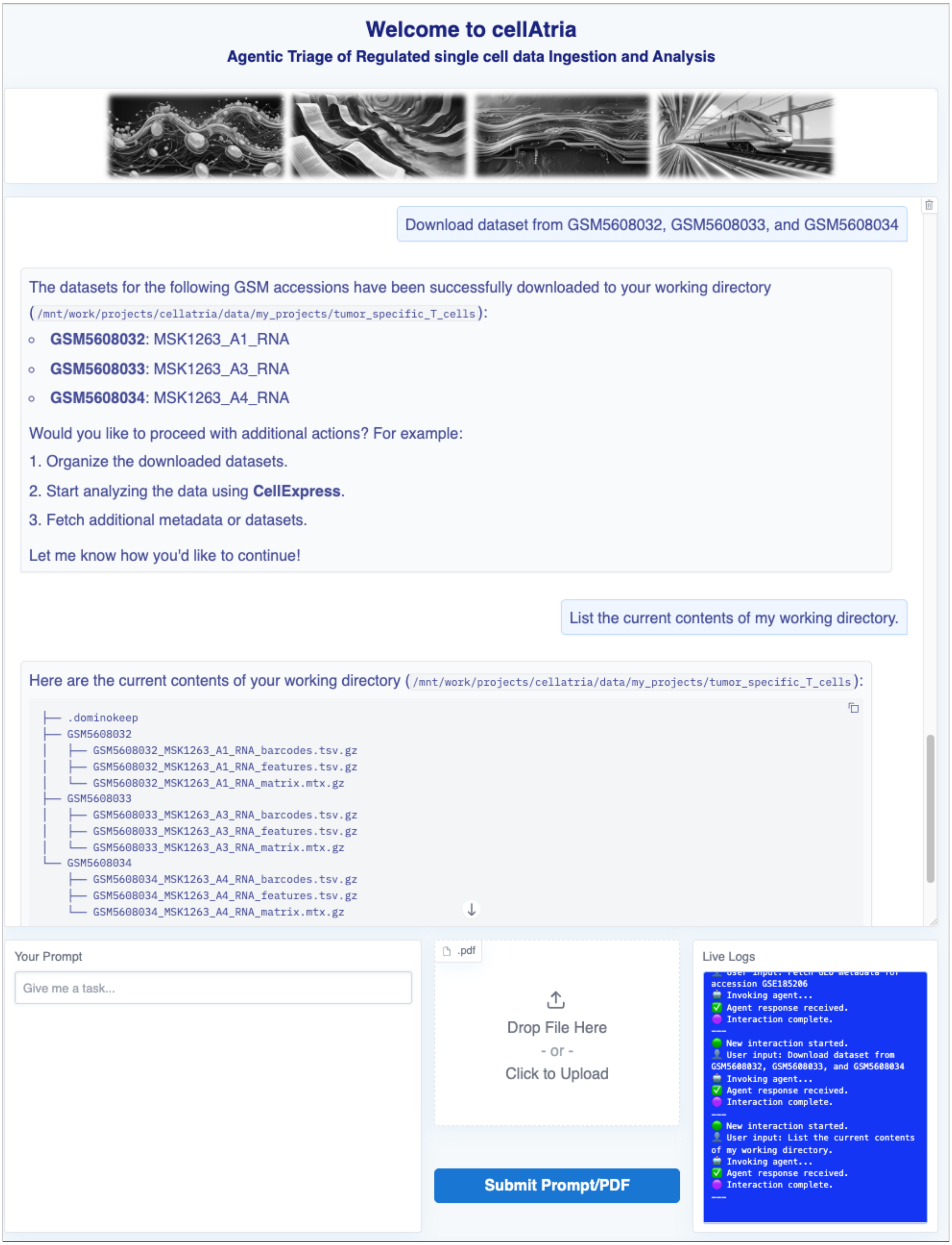


**Supplementary Figure S15. GSM-based sample-level dataset acquisition in CellAtria.** Upon user request, the agent downloads the specified GSM accessions and confirms successful retrieval. The interface presents a structured view of the file contents for each sample. The integrated live log viewer provides real-time feedback on execution status and confirms successful completion. Note: the system-level panels (backend, terminal, file browser, and chat export) have been cropped for improved visibility.

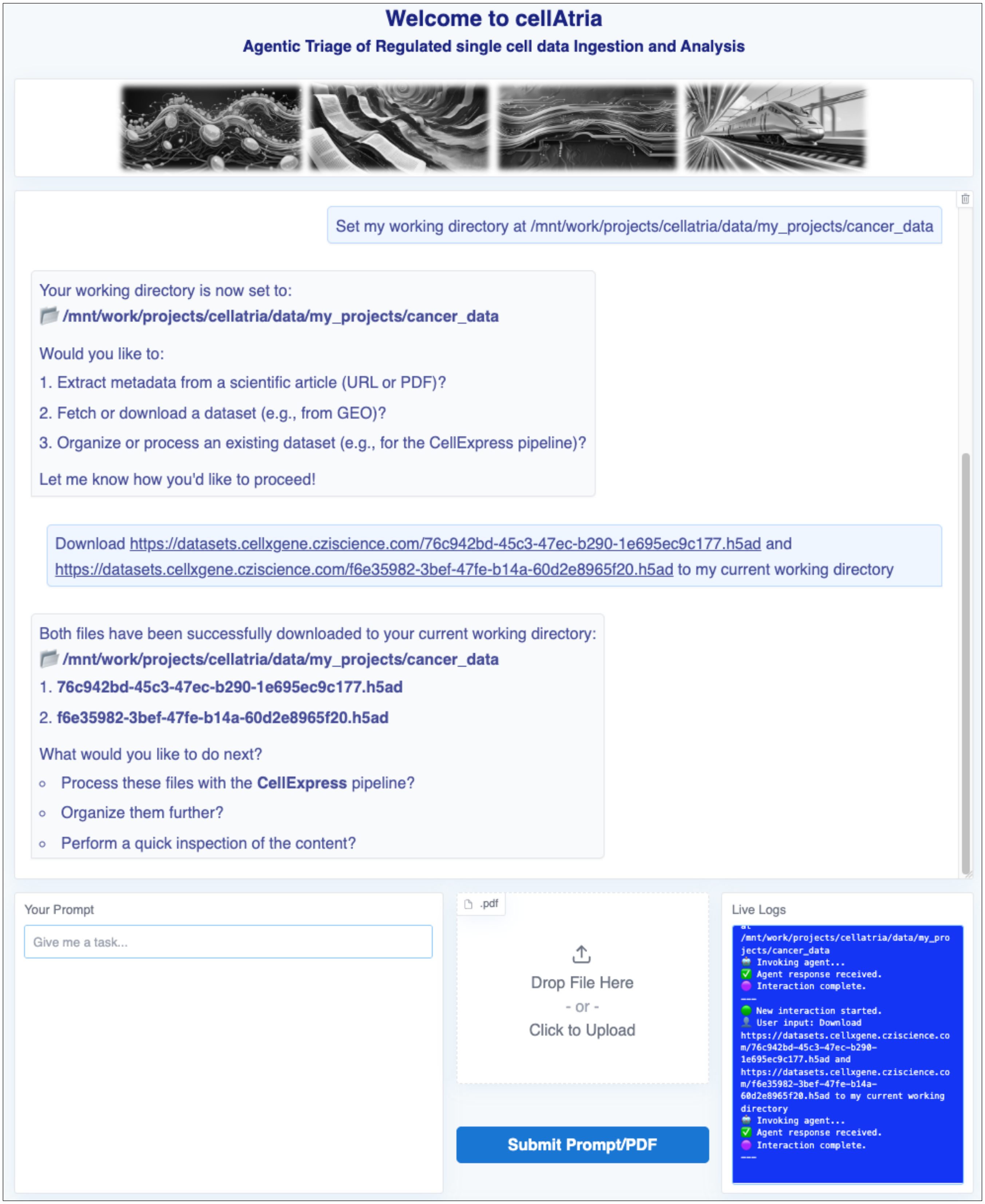


**Supplementary Figure S16. Direct download and workspace integration of preprocessed datasets.** Upon receiving a user prompt, the agent sets the working directory and confirms the change with contextual follow-up options. When provided with direct URLs, the agent retrieves and stores the datasets in the specified location. The integrated live log viewer provides real-time feedback on execution status and confirms successful completion. Note: the system-level panels (backend, terminal, file browser, and chat export) have been cropped for improved visibility.

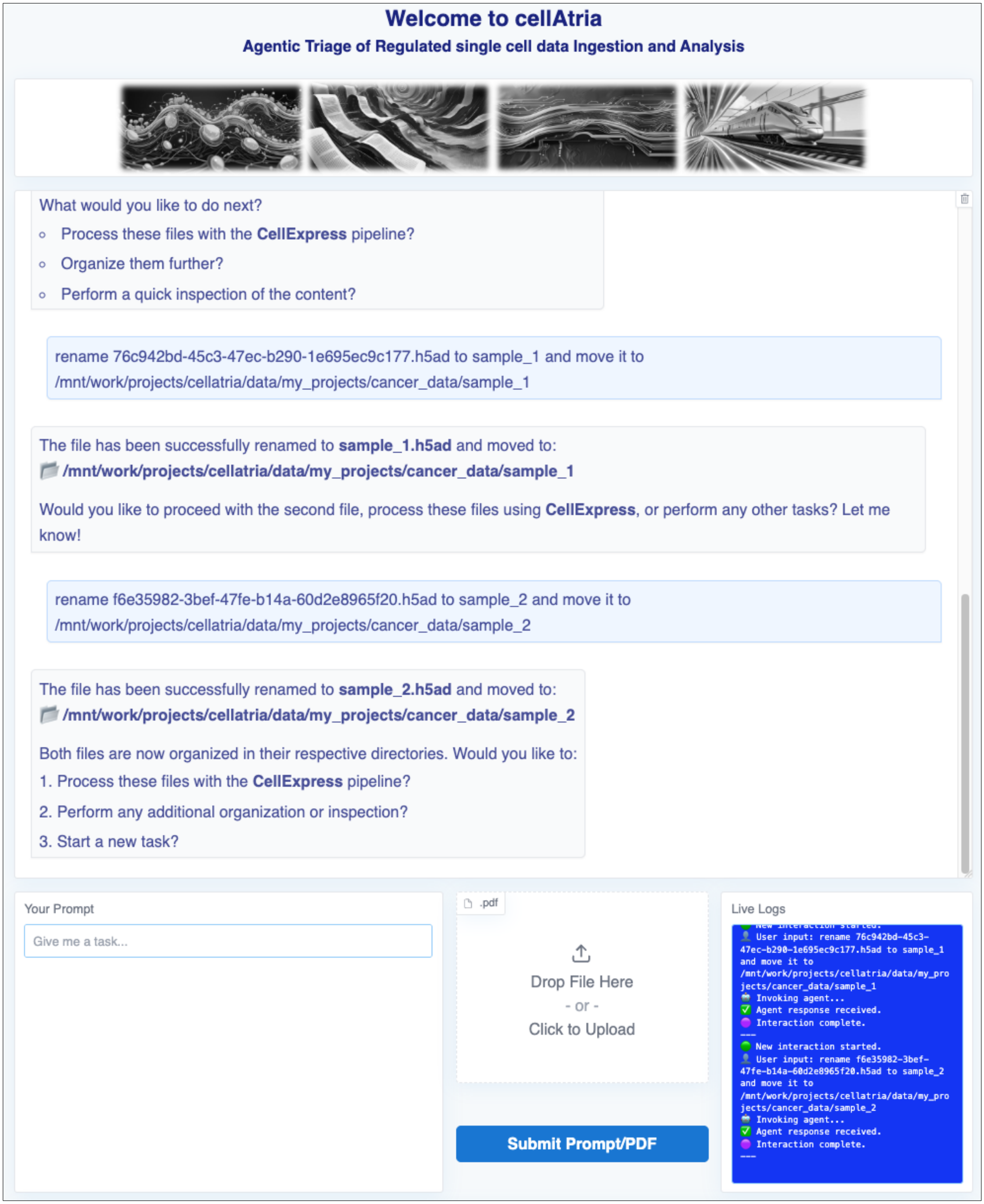


**Supplementary Figure S17. Dataset renaming and organization within the working directory.** Upon user request, the agent renames the datasets and organizes them into sample-specific subdirectories. The integrated live log viewer provides real-time feedback on execution status and confirms successful completion. Note: the system-level panels (backend, terminal, file browser, and chat export) have been cropped for improved visibility.

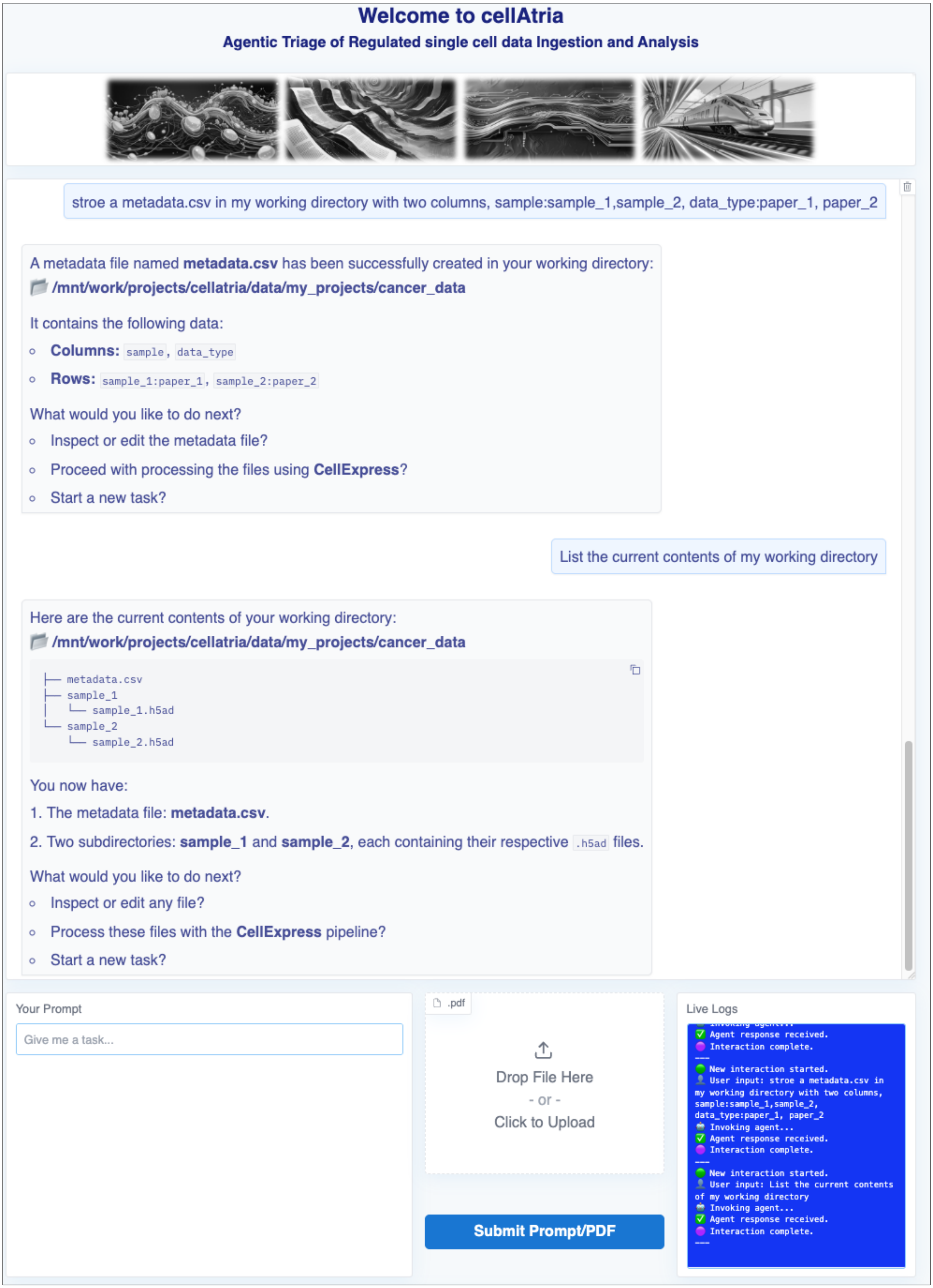


**Supplementary Figure S18. Metadata file creation using CellAtria.** Upon user request, the agent generates a structured metadata file based on the provided inputs. The live log viewer provides real-time feedback on execution status and confirms successful completion. Note: the system-level panels (backend, terminal, file browser, and chat export) have been cropped for improved visibility.

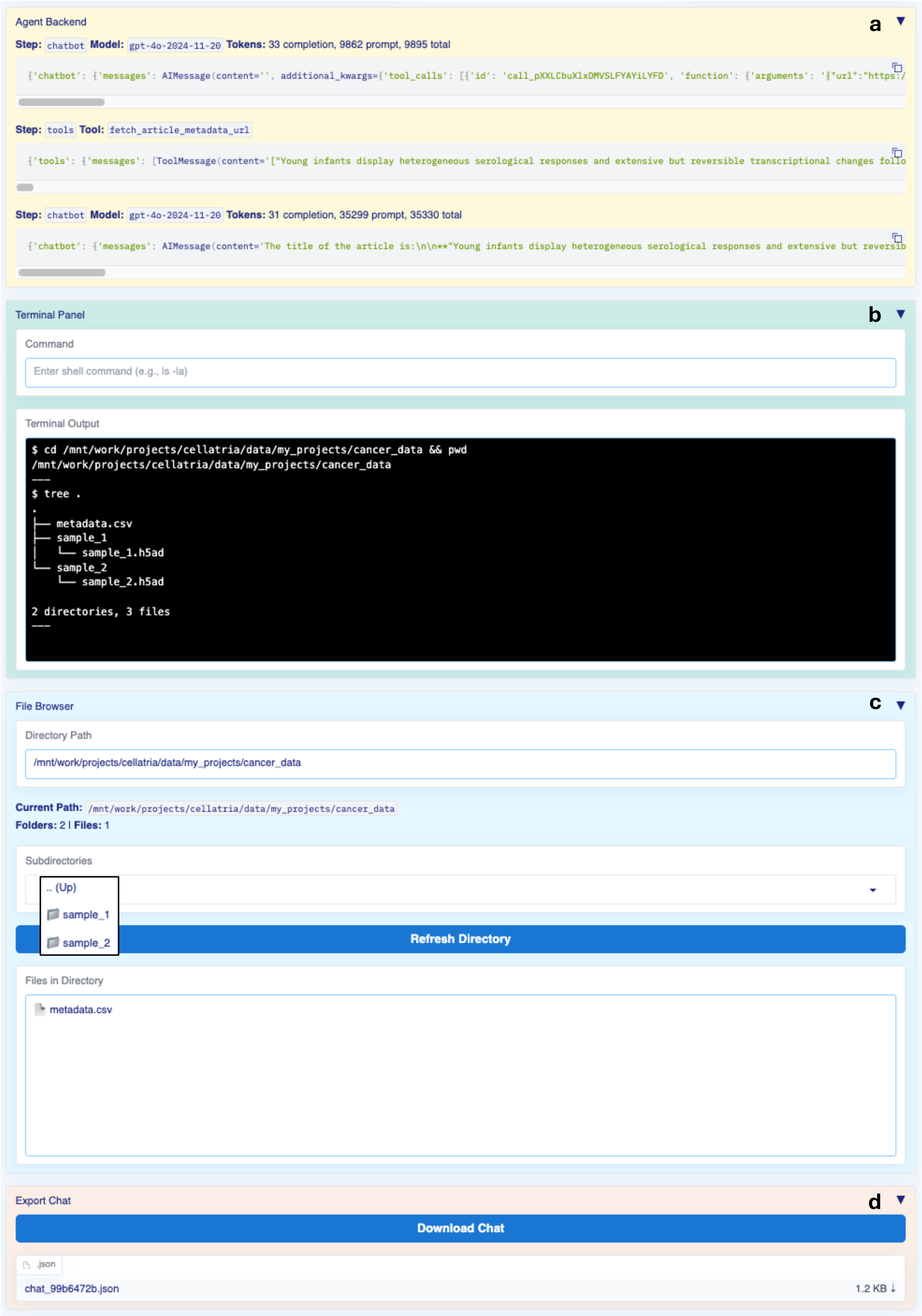


**Supplementary Figure S19. System-level panels of the CellAtria user interface. a)** The "Agent Backend" panel displays step-by-step agent reasoning, tool calls, and backend responses. **b)** The "Terminal Panel" allows execution of shell commands within the agent runtime. **c)** The "File Browser" provides a navigable view of the directory structure. "Refresh Directory" updates the view when a new "Directory Path" is provided, and "Files in Directory" lists the files at that location. **d)** The "Export Chat" allows users to download the agent-user interaction via the "Download Chat" button. Note: The header and chatbot window were cropped to enhance panel visibility.

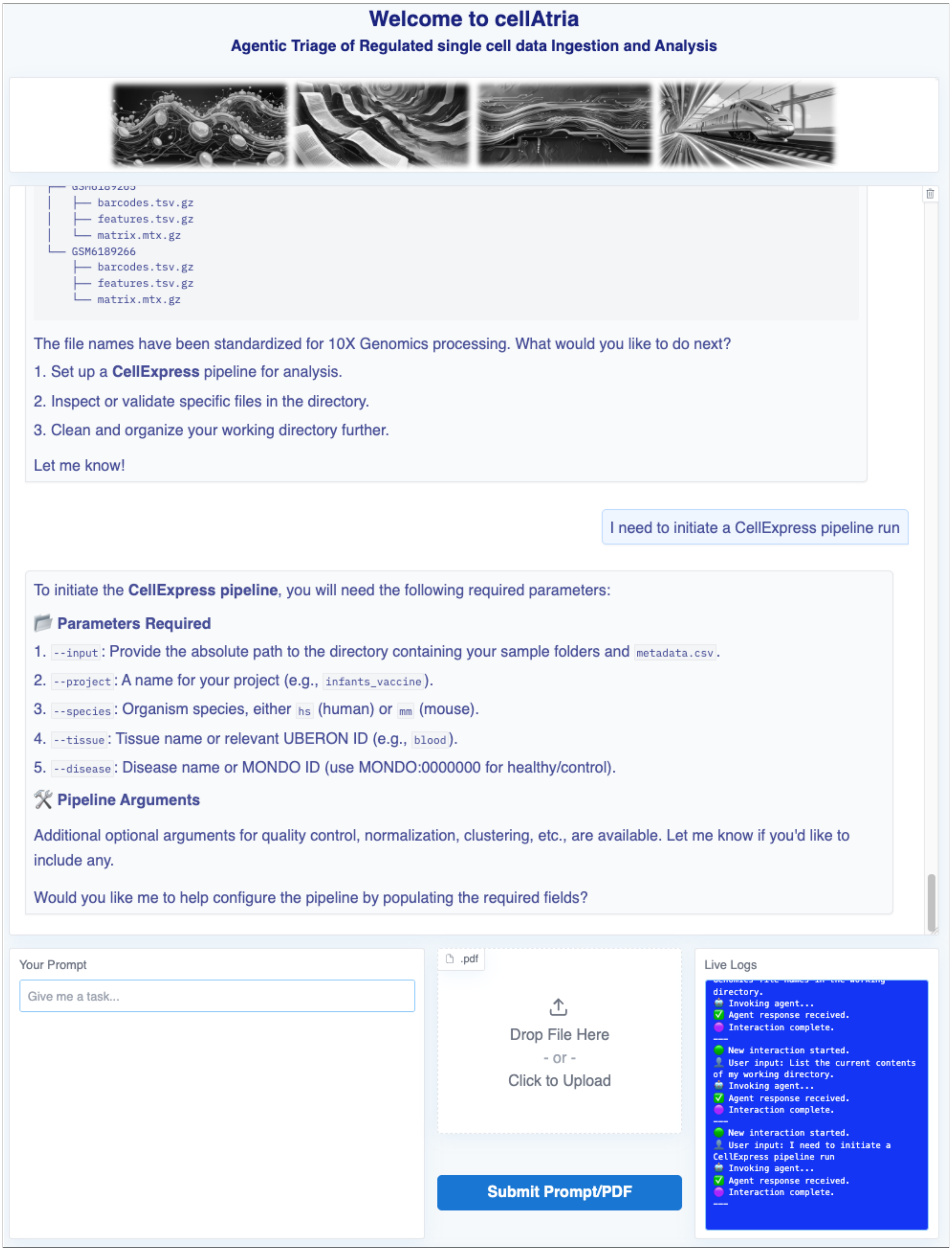


**Supplementary Figure S20. Initialization of the CellExpress analysis pipeline.** In response to user input, CellAtria displays the required parameters for initiating CellExpress and prompts the user to specify additional configurable options, enabling tailored pipeline execution. Note: the integrated live log viewer provides real-time feedback on execution status and confirms successful completion. The system-level panels (backend, terminal, file browser, and chat export) have been cropped for improved visibility.

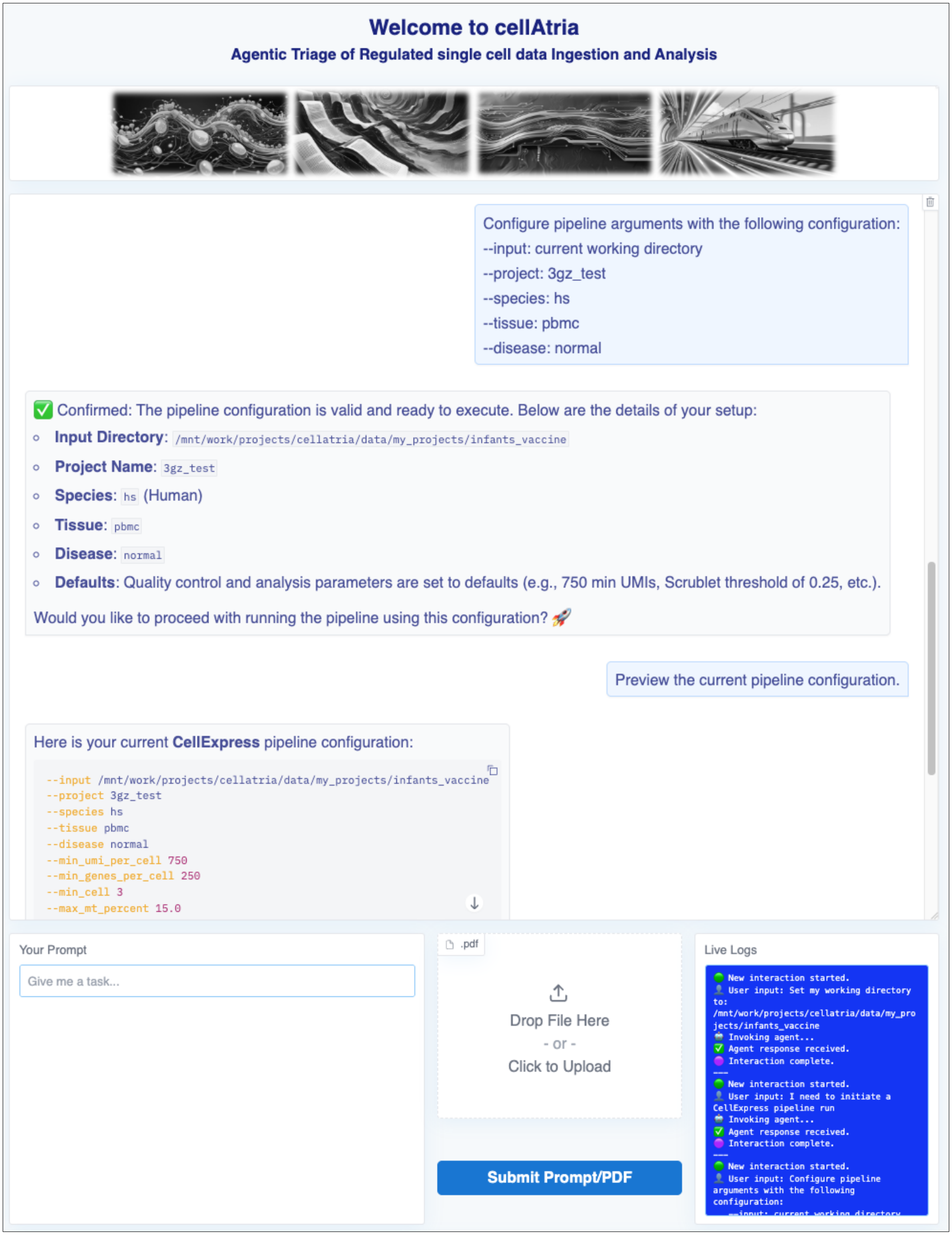


**Supplementary Figure S21. CellExpress pipeline configuration preview and validation.** The agent validates user-provided parameters for the CellExpress pipeline and confirms readiness for execution. Default quality control thresholds are applied automatically. A YAML-formatted configuration summary is presented to the user for final review prior to pipeline initiation. The integrated live log viewer provides real-time feedback on execution status and confirms successful completion. Note: the system-level panels (backend, terminal, file browser, and chat export) have been cropped for improved visibility.

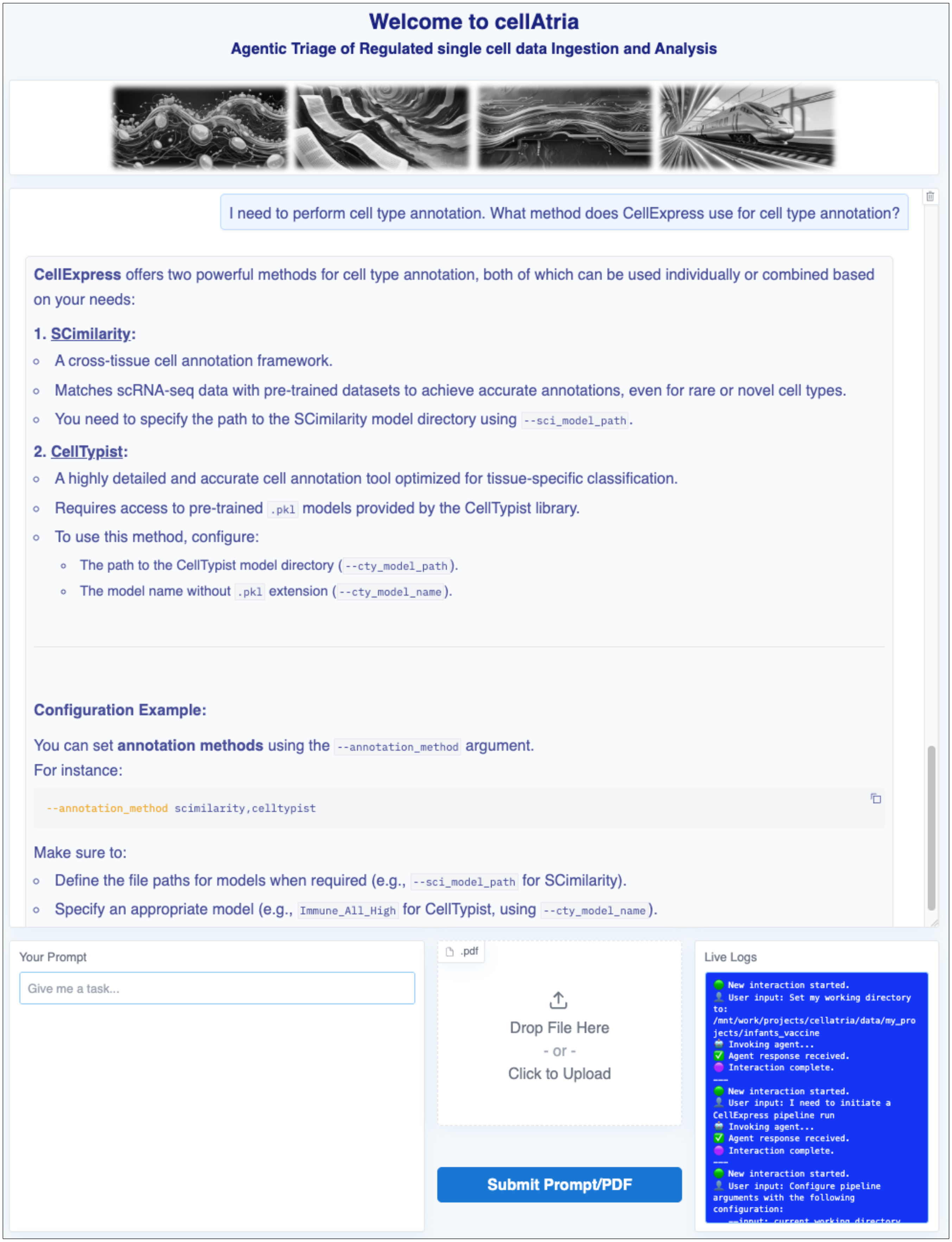


**Supplementary Figure S22. Cell type annotation strategies in CellExpress.** Upon user request, the agent enumerates the automated cell type annotation methods supported by CellExpress. The provided configuration example illustrates how users specify annotation strategies and corresponding model paths through pipeline arguments. The integrated live log viewer provides real-time feedback on execution status and confirms successful completion. Note: the system-level panels (backend, terminal, file browser, and chat export) have been cropped for improved visibility.

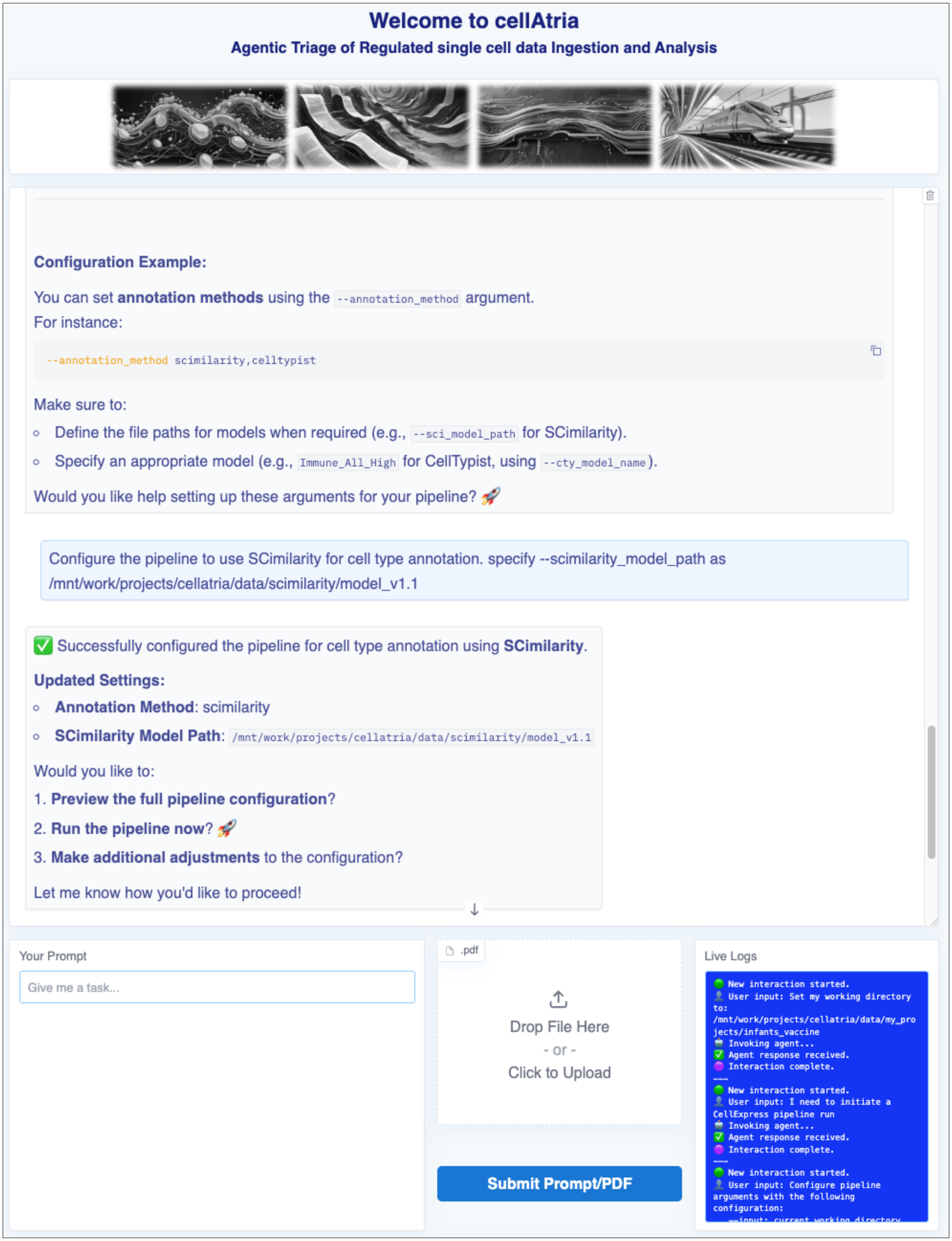


**Supplementary Figure S23. Configuration of SCimilarity-based cell type annotation in CellExpress.** In response to user instruction, the agent configures the CellExpress pipeline to utilize SCimilarity for automated cell type annotation. The interface confirms the updated parameters and presents follow-up actionable options to proceed with execution. The integrated live log viewer provides real-time feedback on execution status and confirms successful completion. Note: the system-level panels (backend, terminal, file browser, and chat export) have been cropped for improved visibility.

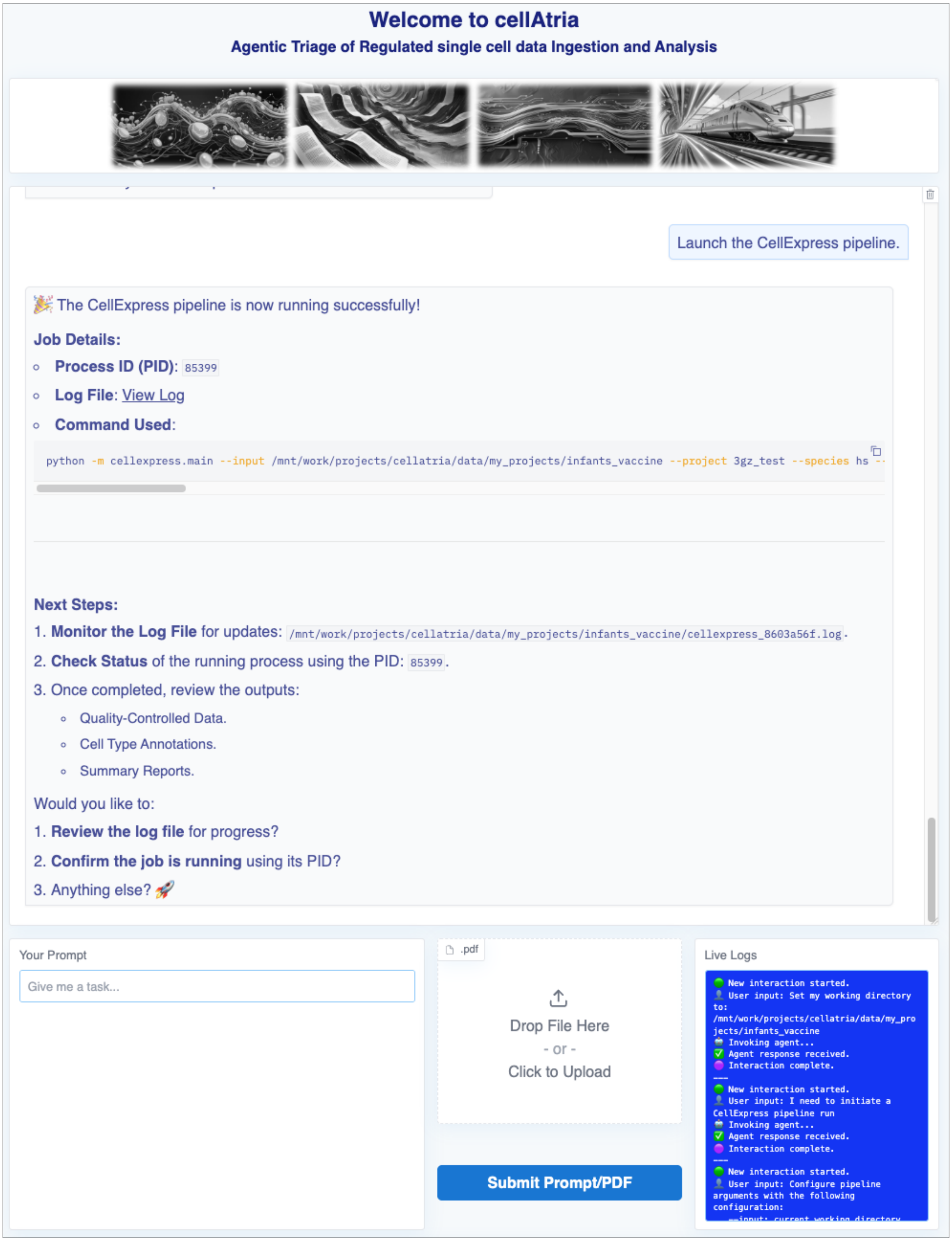


**Supplementary Figure S24. Launch and background execution of the CellExpress pipeline.** Upon receiving the execution request, the agent initiates the CellExpress pipeline in detached mode using the specified configuration. The process ID (PID) and log file path are displayed to facilitate runtime monitoring. The integrated live log viewer provides real-time feedback on execution status and confirms successful completion. Note: the system-level panels (backend, terminal, file browser, and chat export) have been cropped for improved visibility.

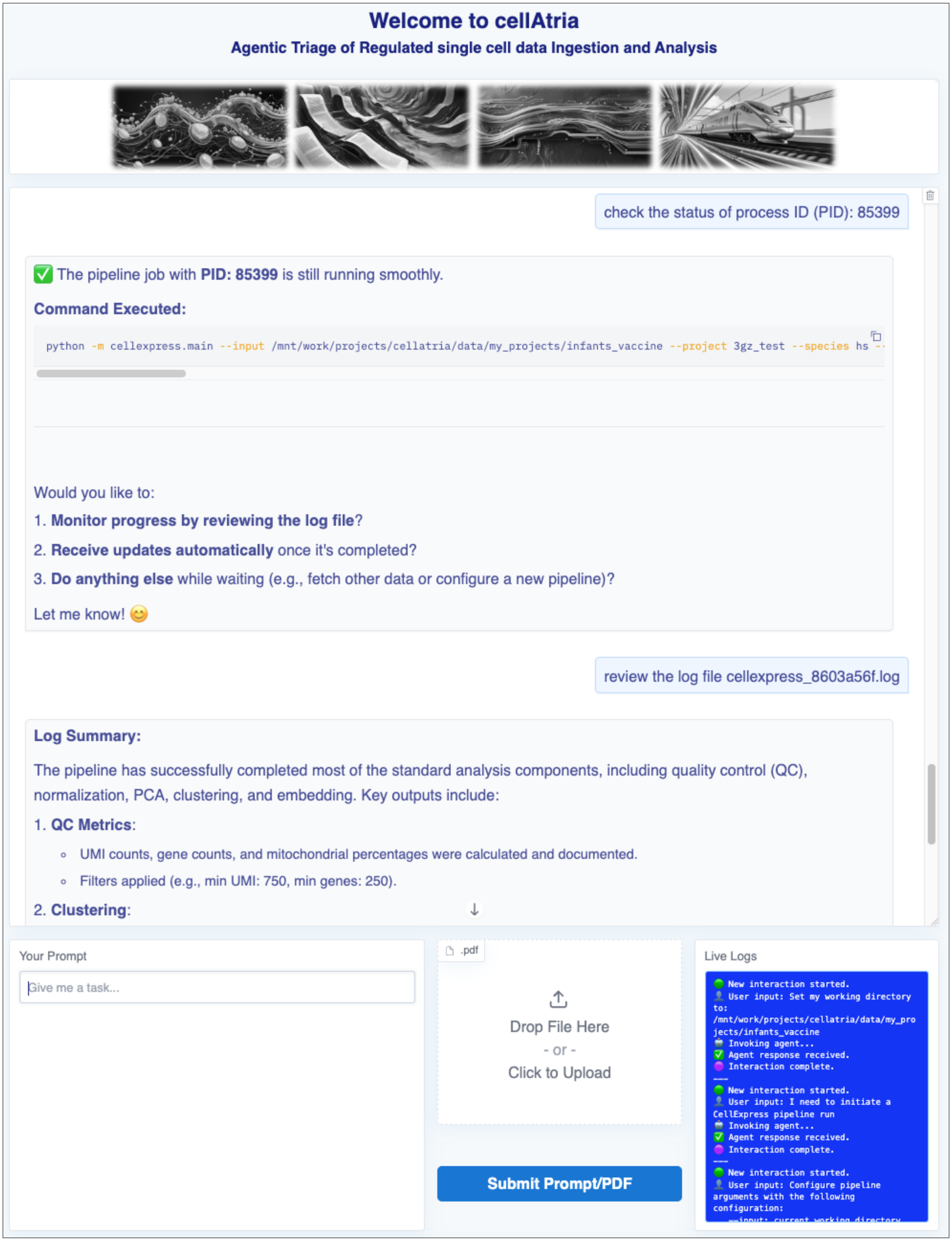


**Supplementary Figure S25. Real-time monitoring and log inspection of an active CellExpress pipeline.** Given the process ID, the agent confirms the active execution status of the CellExpress pipeline, displays the full launch command, and provides access to the log file for real-time inspection. Upon user request, the agent parses the log to summarize completed stages of the pipeline. The integrated live log viewer provides real-time feedback on execution status and confirms successful completion. Note: the system-level panels (backend, terminal, file browser, and chat export) have been cropped for improved visibility.

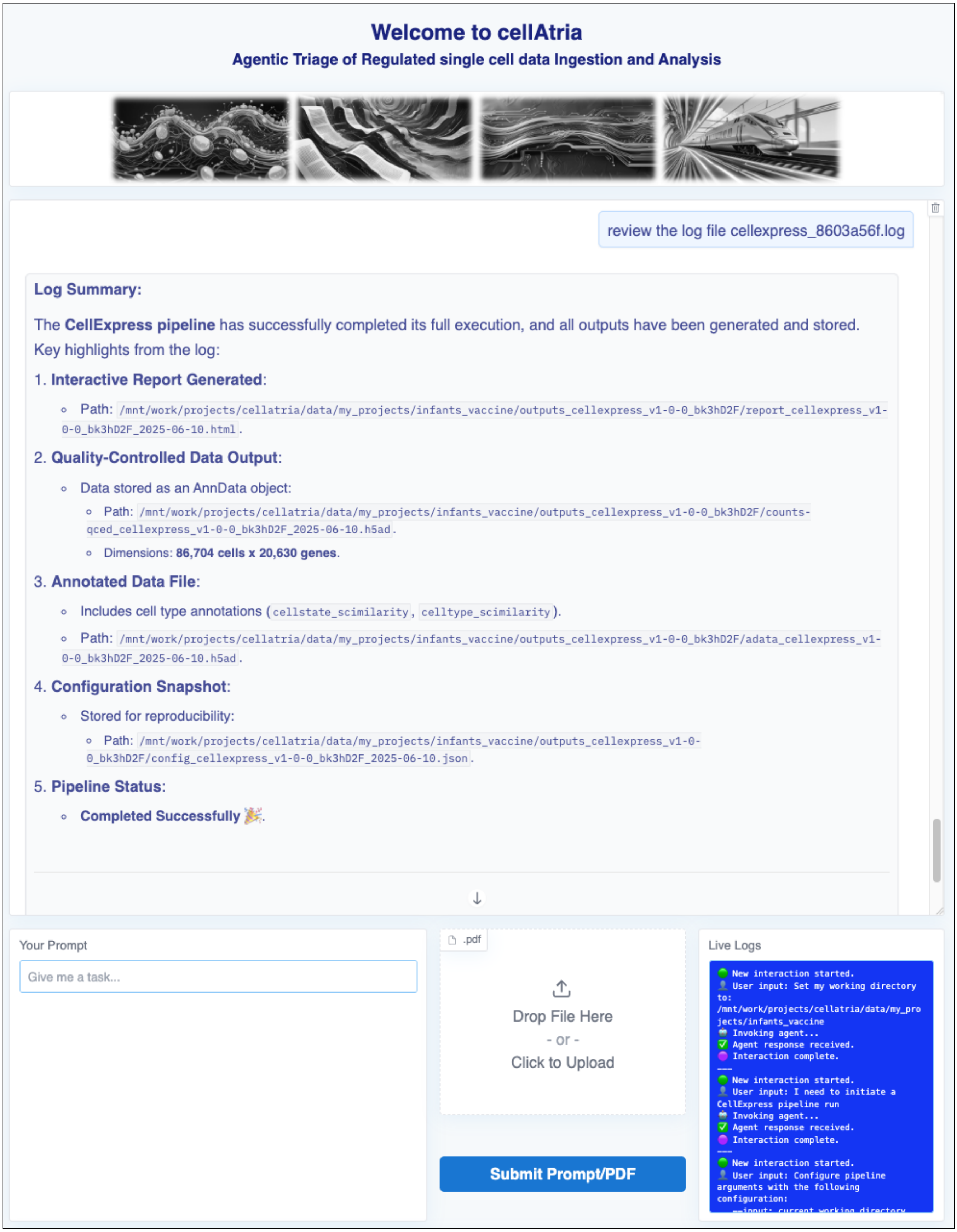


**Supplementary Figure S26. Completion of the CellExpress pipeline and structured output verification.** Upon user request, the agent confirms successful completion of the CellExpress pipeline and enumerates the resulting output files. The integrated live log viewer provides real-time feedback on execution status and confirms successful completion. Note: the system-level panels (backend, terminal, file browser, and chat export) have been cropped for improved visibility.

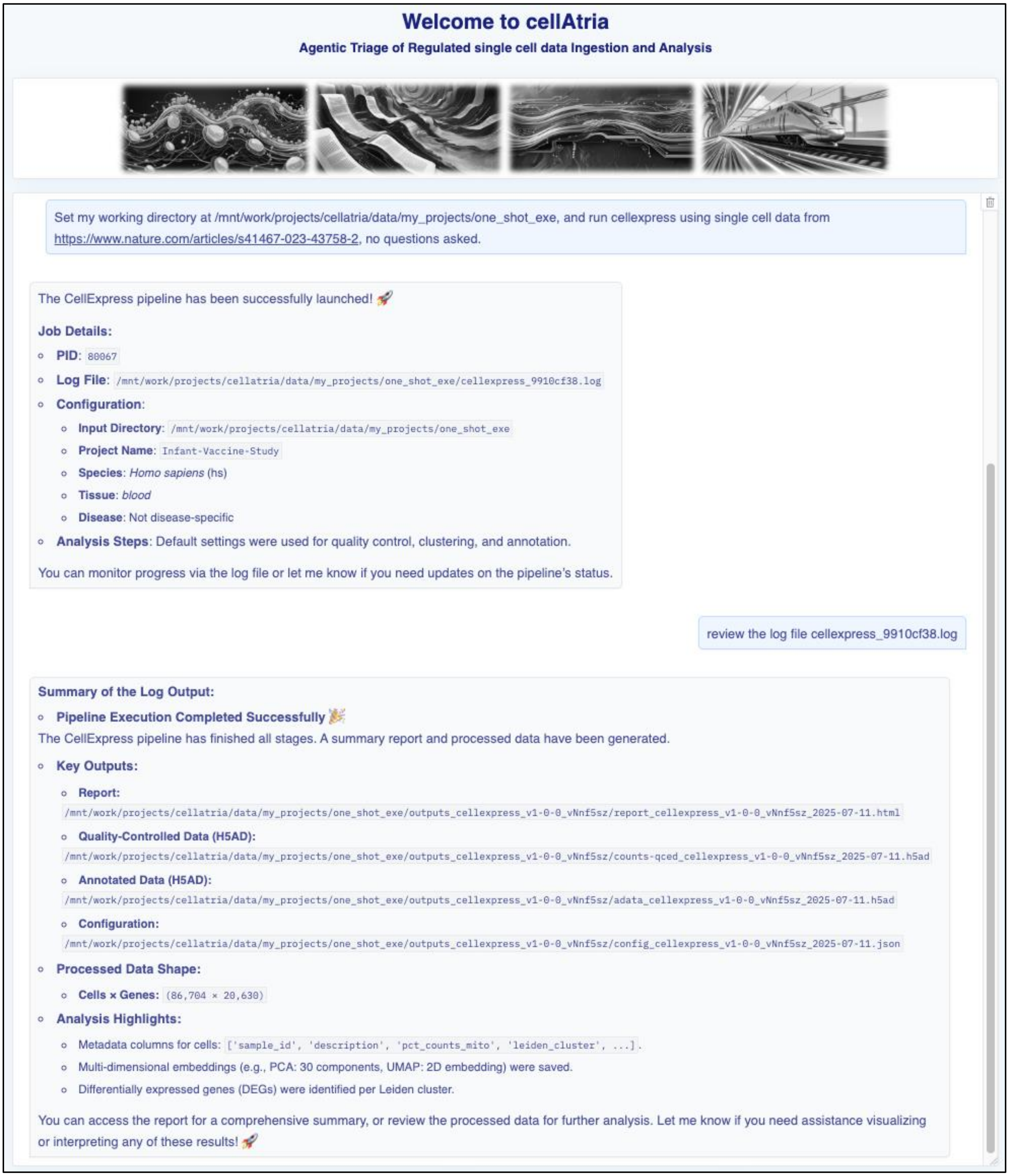


**Supplementary Figure S27. Autonomous execution of the CellExpress pipeline initiated by a single prompt.** The CellAtria interface demonstrates complete orchestration of the document-to-analysis workflow following submission of a high-level user prompt - without iterative user-agent interactions. Note: the log viewer and the system-level panels (backend, terminal, file browser, and chat export) have been cropped for improved visibility.

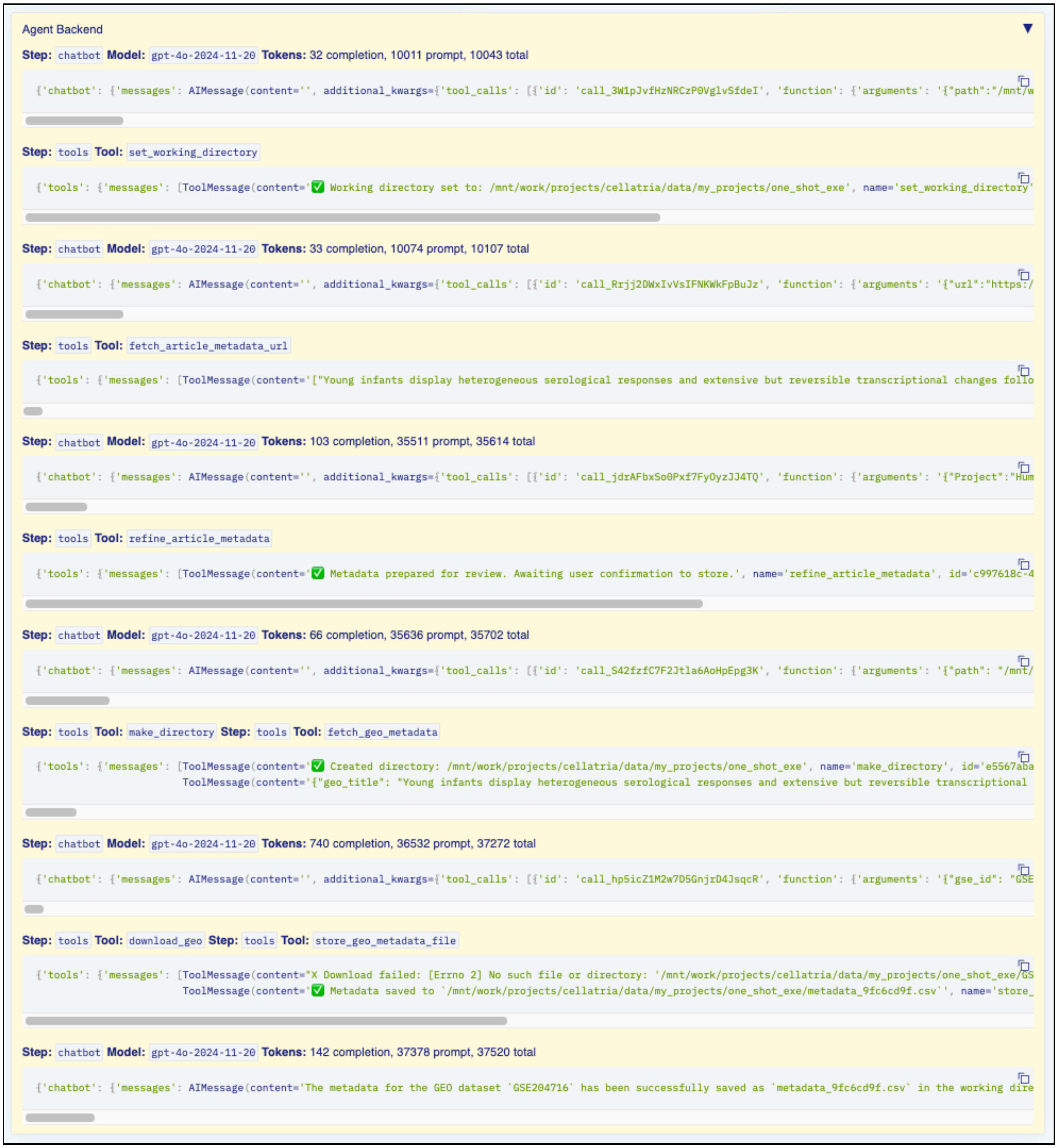


**Supplementary Figure S28. Agentic backend reasoning trace for one-shot execution workflow.** Internal execution trace showing stepwise agent behavior as it interprets user intent and autonomously orchestrates tool invocations without requiring follow-up prompts. Each step documents how the agent decomposed a single compound prompt into a series of reasoning and execution actions. "chatbot" entries reflect language model outputs with tool-calling instructions, while "tools" steps record the successful execution of those calls. Metadata such as model version, token usage, and generated text is included for transparency and reproducibility. Note: the header, chatbot window, log viewer, and system-level panels (including backend trace, terminal, file browser, and chat export) have been cropped to enhance visual clarity.
