## Supplementary material for "An Agentic AI Framework for Ingestion and Standardization of Single-Cell RNA-seq Data Analysis": Supplementary_Material_S2.html

  Newborn\_PBMC\_Immunization


### Newborn\_PBMC\_Immunization

###### Identifier: cellexpress\_v1-0-0\_ziFzM4S

###### Date: 23 July, 2025 | Runtime: 16.03 min


---

### 1 ExpressSummary

Read me

#### 1.1 Sample & Experimental Context

| Species | Disease | Tissue |
| --- | --- | --- |
| human | normal | blood |

---

#### 1.2 Cell Count & Expression Summary

---

### 2 ExpressSettings

Read me

| Argument | Description | Entered Value |
| --- | --- | --- |
| species | Species type used in the analysis. | hs |
| tissue | Tissue type for the dataset. | blood |
| disease | Disease name | normal |
| min\_umi\_per\_cell | Minimum UMI counts required per cell. | {‘S1’: 750, ‘S2’: 750, ‘S3’: 750, ‘S4’: 750, ‘S5’: 750, ‘S6’: 750, ‘S7’: 750, ‘S8’: 750, ‘S9’: 750, ‘S10’: 750, ‘S11’: 750, ‘S12’: 750, ‘S13’: 750, ‘S14’: 750, ‘S15’: 750, ‘S16’: 750, ‘S17’: 750, ‘S18’: 750} |
| max\_umi\_per\_cell | Maximum UMI counts allowed per cell. | {‘S1’: inf, ‘S2’: inf, ‘S3’: inf, ‘S4’: inf, ‘S5’: inf, ‘S6’: inf, ‘S7’: inf, ‘S8’: inf, ‘S9’: inf, ‘S10’: inf, ‘S11’: inf, ‘S12’: inf, ‘S13’: inf, ‘S14’: inf, ‘S15’: inf, ‘S16’: inf, ‘S17’: inf, ‘S18’: inf} |
| min\_genes\_per\_cell | Minimum number of detected genes per cell. | {‘S1’: 250, ‘S2’: 250, ‘S3’: 250, ‘S4’: 250, ‘S5’: 250, ‘S6’: 250, ‘S7’: 250, ‘S8’: 250, ‘S9’: 250, ‘S10’: 250, ‘S11’: 250, ‘S12’: 250, ‘S13’: 250, ‘S14’: 250, ‘S15’: 250, ‘S16’: 250, ‘S17’: 250, ‘S18’: 250} |
| max\_genes\_per\_cell | Maximum number of detected genes per cell. | {‘S1’: inf, ‘S2’: inf, ‘S3’: inf, ‘S4’: inf, ‘S5’: inf, ‘S6’: inf, ‘S7’: inf, ‘S8’: inf, ‘S9’: inf, ‘S10’: inf, ‘S11’: inf, ‘S12’: inf, ‘S13’: inf, ‘S14’: inf, ‘S15’: inf, ‘S16’: inf, ‘S17’: inf, ‘S18’: inf} |
| min\_cell | Minimum number of cells in which a gene must be detected to be retained. | {‘S1’: 3, ‘S2’: 3, ‘S3’: 3, ‘S4’: 3, ‘S5’: 3, ‘S6’: 3, ‘S7’: 3, ‘S8’: 3, ‘S9’: 3, ‘S10’: 3, ‘S11’: 3, ‘S12’: 3, ‘S13’: 3, ‘S14’: 3, ‘S15’: 3, ‘S16’: 3, ‘S17’: 3, ‘S18’: 3} |
| max\_mt\_percent | Maximum allowed percentage of mitochondrial content per cell. | {‘S1’: 15.0, ‘S2’: 15.0, ‘S3’: 15.0, ‘S4’: 15.0, ‘S5’: 15.0, ‘S6’: 15.0, ‘S7’: 15.0, ‘S8’: 15.0, ‘S9’: 15.0, ‘S10’: 15.0, ‘S11’: 15.0, ‘S12’: 15.0, ‘S13’: 15.0, ‘S14’: 15.0, ‘S15’: 15.0, ‘S16’: 15.0, ‘S17’: 15.0, ‘S18’: 15.0} |
| doublet\_method | Method used for doublet identification. | scrublet |
| scrublet\_cutoff | Threshold for filtering doublets based on Scrublet score. | 0.25 |
| norm\_target\_sum | Target sum for normalization, defining the total counts per cell. | 10000.0 |
| n\_top\_genes | Number of highly variable genes to retain for downstream analysis. | 2000 |
| regress\_out | Whether total counts and mitochondrial percentages are regressed out. | yes |
| scale\_max\_value | Maximum value for data scaling. | 10 |
| n\_pcs | Number of principal components used in PCA. | 30 |
| batch\_correction | Batch correction method applied. | harmony |
| batch\_vars | Metadata column(s) used for batch correction. | sample\_id |
| n\_neighbors | Number of neighbors used for kNN graph construction. | 15 |
| resolution | Resolution parameter for Leiden clustering. | 0.6 |
| compute\_tsne | Indicates whether t-SNE embedding is computed. | yes |
| annotation\_method | Method used for cell annotation. | scimilarity,celltypist |
| sci\_model\_path | Path to the SCimilarity model for annotation. | /mnt/work/projects/cellatria/data/scimilarity/model\_v1.1 |
| cty\_model\_path | Path to the Celltypist model for annotation. | /mnt/work/projects/cellatria/data/celltypist/model\_v1.6.3 |
| cty\_model\_name | Celltypist model used. | Immune\_All\_Low |
| pval\_threshold | P-value threshold for filtering differentially expressed genes (DEGs). | 0.05 |
| logfc\_threshold | Log fold-change (logFC) threshold for filtering DEGs. | 0.25 |
| dea\_method | Statistical test used for differential expression analysis. | wilcoxon |
| top\_n\_deg\_leidn | Number of top differentially expressed genes (DEGs) to return per ‘leiden’ clusters. | 100 |
| top\_n\_deg\_scim | Number of top differentially expressed genes (DEGs) to return per ‘scimilarity’ annotated cells. | 100 |
| top\_n\_deg\_cltpst | Number of top differentially expressed genes (DEGs) to return per ‘celltypist’ annotated cells. | 100 |
| pts\_threshold | Minimum fraction of cells expressing a gene for it to be considered a DEG. | 0.1 |
| doc\_url | URL to the published documentation associated with the analysis. | https://www.nature.com/articles/s41467-023-43758-2 |
| fix\_gene\_names | Fix gene names if Ensembl IDs are detected. | no |
| limit\_threads | Apply thread limits to avoid memory crashes | 1 |

---

### 3 ExpressQC

Read me

#### 3.1 Sample-level Characteristics and Quality Control Metrics

##### 3.1.1 Metadata

| sample | sample\_id |
| --- | --- |
| GSM6189249 | S1 |
| GSM6189250 | S2 |
| GSM6189251 | S3 |
| GSM6189252 | S4 |
| GSM6189253 | S5 |
| GSM6189254 | S6 |
| GSM6189255 | S7 |
| GSM6189256 | S8 |
| GSM6189257 | S9 |
| GSM6189258 | S10 |
| GSM6189259 | S11 |
| GSM6189260 | S12 |
| GSM6189261 | S13 |
| GSM6189262 | S14 |
| GSM6189263 | S15 |
| GSM6189264 | S16 |
| GSM6189265 | S17 |
| GSM6189266 | S18 |

##### 3.1.2 Cells & Genes

| sample | sample\_id | pre\_qc\_gene | pre\_qc\_cell | post\_qc\_gene | post\_qc\_cell |
| --- | --- | --- | --- | --- | --- |
| GSM6189249 | S1 | 20,891 | 6,921 | 17,382 | 3,994 |
| GSM6189250 | S2 | 21,246 | 10,918 | 18,086 | 6,737 |
| GSM6189251 | S3 | 21,339 | 8,433 | 18,160 | 6,327 |
| GSM6189252 | S4 | 21,255 | 10,241 | 17,963 | 6,710 |
| GSM6189253 | S5 | 20,769 | 8,359 | 16,376 | 3,249 |
| GSM6189254 | S6 | 21,476 | 11,364 | 18,252 | 8,226 |
| GSM6189255 | S7 | 20,062 | 5,151 | 16,728 | 3,047 |
| GSM6189256 | S8 | 18,979 | 3,523 | 15,416 | 1,860 |
| GSM6189257 | S9 | 20,485 | 5,156 | 16,916 | 2,971 |
| GSM6189258 | S10 | 19,545 | 4,100 | 16,021 | 2,380 |
| GSM6189259 | S11 | 20,467 | 6,138 | 17,047 | 3,795 |
| GSM6189260 | S12 | 20,364 | 5,098 | 16,870 | 3,098 |
| GSM6189261 | S13 | 20,047 | 4,475 | 16,789 | 3,005 |
| GSM6189262 | S14 | 18,662 | 1,459 | 15,213 | 919 |
| GSM6189263 | S15 | 19,920 | 5,125 | 16,673 | 3,139 |
| GSM6189264 | S16 | 20,472 | 5,523 | 17,079 | 3,688 |
| GSM6189265 | S17 | 20,914 | 5,348 | 17,513 | 3,854 |
| GSM6189266 | S18 | 20,554 | 6,159 | 17,091 | 4,102 |

##### 3.1.3 UMI per cell

Vales are post-QC.

| sample | sample\_id | min | q0 | q25 | q50 | q75 | q100 | max |
| --- | --- | --- | --- | --- | --- | --- | --- | --- |
| GSM6189249 | S1 | 764 | 764 | 5,237 | 6,616 | 8,143 | 27,116 | 27,116 |
| GSM6189250 | S2 | 757 | 757 | 4,655 | 5,847 | 7,181 | 43,566 | 43,566 |
| GSM6189251 | S3 | 756 | 756 | 4,944 | 6,186 | 7,588 | 58,054 | 58,054 |
| GSM6189252 | S4 | 761 | 761 | 4,529 | 5,646 | 7,056 | 57,462 | 57,462 |
| GSM6189253 | S5 | 750 | 750 | 1,265 | 2,076 | 3,974 | 52,899 | 52,899 |
| GSM6189254 | S6 | 751 | 751 | 4,149 | 5,382 | 6,844 | 42,904 | 42,904 |
| GSM6189255 | S7 | 786 | 786 | 5,162 | 6,700 | 8,226 | 29,128 | 29,128 |
| GSM6189256 | S8 | 750 | 750 | 3,684 | 5,716 | 7,362 | 53,772 | 53,772 |
| GSM6189257 | S9 | 758 | 758 | 5,026 | 6,795 | 8,512 | 61,753 | 61,753 |
| GSM6189258 | S10 | 751 | 751 | 3,812 | 6,080 | 7,905 | 84,052 | 84,052 |
| GSM6189259 | S11 | 751 | 751 | 4,285 | 5,935 | 7,516 | 99,574 | 99,574 |
| GSM6189260 | S12 | 753 | 753 | 4,909 | 6,483 | 8,129 | 45,268 | 45,268 |
| GSM6189261 | S13 | 760 | 760 | 4,204 | 5,564 | 6,747 | 32,149 | 32,149 |
| GSM6189262 | S14 | 991 | 991 | 5,843 | 7,439 | 9,277 | 41,415 | 41,415 |
| GSM6189263 | S15 | 755 | 755 | 4,719 | 5,977 | 7,332 | 38,681 | 38,681 |
| GSM6189264 | S16 | 751 | 751 | 4,347 | 5,986 | 7,525 | 82,498 | 82,498 |
| GSM6189265 | S17 | 752 | 752 | 4,518 | 6,134 | 8,228 | 89,182 | 89,182 |
| GSM6189266 | S18 | 753 | 753 | 4,012 | 5,414 | 6,914 | 40,118 | 40,118 |

##### 3.1.4 Gene per cell

Vales are post-QC.

| sample | sample\_id | min | q0 | q25 | q50 | q75 | q100 | max |
| --- | --- | --- | --- | --- | --- | --- | --- | --- |
| GSM6189249 | S1 | 303 | 303 | 1,565 | 1,779 | 2,022 | 4,621 | 4,621 |
| GSM6189250 | S2 | 405 | 405 | 1,549 | 1,784 | 2,052 | 5,429 | 5,429 |
| GSM6189251 | S3 | 346 | 346 | 1,522 | 1,726 | 1,966 | 5,622 | 5,622 |
| GSM6189252 | S4 | 367 | 367 | 1,395 | 1,603 | 1,888 | 6,527 | 6,527 |
| GSM6189253 | S5 | 251 | 251 | 725 | 1,056 | 1,546 | 6,306 | 6,306 |
| GSM6189254 | S6 | 253 | 253 | 1,372 | 1,576 | 1,824 | 5,595 | 5,595 |
| GSM6189255 | S7 | 336 | 336 | 1,472 | 1,692 | 1,930 | 4,703 | 4,703 |
| GSM6189256 | S8 | 352 | 352 | 1,272 | 1,622 | 1,918 | 6,641 | 6,641 |
| GSM6189257 | S9 | 257 | 257 | 1,536 | 1,798 | 2,064 | 5,852 | 5,852 |
| GSM6189258 | S10 | 258 | 258 | 1,281 | 1,635 | 1,898 | 8,324 | 8,324 |
| GSM6189259 | S11 | 250 | 250 | 1,303 | 1,591 | 1,878 | 7,383 | 7,383 |
| GSM6189260 | S12 | 276 | 276 | 1,442 | 1,688 | 1,957 | 5,130 | 5,130 |
| GSM6189261 | S13 | 290 | 290 | 1,322 | 1,541 | 1,769 | 5,030 | 5,030 |
| GSM6189262 | S14 | 449 | 449 | 1,634 | 1,904 | 2,200 | 5,739 | 5,739 |
| GSM6189263 | S15 | 250 | 250 | 1,339 | 1,541 | 1,766 | 5,068 | 5,068 |
| GSM6189264 | S16 | 254 | 254 | 1,341 | 1,570 | 1,880 | 7,114 | 7,114 |
| GSM6189265 | S17 | 259 | 259 | 1,474 | 1,802 | 2,289 | 6,891 | 6,891 |
| GSM6189266 | S18 | 261 | 261 | 1,260 | 1,492 | 1,768 | 5,273 | 5,273 |

##### 3.1.5 Mitochondrial (%)

Vales are post-QC.

| sample | sample\_id | min | q0 | q25 | q50 | q75 | q100 | max |
| --- | --- | --- | --- | --- | --- | --- | --- | --- |
| GSM6189249 | S1 | 0.0% | 0.0% | 6.1% | 7.3% | 8.9% | 15.0% | 15.0% |
| GSM6189250 | S2 | 0.3% | 0.3% | 4.6% | 5.6% | 6.9% | 15.0% | 15.0% |
| GSM6189251 | S3 | 0.1% | 0.1% | 5.8% | 6.9% | 8.2% | 15.0% | 15.0% |
| GSM6189252 | S4 | 0.1% | 0.1% | 6.7% | 7.9% | 9.3% | 15.0% | 15.0% |
| GSM6189253 | S5 | 0.0% | 0.0% | 3.7% | 6.2% | 9.9% | 15.0% | 15.0% |
| GSM6189254 | S6 | 0.0% | 0.0% | 4.6% | 5.5% | 6.8% | 15.0% | 15.0% |
| GSM6189255 | S7 | 0.1% | 0.1% | 4.6% | 5.6% | 7.1% | 15.0% | 15.0% |
| GSM6189256 | S8 | 0.1% | 0.1% | 3.7% | 4.7% | 6.4% | 14.9% | 14.9% |
| GSM6189257 | S9 | 0.0% | 0.0% | 4.1% | 5.1% | 6.7% | 15.0% | 15.0% |
| GSM6189258 | S10 | 0.1% | 0.1% | 4.0% | 5.2% | 7.2% | 15.0% | 15.0% |
| GSM6189259 | S11 | 0.0% | 0.0% | 3.7% | 4.7% | 6.2% | 15.0% | 15.0% |
| GSM6189260 | S12 | 0.0% | 0.0% | 4.1% | 5.0% | 6.6% | 15.0% | 15.0% |
| GSM6189261 | S13 | 0.3% | 0.3% | 5.5% | 6.7% | 8.1% | 15.0% | 15.0% |
| GSM6189262 | S14 | 0.0% | 0.0% | 4.6% | 5.6% | 7.0% | 14.9% | 14.9% |
| GSM6189263 | S15 | 0.0% | 0.0% | 5.1% | 6.3% | 7.7% | 15.0% | 15.0% |
| GSM6189264 | S16 | 0.0% | 0.0% | 5.7% | 7.0% | 8.7% | 15.0% | 15.0% |
| GSM6189265 | S17 | 0.0% | 0.0% | 4.5% | 5.5% | 6.7% | 15.0% | 15.0% |
| GSM6189266 | S18 | 0.2% | 0.2% | 5.2% | 6.5% | 8.3% | 15.0% | 15.0% |

---

#### 3.2 Distribution Plots for Quality Control Metrics

Thresholds, represented by dark-red dashed lines, were implemented to filter the data and only retain cells of high quality.

##### 3.2.1 UMI per cell distribution

##### 3.2.2 Gene per cell distribution

##### 3.2.3 Mitochondrial (%) per cell distribution

##### 3.2.4 Total UMI Counts vs. Genes Detected per cell

The gray dashed line indicates the identity line (y = x)

---

#### 3.3 Barcodes Contamination

##### 3.3.1 Count - before QC

Number of barcodes shared between pairs of samples pre-QC.

|  | S1 | S2 | S3 | S4 | S5 | S6 | S7 | S8 | S9 | S10 | S11 | S12 | S13 | S14 | S15 | S16 | S17 | S18 |
| --- | --- | --- | --- | --- | --- | --- | --- | --- | --- | --- | --- | --- | --- | --- | --- | --- | --- | --- |
| S1 | 6921 | 18 | 21 | 25 | 10 | 20 | 13 | 5 | 8 | 5 | 12 | 8 | 13 | 3 | 7 | 12 | 12 | 9 |
| S2 | 18 | 10918 | 24 | 30 | 30 | 23 | 12 | 7 | 21 | 20 | 23 | 13 | 13 | 5 | 21 | 15 | 15 | 17 |
| S3 | 21 | 24 | 8433 | 12 | 15 | 31 | 14 | 15 | 4 | 7 | 16 | 14 | 11 | 5 | 11 | 9 | 8 | 8 |
| S4 | 25 | 30 | 12 | 10241 | 22 | 38 | 10 | 5 | 11 | 12 | 23 | 16 | 10 | 5 | 23 | 11 | 14 | 18 |
| S5 | 10 | 30 | 15 | 22 | 8359 | 33 | 11 | 10 | 13 | 8 | 13 | 12 | 12 | 1 | 10 | 11 | 8 | 20 |
| S6 | 20 | 23 | 31 | 38 | 33 | 11364 | 16 | 14 | 14 | 14 | 22 | 19 | 10 | 4 | 18 | 15 | 18 | 27 |
| S7 | 13 | 12 | 14 | 10 | 11 | 16 | 5151 | 2 | 6 | 4 | 10 | 6 | 9 | 2 | 7 | 8 | 9 | 8 |
| S8 | 5 | 7 | 15 | 5 | 10 | 14 | 2 | 3523 | 4 | 8 | 4 | 3 | 7 | 0 | 6 | 7 | 3 | 4 |
| S9 | 8 | 21 | 4 | 11 | 13 | 14 | 6 | 4 | 5156 | 8 | 5 | 1 | 6 | 1 | 2 | 6 | 10 | 8 |
| S10 | 5 | 20 | 7 | 12 | 8 | 14 | 4 | 8 | 8 | 4100 | 7 | 8 | 3 | 1 | 10 | 6 | 10 | 7 |
| S11 | 12 | 23 | 16 | 23 | 13 | 22 | 10 | 4 | 5 | 7 | 6138 | 8 | 8 | 4 | 14 | 10 | 11 | 11 |
| S12 | 8 | 13 | 14 | 16 | 12 | 19 | 6 | 3 | 1 | 8 | 8 | 5098 | 8 | 7 | 6 | 10 | 4 | 5 |
| S13 | 13 | 13 | 11 | 10 | 12 | 10 | 9 | 7 | 6 | 3 | 8 | 8 | 4475 | 1 | 4 | 9 | 7 | 11 |
| S14 | 3 | 5 | 5 | 5 | 1 | 4 | 2 | 0 | 1 | 1 | 4 | 7 | 1 | 1459 | 4 | 2 | 2 | 5 |
| S15 | 7 | 21 | 11 | 23 | 10 | 18 | 7 | 6 | 2 | 10 | 14 | 6 | 4 | 4 | 5125 | 8 | 4 | 17 |
| S16 | 12 | 15 | 9 | 11 | 11 | 15 | 8 | 7 | 6 | 6 | 10 | 10 | 9 | 2 | 8 | 5523 | 10 | 15 |
| S17 | 12 | 15 | 8 | 14 | 8 | 18 | 9 | 3 | 10 | 10 | 11 | 4 | 7 | 2 | 4 | 10 | 5348 | 49 |
| S18 | 9 | 17 | 8 | 18 | 20 | 27 | 8 | 4 | 8 | 7 | 11 | 5 | 11 | 5 | 17 | 15 | 49 | 6159 |

##### 3.3.2 Count - post QC

Number of barcodes shared between pairs of samples post-QC.

|  | S1 | S2 | S3 | S4 | S5 | S6 | S7 | S8 | S9 | S10 | S11 | S12 | S13 | S14 | S15 | S16 | S17 | S18 |
| --- | --- | --- | --- | --- | --- | --- | --- | --- | --- | --- | --- | --- | --- | --- | --- | --- | --- | --- |
| S1 | 3994 | 7 | 10 | 10 | 3 | 10 | 4 | 1 | 5 | 1 | 6 | 6 | 6 | 2 | 2 | 4 | 4 | 4 |
| S2 | 7 | 6737 | 12 | 12 | 6 | 11 | 6 | 2 | 10 | 5 | 5 | 6 | 5 | 3 | 11 | 2 | 3 | 6 |
| S3 | 10 | 12 | 6327 | 5 | 6 | 18 | 8 | 7 | 0 | 5 | 7 | 9 | 5 | 4 | 5 | 3 | 5 | 2 |
| S4 | 10 | 12 | 5 | 6710 | 5 | 20 | 5 | 1 | 6 | 4 | 9 | 7 | 5 | 2 | 10 | 4 | 10 | 3 |
| S5 | 3 | 6 | 6 | 5 | 3249 | 13 | 2 | 1 | 4 | 2 | 3 | 1 | 4 | 0 | 3 | 5 | 0 | 5 |
| S6 | 10 | 11 | 18 | 20 | 13 | 8226 | 6 | 7 | 4 | 4 | 11 | 6 | 3 | 2 | 12 | 6 | 11 | 14 |
| S7 | 4 | 6 | 8 | 5 | 2 | 6 | 3047 | 1 | 1 | 0 | 2 | 3 | 5 | 0 | 2 | 3 | 4 | 2 |
| S8 | 1 | 2 | 7 | 1 | 1 | 7 | 1 | 1860 | 1 | 3 | 0 | 2 | 2 | 0 | 0 | 2 | 2 | 1 |
| S9 | 5 | 10 | 0 | 6 | 4 | 4 | 1 | 1 | 2971 | 1 | 2 | 0 | 2 | 0 | 1 | 1 | 5 | 2 |
| S10 | 1 | 5 | 5 | 4 | 2 | 4 | 0 | 3 | 1 | 2380 | 2 | 3 | 1 | 1 | 3 | 0 | 3 | 3 |
| S11 | 6 | 5 | 7 | 9 | 3 | 11 | 2 | 0 | 2 | 2 | 3795 | 2 | 2 | 2 | 3 | 5 | 5 | 4 |
| S12 | 6 | 6 | 9 | 7 | 1 | 6 | 3 | 2 | 0 | 3 | 2 | 3098 | 3 | 1 | 1 | 3 | 0 | 4 |
| S13 | 6 | 5 | 5 | 5 | 4 | 3 | 5 | 2 | 2 | 1 | 2 | 3 | 3005 | 0 | 1 | 3 | 4 | 9 |
| S14 | 2 | 3 | 4 | 2 | 0 | 2 | 0 | 0 | 0 | 1 | 2 | 1 | 0 | 919 | 2 | 0 | 2 | 2 |
| S15 | 2 | 11 | 5 | 10 | 3 | 12 | 2 | 0 | 1 | 3 | 3 | 1 | 1 | 2 | 3139 | 5 | 2 | 8 |
| S16 | 4 | 2 | 3 | 4 | 5 | 6 | 3 | 2 | 1 | 0 | 5 | 3 | 3 | 0 | 5 | 3688 | 3 | 4 |
| S17 | 4 | 3 | 5 | 10 | 0 | 11 | 4 | 2 | 5 | 3 | 5 | 0 | 4 | 2 | 2 | 3 | 3854 | 1 |
| S18 | 4 | 6 | 2 | 3 | 5 | 14 | 2 | 1 | 2 | 3 | 4 | 4 | 9 | 2 | 8 | 4 | 1 | 4102 |

##### 3.3.3 Jaccard Index - before QC

Fraction (%) of barcodes shared between pairs of samples pre-QC.

|  | S1 | S2 | S3 | S4 | S5 | S6 | S7 | S8 | S9 | S10 | S11 | S12 | S13 | S14 | S15 | S16 | S17 | S18 |
| --- | --- | --- | --- | --- | --- | --- | --- | --- | --- | --- | --- | --- | --- | --- | --- | --- | --- | --- |
| S1 | 100.00 | 0.20 | 0.27 | 0.29 | 0.13 | 0.22 | 0.22 | 0.10 | 0.13 | 0.09 | 0.18 | 0.13 | 0.23 | 0.07 | 0.12 | 0.19 | 0.20 | 0.14 |
| S2 | 0.20 | 100.00 | 0.25 | 0.28 | 0.31 | 0.21 | 0.15 | 0.10 | 0.26 | 0.27 | 0.27 | 0.16 | 0.17 | 0.08 | 0.26 | 0.18 | 0.18 | 0.20 |
| S3 | 0.27 | 0.25 | 100.00 | 0.13 | 0.18 | 0.31 | 0.21 | 0.25 | 0.06 | 0.11 | 0.22 | 0.21 | 0.17 | 0.10 | 0.16 | 0.13 | 0.12 | 0.11 |
| S4 | 0.29 | 0.28 | 0.13 | 100.00 | 0.24 | 0.35 | 0.13 | 0.07 | 0.14 | 0.17 | 0.28 | 0.21 | 0.14 | 0.09 | 0.30 | 0.14 | 0.18 | 0.22 |
| S5 | 0.13 | 0.31 | 0.18 | 0.24 | 100.00 | 0.33 | 0.16 | 0.17 | 0.19 | 0.13 | 0.18 | 0.18 | 0.19 | 0.02 | 0.15 | 0.16 | 0.12 | 0.28 |
| S6 | 0.22 | 0.21 | 0.31 | 0.35 | 0.33 | 100.00 | 0.19 | 0.19 | 0.17 | 0.18 | 0.25 | 0.23 | 0.13 | 0.06 | 0.22 | 0.18 | 0.22 | 0.31 |
| S7 | 0.22 | 0.15 | 0.21 | 0.13 | 0.16 | 0.19 | 100.00 | 0.05 | 0.12 | 0.09 | 0.18 | 0.12 | 0.19 | 0.06 | 0.14 | 0.15 | 0.17 | 0.14 |
| S8 | 0.10 | 0.10 | 0.25 | 0.07 | 0.17 | 0.19 | 0.05 | 100.00 | 0.09 | 0.21 | 0.08 | 0.07 | 0.18 | 0.00 | 0.14 | 0.15 | 0.07 | 0.08 |
| S9 | 0.13 | 0.26 | 0.06 | 0.14 | 0.19 | 0.17 | 0.12 | 0.09 | 100.00 | 0.17 | 0.09 | 0.02 | 0.12 | 0.03 | 0.04 | 0.11 | 0.19 | 0.14 |
| S10 | 0.09 | 0.27 | 0.11 | 0.17 | 0.13 | 0.18 | 0.09 | 0.21 | 0.17 | 100.00 | 0.14 | 0.17 | 0.07 | 0.04 | 0.22 | 0.12 | 0.21 | 0.14 |
| S11 | 0.18 | 0.27 | 0.22 | 0.28 | 0.18 | 0.25 | 0.18 | 0.08 | 0.09 | 0.14 | 100.00 | 0.14 | 0.15 | 0.11 | 0.25 | 0.17 | 0.19 | 0.18 |
| S12 | 0.13 | 0.16 | 0.21 | 0.21 | 0.18 | 0.23 | 0.12 | 0.07 | 0.02 | 0.17 | 0.14 | 100.00 | 0.17 | 0.21 | 0.12 | 0.19 | 0.08 | 0.09 |
| S13 | 0.23 | 0.17 | 0.17 | 0.14 | 0.19 | 0.13 | 0.19 | 0.18 | 0.12 | 0.07 | 0.15 | 0.17 | 100.00 | 0.03 | 0.08 | 0.18 | 0.14 | 0.21 |
| S14 | 0.07 | 0.08 | 0.10 | 0.09 | 0.02 | 0.06 | 0.06 | 0.00 | 0.03 | 0.04 | 0.11 | 0.21 | 0.03 | 100.00 | 0.12 | 0.06 | 0.06 | 0.13 |
| S15 | 0.12 | 0.26 | 0.16 | 0.30 | 0.15 | 0.22 | 0.14 | 0.14 | 0.04 | 0.22 | 0.25 | 0.12 | 0.08 | 0.12 | 100.00 | 0.15 | 0.08 | 0.30 |
| S16 | 0.19 | 0.18 | 0.13 | 0.14 | 0.16 | 0.18 | 0.15 | 0.15 | 0.11 | 0.12 | 0.17 | 0.19 | 0.18 | 0.06 | 0.15 | 100.00 | 0.18 | 0.26 |
| S17 | 0.20 | 0.18 | 0.12 | 0.18 | 0.12 | 0.22 | 0.17 | 0.07 | 0.19 | 0.21 | 0.19 | 0.08 | 0.14 | 0.06 | 0.08 | 0.18 | 100.00 | 0.85 |
| S18 | 0.14 | 0.20 | 0.11 | 0.22 | 0.28 | 0.31 | 0.14 | 0.08 | 0.14 | 0.14 | 0.18 | 0.09 | 0.21 | 0.13 | 0.30 | 0.26 | 0.85 | 100.00 |

##### 3.3.4 Jaccard Index - post QC

Fraction (%) of barcodes shared between pairs of samples post-QC.

|  | S1 | S2 | S3 | S4 | S5 | S6 | S7 | S8 | S9 | S10 | S11 | S12 | S13 | S14 | S15 | S16 | S17 | S18 |
| --- | --- | --- | --- | --- | --- | --- | --- | --- | --- | --- | --- | --- | --- | --- | --- | --- | --- | --- |
| S1 | 100.00 | 0.13 | 0.19 | 0.19 | 0.08 | 0.16 | 0.11 | 0.03 | 0.14 | 0.03 | 0.15 | 0.17 | 0.17 | 0.08 | 0.06 | 0.10 | 0.10 | 0.10 |
| S2 | 0.13 | 100.00 | 0.18 | 0.18 | 0.12 | 0.15 | 0.12 | 0.05 | 0.21 | 0.11 | 0.09 | 0.12 | 0.10 | 0.08 | 0.22 | 0.04 | 0.06 | 0.11 |
| S3 | 0.19 | 0.18 | 100.00 | 0.08 | 0.13 | 0.25 | 0.17 | 0.17 | 0.00 | 0.11 | 0.14 | 0.19 | 0.11 | 0.11 | 0.11 | 0.06 | 0.10 | 0.04 |
| S4 | 0.19 | 0.18 | 0.08 | 100.00 | 0.10 | 0.27 | 0.10 | 0.02 | 0.12 | 0.09 | 0.17 | 0.14 | 0.10 | 0.05 | 0.20 | 0.08 | 0.19 | 0.06 |
| S5 | 0.08 | 0.12 | 0.13 | 0.10 | 100.00 | 0.23 | 0.06 | 0.04 | 0.13 | 0.07 | 0.09 | 0.03 | 0.13 | 0.00 | 0.09 | 0.14 | 0.00 | 0.14 |
| S6 | 0.16 | 0.15 | 0.25 | 0.27 | 0.23 | 100.00 | 0.11 | 0.14 | 0.07 | 0.08 | 0.18 | 0.11 | 0.05 | 0.04 | 0.21 | 0.10 | 0.18 | 0.23 |
| S7 | 0.11 | 0.12 | 0.17 | 0.10 | 0.06 | 0.11 | 100.00 | 0.04 | 0.03 | 0.00 | 0.06 | 0.10 | 0.17 | 0.00 | 0.06 | 0.09 | 0.12 | 0.06 |
| S8 | 0.03 | 0.05 | 0.17 | 0.02 | 0.04 | 0.14 | 0.04 | 100.00 | 0.04 | 0.14 | 0.00 | 0.08 | 0.08 | 0.00 | 0.00 | 0.07 | 0.07 | 0.03 |
| S9 | 0.14 | 0.21 | 0.00 | 0.12 | 0.13 | 0.07 | 0.03 | 0.04 | 100.00 | 0.04 | 0.06 | 0.00 | 0.07 | 0.00 | 0.03 | 0.03 | 0.15 | 0.06 |
| S10 | 0.03 | 0.11 | 0.11 | 0.09 | 0.07 | 0.08 | 0.00 | 0.14 | 0.04 | 100.00 | 0.06 | 0.11 | 0.04 | 0.06 | 0.11 | 0.00 | 0.10 | 0.09 |
| S11 | 0.15 | 0.09 | 0.14 | 0.17 | 0.09 | 0.18 | 0.06 | 0.00 | 0.06 | 0.06 | 100.00 | 0.06 | 0.06 | 0.08 | 0.09 | 0.13 | 0.13 | 0.10 |
| S12 | 0.17 | 0.12 | 0.19 | 0.14 | 0.03 | 0.11 | 0.10 | 0.08 | 0.00 | 0.11 | 0.06 | 100.00 | 0.10 | 0.05 | 0.03 | 0.09 | 0.00 | 0.11 |
| S13 | 0.17 | 0.10 | 0.11 | 0.10 | 0.13 | 0.05 | 0.17 | 0.08 | 0.07 | 0.04 | 0.06 | 0.10 | 100.00 | 0.00 | 0.03 | 0.09 | 0.12 | 0.25 |
| S14 | 0.08 | 0.08 | 0.11 | 0.05 | 0.00 | 0.04 | 0.00 | 0.00 | 0.00 | 0.06 | 0.08 | 0.05 | 0.00 | 100.00 | 0.10 | 0.00 | 0.08 | 0.08 |
| S15 | 0.06 | 0.22 | 0.11 | 0.20 | 0.09 | 0.21 | 0.06 | 0.00 | 0.03 | 0.11 | 0.09 | 0.03 | 0.03 | 0.10 | 100.00 | 0.15 | 0.06 | 0.22 |
| S16 | 0.10 | 0.04 | 0.06 | 0.08 | 0.14 | 0.10 | 0.09 | 0.07 | 0.03 | 0.00 | 0.13 | 0.09 | 0.09 | 0.00 | 0.15 | 100.00 | 0.08 | 0.10 |
| S17 | 0.10 | 0.06 | 0.10 | 0.19 | 0.00 | 0.18 | 0.12 | 0.07 | 0.15 | 0.10 | 0.13 | 0.00 | 0.12 | 0.08 | 0.06 | 0.08 | 100.00 | 0.03 |
| S18 | 0.10 | 0.11 | 0.04 | 0.06 | 0.14 | 0.23 | 0.06 | 0.03 | 0.06 | 0.09 | 0.10 | 0.11 | 0.25 | 0.08 | 0.22 | 0.10 | 0.03 | 100.00 |

---

#### 3.4 Doublet Cell Detection using Scrublet Scoring System

Observed scores are used for doublet classification. Dashed line indicates the threshold used to identify doublets.

##### 3.4.1 Observed scores

##### 3.4.2 Simulated scores

---

#### 3.5 Impact of Quality Control on Cell, Gene, and UMI Abundances

##### 3.5.1 Number of cells and genes per sample

circle and diamonds refer to before and after QC, respectively.

##### 3.5.2 Average number of UMIs/cell and genes/cell per sample

The error bars represent the standard deviation of the number of UMIs and genes across cells per sample.

---

### 4 ExpressObject

Read me

#### 4.1 Key QC Metrics of Merged Object

|  | min | q0 | q25 | q50 | q75 | q100 | max |
| --- | --- | --- | --- | --- | --- | --- | --- |
| UMI per cell | 750 | 750 | 4,358 | 5,871 | 7,460 | 99,574 | 99,574 |
| Gene per cell | 250 | 250 | 1,387 | 1,643 | 1,936 | 8,324 | 8,324 |
| Mitochondrial (%) | 0.00 | 0.00 | 4.79 | 6.14 | 7.85 | 15.00 | 15.00 |

---

#### 4.2 Dictionary of Computed Metadata Fields and their Descriptions

| Metadata | Description |
| --- | --- |
| `sample_id` | Unique identifier for each sample, used to track each sample. |
| `sample` | Sample name provided by the user, matching the input file or experimental label. |
| `n_genes_by_counts` | Number of detected genes per cell |
| `total_counts` | Total number of UMIs per cell |
| `total_counts_mito` | Total UMIs from mitochondrial genes per cell |
| `pct_counts_mito` | Percentage of mitochondrial UMIs per cell |
| `n_genes` | Number of detected genes per cell |
| `n_counts` | Total number of UMIs per cell |
| `doublet_scores_obs` | Scrublet doublet score for each observed cell |
| `predicted_doublet` | Doublet prediction label (True/False) |
| `leiden_cluster` | Leiden cluster assignment |
| `cellstate_scimilarity` | Predicted cell state from scimilarity |
| `celltype_scimilarity` | Curated cell type using pipeline ontology |
| `cellstate_celltypist` | Predicted cell state from celltypist |
| `celltype_celltypist` | Curated cell type using pipeline ontology |

---

#### 4.3 Visualization of Merged scRNA-Seq Object in 2D Space

##### 4.3.1 UMAP Projection

##### 4.3.2 UMAP Projection (no-batch correction)

##### 4.3.3 TSNE Projection

##### 4.3.4 TSNE Projection (no-batch correction)

---

#### 4.4 Visualizing Cell QC Metrics Using Dimensionality Reduction

##### 4.4.1 UMI per cell - UMAP Projection

##### 4.4.2 Gene per cell - UMAP Projection

##### 4.4.3 Mitochondrial (%) per cell - UMAP Projection

##### 4.4.4 UMI per cell - TSNE Projection

##### 4.4.5 Gene per cell - TSNE Projection

##### 4.4.6 Mitochondrial (%) per cell - TSNE Projection

---

### 5 ExpressCluster

Read me

#### 5.1 Visualizing Cell Populations with Dimensionality Reduction

##### 5.1.1 UMAP Projection

##### 5.1.2 UMAP Projection (no-batch correction)

##### 5.1.3 TSNE Projection

##### 5.1.4 TSNE Projection (no-batch correction)

---

#### 5.2 Cell Composition and Abundance in Clusters

Values at the top of each bar indicate the percentage of cells

##### 5.2.1 Cluster Frequency

##### 5.2.2 Cluster Frequency (no-batch correction)

---

#### 5.3 Sample Imbalance in Clusters Using Gini Indices

The Gini coefficient is used to quantify sample-level or batch-level imbalance within each Leiden cluster. Higher Gini values indicate skewed representation (e.g., one sample dominating a cluster), which may be indicative of batch effects or sampling artifacts. Values next to each point reflect the number of cells per cluster. A horizontal red dashed line at Gini = 0.5 is provided as a visual threshold to aid interpretation.

##### 5.3.1 Dispersion Curve by ‘sample\_id’ (default plotting)

Computed using batch-corrected data (if enabled)

##### 5.3.2 Dispersion Curve by sample\_id (no batch correction)

##### 5.3.3 Dispersion Curve by sample\_id (after batch correction)

---

### 6 ExpressAnnotation

Read me

#### 6.1 Visualizing Distinct Cell Type Populations with Dimensionality Reduction

##### 6.1.1 scimilarity UMAP Projection

##### 6.1.2 scimilarity TSNE Projection

##### 6.1.3 Celltypist UMAP Projection

##### 6.1.4 Celltypist TSNE Projection

---

#### 6.2 Cell Type Population Frequency

Values at the top of each bar indicate the percentage of cells

##### 6.2.1 SCimilarity (cell type)

##### 6.2.2 SCimilarity (cell state)

##### 6.2.3 CellTypist (cell type)

##### 6.2.4 CellTypist (cell state)

---

#### 6.3 Cell Type Population Composition

The curated ontology (cell type) acts as a reference framework, while annotation methods refine classifications into more detailed (cell state) categories. The network visualizes hierarchical mapping from cellstate (real annotation) to celltype (curated).

##### 6.3.1 SCimilarity

##### 6.3.2 CellTypist

---

### 7 ExpressMarkers

Read me

#### 7.1 Expression Signatures Genes for Each Group

Top markers (p\_val\_adj < 0.05 and expressed in more than 0.1 fraction of cells) for each group.

##### 7.1.1 Leiden Cluster

##### 7.1.2 SCimilarity Group

##### 7.1.3 CellTypist Group

---

### 8 ExpressRefs

Read me

Documentation: View Paper

Data Availability: Access Data

---

### 9 ExpressWorkspace

Read me

Computing environment information

#### 9.1 Python

Python Environment (Python 3.12.9 )

| Package | Version |
| --- | --- |
| Brotli | 1.1.0 |
| Deprecated | 1.2.18 |
| GEOparse | 2.0.4 |
| GitPython | 3.1.44 |
| MarkupSafe | 3.0.2 |
| PyPrind | 2.11.3 |
| PySocks | 1.7.1 |
| PyYAML | 6.0.2 |
| Send2Trash | 1.8.3 |
| accelerate | 1.8.1 |
| aiofiles | 24.1.0 |
| aiohappyeyeballs | 2.6.1 |
| aiohttp | 3.12.13 |
| aiosignal | 1.3.2 |
| anndata | 0.11.4 |
| annotated-types | 0.7.0 |
| annoy | 1.17.3 |
| anthropic | 0.54.0 |
| anyio | 4.8.0 |
| archspec | 0.2.3 |
| argon2-cffi | 23.1.0 |
| argon2-cffi-bindings | 21.2.0 |
| array-api-compat | 1.12.0 |
| arrow | 1.3.0 |
| asciitree | 0.3.3 |
| asttokens | 3.0.0 |
| async-lru | 2.0.4 |
| attrs | 25.1.0 |
| babel | 2.17.0 |
| beautifulsoup4 | 4.13.3 |
| bioturing-connector | 1.14.0 |
| bleach | 6.2.0 |
| boltons | 24.0.0 |
| cached-property | 1.5.2 |
| cachetools | 5.5.2 |
| captum | 0.8.0 |
| celltypist | 1.6.3 |
| certifi | 2025.1.31 |
| cffi | 1.17.1 |
| charset-normalizer | 3.4.0 |
| circlify | 0.15.0 |
| click | 8.2.1 |
| colorama | 0.4.6 |
| comm | 0.2.2 |
| conda | 25.1.1 |
| conda-libmamba-solver | 25.1.1 |
| conda-package-handling | 2.3.0 |
| conda-package-streaming | 0.10.0 |
| contourpy | 1.3.2 |
| cycler | 0.12.1 |
| cython | 3.1.2 |
| dataclasses-json | 0.6.7 |
| debugpy | 1.8.12 |
| decorator | 5.1.1 |
| defusedxml | 0.7.1 |
| distro | 1.9.0 |
| entmax | 1.3 |
| et-xmlfile | 2.0.0 |
| exceptiongroup | 1.2.2 |
| executing | 2.1.0 |
| fa2-modified | 0.3.10 |
| fastapi | 0.115.13 |
| fasteners | 0.19 |
| fastjsonschema | 2.21.1 |
| ffmpy | 0.6.0 |
| filelock | 3.18.0 |
| fonttools | 4.58.4 |
| fqdn | 1.5.1 |
| frozendict | 2.4.6 |
| frozenlist | 1.7.0 |
| fsspec | 2025.5.1 |
| gitdb | 4.0.12 |
| google-ai-generativelanguage | 0.6.15 |
| google-api-core | 2.25.1 |
| google-api-python-client | 2.173.0 |
| google-auth | 2.40.3 |
| google-auth-httplib2 | 0.2.0 |
| google-generativeai | 0.8.5 |
| googleapis-common-protos | 1.70.0 |
| gradio | 5.34.2 |
| gradio-client | 1.10.3 |
| graphviz | 0.21 |
| greenlet | 3.2.3 |
| groovy | 0.1.2 |
| grpcio | 1.73.0 |
| grpcio-status | 1.71.0 |
| h11 | 0.14.0 |
| h2 | 4.1.0 |
| h5py | 3.14.0 |
| harmonypy | 0.0.10 |
| hf-xet | 1.1.5 |
| hnswlib | 0.8.0 |
| hpack | 4.0.0 |
| httpcore | 1.0.7 |
| httplib2 | 0.22.0 |
| httpx | 0.28.1 |
| httpx-sse | 0.4.0 |
| huggingface-hub | 0.33.0 |
| hyperframe | 6.0.1 |
| idna | 3.10 |
| igraph | 0.11.9 |
| imageio | 2.37.0 |
| importlib-metadata | 8.6.1 |
| importlib-resources | 6.5.2 |
| ipykernel | 6.29.5 |
| ipython | 8.32.0 |
| isoduration | 20.11.0 |
| jedi | 0.19.2 |
| jinja2 | 3.1.5 |
| jiter | 0.10.0 |
| joblib | 1.5.1 |
| json5 | 0.10.0 |
| jsonpatch | 1.33 |
| jsonpointer | 3.0.0 |
| jsonschema | 4.23.0 |
| jsonschema-specifications | 2024.10.1 |
| jupyter-client | 8.6.3 |
| jupyter-core | 5.7.2 |
| jupyter-events | 0.12.0 |
| jupyter-lsp | 2.2.5 |
| jupyter-server | 2.15.0 |
| jupyter-server-mathjax | 0.2.6 |
| jupyter-server-terminals | 0.5.3 |
| jupyterlab | 4.3.5 |
| jupyterlab-git | 0.51.0 |
| jupyterlab-pygments | 0.3.0 |
| jupyterlab-server | 2.27.3 |
| jupyterlab-widgets | 3.0.13 |
| kiwisolver | 1.4.8 |
| langchain | 0.3.26 |
| langchain-anthropic | 0.3.15 |
| langchain-community | 0.3.26 |
| langchain-core | 0.3.66 |
| langchain-openai | 0.3.24 |
| langchain-text-splitters | 0.3.8 |
| langchainhub | 0.1.21 |
| langgraph | 0.4.8 |
| langgraph-checkpoint | 2.1.0 |
| langgraph-prebuilt | 0.2.2 |
| langgraph-sdk | 0.1.70 |
| langsmith | 0.4.1 |
| lazy-loader | 0.4 |
| legacy-api-wrap | 1.4.1 |
| leidenalg | 0.10.2 |
| libmambapy | 2.0.5 |
| lightning-utilities | 0.14.3 |
| llvmlite | 0.44.0 |
| markdown-it-py | 3.0.0 |
| marshmallow | 3.26.1 |
| matplotlib | 3.10.3 |
| matplotlib-inline | 0.1.7 |
| mdurl | 0.1.2 |
| menuinst | 2.2.0 |
| mistune | 3.1.2 |
| mpmath | 1.3.0 |
| multidict | 6.5.0 |
| mypy-extensions | 1.1.0 |
| narwhals | 1.43.1 |
| natsort | 8.4.0 |
| nbclient | 0.10.2 |
| nbconvert | 7.16.6 |
| nbdime | 4.0.2 |
| nbformat | 5.10.4 |
| nest-asyncio | 1.6.0 |
| networkx | 3.5 |
| nose | 1.3.7 |
| notebook-shim | 0.2.4 |
| numba | 0.61.2 |
| numcodecs | 0.15.1 |
| numpy | 1.26.4 |
| nvidia-cublas-cu12 | 12.6.4.1 |
| nvidia-cuda-cupti-cu12 | 12.6.80 |
| nvidia-cuda-nvrtc-cu12 | 12.6.77 |
| nvidia-cuda-runtime-cu12 | 12.6.77 |
| nvidia-cudnn-cu12 | 9.5.1.17 |
| nvidia-cufft-cu12 | 11.3.0.4 |
| nvidia-cufile-cu12 | 1.11.1.6 |
| nvidia-curand-cu12 | 10.3.7.77 |
| nvidia-cusolver-cu12 | 11.7.1.2 |
| nvidia-cusparse-cu12 | 12.5.4.2 |
| nvidia-cusparselt-cu12 | 0.6.3 |
| nvidia-nccl-cu12 | 2.26.2 |
| nvidia-nvjitlink-cu12 | 12.6.85 |
| nvidia-nvtx-cu12 | 12.6.77 |
| obonet | 1.1.1 |
| openai | 1.90.0 |
| openpyxl | 3.1.5 |
| optree | 0.16.0 |
| orjson | 3.10.18 |
| ormsgpack | 1.10.0 |
| overrides | 7.7.0 |
| packaging | 24.2 |
| pandas | 2.3.0 |
| pandocfilters | 1.5.0 |
| parso | 0.8.4 |
| patsy | 1.0.1 |
| pexpect | 4.9.0 |
| pickleshare | 0.7.5 |
| pillow | 11.2.1 |
| pip | 24.2 |
| pkgutil-resolve-name | 1.3.10 |
| platformdirs | 4.3.6 |
| plotly | 6.1.2 |
| pluggy | 1.5.0 |
| prometheus-client | 0.21.1 |
| prompt-toolkit | 3.0.50 |
| propcache | 0.3.2 |
| proto-plus | 1.26.1 |
| protobuf | 5.29.5 |
| psutil | 6.1.1 |
| ptyprocess | 0.7.0 |
| pure-eval | 0.2.3 |
| pyarrow | 20.0.0 |
| pyasn1 | 0.6.1 |
| pyasn1-modules | 0.4.2 |
| pycosat | 0.6.6 |
| pycparser | 2.22 |
| pydantic | 2.11.7 |
| pydantic-core | 2.33.2 |
| pydantic-settings | 2.10.0 |
| pydub | 0.25.1 |
| pygments | 2.19.1 |
| pymupdf | 1.26.1 |
| pynndescent | 0.5.13 |
| pyparsing | 3.2.3 |
| python-dateutil | 2.9.0.post0 |
| python-dotenv | 1.1.0 |
| python-json-logger | 2.0.7 |
| python-multipart | 0.0.20 |
| pytorch-lightning | 2.5.2 |
| pytz | 2025.1 |
| pyzmq | 26.2.1 |
| referencing | 0.36.2 |
| regex | 2024.11.6 |
| requests | 2.32.3 |
| requests-toolbelt | 1.0.0 |
| rfc3339-validator | 0.1.4 |
| rfc3986-validator | 0.1.1 |
| rich | 14.0.0 |
| rpds-py | 0.22.3 |
| rsa | 4.9.1 |
| ruamel.yaml | 0.18.10 |
| ruamel.yaml.clib | 0.2.8 |
| ruff | 0.12.0 |
| safehttpx | 0.1.6 |
| safetensors | 0.5.3 |
| scanpy | 1.11.2 |
| scikit-image | 0.25.2 |
| scikit-learn | 1.7.0 |
| scimilarity | 0.4.0 |
| scipy | 1.15.3 |
| scrublet | 0.2.3 |
| seaborn | 0.13.2 |
| semantic-version | 2.10.0 |
| session-info2 | 0.1.2 |
| setuptools | 75.1.0 |
| shellingham | 1.5.4 |
| six | 1.17.0 |
| smmap | 5.0.2 |
| sniffio | 1.3.1 |
| solvebio | 2.31.0 |
| soupsieve | 2.5 |
| sqlalchemy | 2.0.41 |
| stack-data | 0.6.3 |
| starlette | 0.46.2 |
| statsmodels | 0.14.4 |
| sympy | 1.14.0 |
| tblib | 1.7.0 |
| tenacity | 9.1.2 |
| terminado | 0.18.1 |
| texttable | 1.7.0 |
| threadpoolctl | 3.6.0 |
| tifffile | 2025.6.11 |
| tiktoken | 0.9.0 |
| tiledb | 0.34.0 |
| tiledb-cloud | 0.13.0 |
| tiledb-vector-search | 0.13.0 |
| tinycss2 | 1.4.0 |
| tokenizers | 0.21.1 |
| tomli | 2.2.1 |
| tomlkit | 0.13.3 |
| torch | 2.7.1 |
| torchaudio | 2.7.1 |
| torchmetrics | 1.7.3 |
| torchopt | 0.7.3 |
| torchsummary | 1.5.1 |
| torchvision | 0.22.1 |
| tornado | 6.4.2 |
| tqdm | 4.66.5 |
| traitlets | 5.14.3 |
| transformers | 4.52.4 |
| triton | 3.3.1 |
| truststore | 0.9.2 |
| typer | 0.16.0 |
| types-python-dateutil | 2.9.0.20241206 |
| types-requests | 2.32.4.20250611 |
| typing-extensions | 4.12.2 |
| typing-inspect | 0.9.0 |
| typing-inspection | 0.4.1 |
| typing-utils | 0.1.0 |
| tzdata | 2025.2 |
| umap-learn | 0.5.7 |
| uri-template | 1.3.0 |
| uritemplate | 4.2.0 |
| urllib3 | 2.2.3 |
| uvicorn | 0.34.3 |
| wcwidth | 0.2.13 |
| webcolors | 24.11.1 |
| webencodings | 0.5.1 |
| websocket-client | 1.8.0 |
| websockets | 15.0.1 |
| wheel | 0.44.0 |
| wrapt | 1.17.2 |
| xxhash | 3.5.0 |
| yarl | 1.20.1 |
| zarr | 2.18.7 |
| zipp | 3.21.0 |
| zstandard | 0.23.0 |
| autocommand | 2.2.2 |
| backports.tarfile | 1.2.0 |
| inflect | 7.3.1 |
| jaraco.collections | 5.1.0 |
| jaraco.context | 5.3.0 |
| jaraco.functools | 4.0.1 |
| jaraco.text | 3.12.1 |
| more-itertools | 10.3.0 |
| typeguard | 4.3.0 |

## 9.2 R

R Environment (R 4.4.1 )

| Package | Version |
| --- | --- |
| abind | 1.4-8 |
| ape | 5.8-1 |
| aplot | 0.2.7 |
| arrow | 17.0.0.1 |
| askpass | 1.2.1 |
| assertthat | 0.2.1 |
| aws.s3 | 0.3.22 |
| aws.signature | 0.6.0 |
| backports | 1.5.0 |
| base64enc | 0.1-3 |
| beachmat | 2.18.1 |
| beeswarm | 0.4.0 |
| BH | 1.87.0-1 |
| Biobase | 2.66.0 |
| BiocGenerics | 0.52.0 |
| BiocManager | 1.30.26 |
| BiocNeighbors | 1.20.2 |
| BiocParallel | 1.36.0 |
| BiocSingular | 1.18.0 |
| BiocVersion | 3.20.0 |
| bit | 4.5.0 |
| bit64 | 4.5.2 |
| bitops | 1.0-9 |
| blob | 1.2.4 |
| bluster | 1.12.0 |
| brew | 1.0-10 |
| brio | 1.1.5 |
| broom | 1.0.7 |
| bslib | 0.8.0 |
| cachem | 1.1.0 |
| Cairo | 1.6-0 |
| callr | 3.7.6 |
| caret | 6.0-93 |
| caTools | 1.18.3 |
| cellranger | 1.1.0 |
| circlize | 0.4.16 |
| classInt | 0.4-10 |
| cli | 3.6.5 |
| clipr | 0.8.0 |
| clock | 0.7.1 |
| clue | 0.3-66 |
| colorspace | 2.1-1 |
| commonmark | 1.9.2 |
| ComplexHeatmap | 2.22.0 |
| ConfigParser | 1.0.0 |
| cowplot | 1.1.3 |
| cpp11 | 0.5.2 |
| crayon | 1.5.3 |
| credentials | 2.0.2 |
| crosstalk | 1.2.1 |
| curl | 5.2.3 |
| data.table | 1.17.6 |
| data.tree | 1.1.0 |
| datamods | 1.5.3 |
| DBI | 1.2.3 |
| dbplyr | 2.5.0 |
| DelayedArray | 0.32.0 |
| DelayedMatrixStats | 1.28.1 |
| deldir | 2.0-4 |
| Deriv | 4.2.0 |
| desc | 1.4.3 |
| DescTools | 0.99.60 |
| devtools | 2.4.5 |
| diagram | 1.6.5 |
| diffobj | 0.3.5 |
| digest | 0.6.37 |
| distributional | 0.5.0 |
| domino | 0.3.1 |
| DominoDataCapture | 0.1.1 |
| DominoDataR | 0.2.4 |
| doParallel | 1.0.17 |
| dotCall64 | 1.2 |
| downlit | 0.4.4 |
| dplyr | 1.1.4 |
| dqrng | 0.4.1 |
| DT | 0.33 |
| dtplyr | 1.3.1 |
| e1071 | 1.7-16 |
| edgeR | 4.4.2 |
| ellipsis | 0.3.2 |
| esquisse | 1.1.2 |
| evaluate | 1.0.1 |
| Exact | 3.3 |
| expm | 1.0-0 |
| fansi | 1.0.6 |
| farver | 2.1.2 |
| fastDummies | 1.7.5 |
| fastmap | 1.2.0 |
| fastmatch | 1.1-6 |
| feather | 0.3.5 |
| fgsea | 1.35.3 |
| fitdistrplus | 1.2-2 |
| FNN | 1.1.4.1 |
| fontawesome | 0.5.2 |
| forcats | 1.0.0 |
| foreach | 1.5.2 |
| forge | 0.2.0 |
| formatR | 1.14 |
| fs | 1.6.6 |
| futile.logger | 1.4.3 |
| futile.options | 1.0.1 |
| future | 1.34.0 |
| future.apply | 1.11.2 |
| gargle | 1.5.2 |
| generics | 0.1.4 |
| GenomeInfoDb | 1.42.3 |
| GenomeInfoDbData | 1.2.13 |
| GenomicRanges | 1.58.0 |
| gert | 2.1.2 |
| getopt | 1.20.4 |
| GetoptLong | 1.0.5 |
| ggbeeswarm | 0.7.2 |
| ggdist | 3.3.3 |
| ggfun | 0.1.9 |
| ggplot2 | 3.5.2 |
| ggplotify | 0.1.2 |
| ggrastr | 1.0.2 |
| ggrepel | 0.9.6 |
| ggridges | 0.5.6 |
| ggtree | 3.10.1 |
| gh | 1.4.1 |
| git2r | 0.33.0 |
| gitcreds | 0.1.2 |
| gld | 2.6.7 |
| glmnet | 4.1-4 |
| GlobalOptions | 0.1.2 |
| globals | 0.16.3 |
| glue | 1.8.0 |
| goftest | 1.2-3 |
| googledrive | 2.1.1 |
| googlesheets4 | 1.1.1 |
| gower | 1.0.1 |
| gplots | 3.2.0 |
| gridExtra | 2.3 |
| gridGraphics | 0.5-1 |
| gridtext | 0.1.5 |
| gtable | 0.3.6 |
| gtools | 3.9.5 |
| hardhat | 1.4.0 |
| harmony | 1.2.3 |
| haven | 2.5.4 |
| hdf5r | 1.3.12 |
| here | 1.0.1 |
| hexbin | 1.28.5 |
| highr | 0.11 |
| hms | 1.1.3 |
| htmltools | 0.5.8.1 |
| htmlwidgets | 1.6.4 |
| httpuv | 1.6.15 |
| httr | 1.4.7 |
| httr2 | 1.0.5 |
| ica | 1.0-3 |
| ids | 1.0.1 |
| igraph | 2.1.4 |
| ini | 0.3.1 |
| ipred | 0.9-15 |
| IRanges | 2.40.1 |
| irlba | 2.3.5.1 |
| isoband | 0.2.7 |
| iterators | 1.0.14 |
| jpeg | 0.1-11 |
| jquerylib | 0.1.4 |
| jsonlite | 2.0.0 |
| kableExtra | 1.4.0 |
| kernlab | 0.9-31 |
| knitr | 1.44 |
| ks | 1.15.1 |
| labeling | 0.4.3 |
| lambda.r | 1.2.4 |
| later | 1.3.2 |
| lava | 1.8.0 |
| lazyeval | 0.2.2 |
| leidenbase | 0.1.35 |
| lgr | 0.4.4 |
| lifecycle | 1.0.4 |
| limma | 3.62.2 |
| listenv | 0.9.1 |
| lme4 | 1.1-37 |
| lmom | 3.2 |
| lmtest | 0.9-40 |
| locfit | 1.5-9.12 |
| lubridate | 1.8.0 |
| magrittr | 2.0.3 |
| markdown | 1.1 |
| MatrixGenerics | 1.18.1 |
| matrixStats | 1.5.0 |
| mclust | 6.1.1 |
| memoise | 2.0.1 |
| metapod | 1.10.1 |
| mime | 0.12 |
| miniUI | 0.1.1.1 |
| minqa | 1.2.8 |
| mlflow | 2.19.0 |
| ModelMetrics | 1.2.2.2 |
| modelr | 0.1.11 |
| multicool | 1.0.1 |
| munsell | 0.5.1 |
| mvtnorm | 1.3-3 |
| Nebulosa | 1.16.0 |
| neuralnet | 1.44.2 |
| nloptr | 2.2.1 |
| numDeriv | 2016.8-1.1 |
| openssl | 2.2.2 |
| optparse | 1.7.5 |
| parallelDist | 0.2.6 |
| parallelly | 1.38.0 |
| patchwork | 1.3.1 |
| pbapply | 1.7-2 |
| pbmcapply | 1.5.1 |
| pheatmap | 1.0.13 |
| phosphoricons | 0.2.1 |
| pillar | 1.10.2 |
| pkgbuild | 1.4.4 |
| pkgconfig | 2.0.3 |
| pkgdown | 2.1.1 |
| pkgload | 1.4.0 |
| plotly | 4.11.0 |
| plumber | 1.2.2 |
| plyr | 1.8.9 |
| png | 0.1-8 |
| polyclip | 1.10-7 |
| pracma | 2.4.4 |
| praise | 1.0.0 |
| prettyunits | 1.2.0 |
| pROC | 1.18.5 |
| processx | 3.8.4 |
| prodlim | 2024.06.25 |
| profvis | 0.4.0 |
| progress | 1.2.3 |
| progressr | 0.15.1 |
| promises | 1.3.0 |
| proxy | 0.4-27 |
| ps | 1.8.0 |
| purrr | 1.0.4 |
| quadprog | 1.5-8 |
| R.methodsS3 | 1.8.2 |
| R.oo | 1.26.0 |
| R.utils | 2.13.0 |
| R6 | 2.6.1 |
| ragg | 1.3.3 |
| randomForest | 4.7-1.1 |
| RANN | 2.6.2 |
| rappdirs | 0.3.3 |
| rbibutils | 2.3 |
| rcmdcheck | 1.4.0 |
| RColorBrewer | 1.1-3 |
| Rcpp | 1.0.14 |
| Rcpp11 | 3.1.2.0.1 |
| RcppAnnoy | 0.0.22 |
| RcppArmadillo | 14.4.3-1 |
| RcppEigen | 0.3.4.0.2 |
| RcppHNSW | 0.6.0 |
| RcppParallel | 5.1.10 |
| RcppProgress | 0.4.2 |
| RcppTOML | 0.2.2 |
| RCurl | 1.98-1.17 |
| Rdpack | 2.6.4 |
| reactable | 0.4.4 |
| reactR | 0.6.1 |
| readr | 2.1.5 |
| readxl | 1.4.5 |
| recipes | 1.1.0 |
| reformulas | 0.4.1 |
| rematch | 2.0.0 |
| rematch2 | 2.1.2 |
| remotes | 2.5.0 |
| reprex | 2.1.1 |
| reshape2 | 1.4.4 |
| reticulate | 1.42.0 |
| rgdal | 1.5-32 |
| RhpcBLASctl | 0.23-42 |
| rio | 1.2.3 |
| rJava | 1.0-11 |
| RJDBC | 0.2-10 |
| rjson | 0.2.23 |
| RJSONIO | 2.0.0 |
| rlang | 1.1.6 |
| rmarkdown | 2.28 |
| ROCR | 1.0-11 |
| rootSolve | 1.8.2.4 |
| roxygen2 | 7.2.1 |
| rprojroot | 2.0.4 |
| RSpectra | 0.16-2 |
| rstudioapi | 0.16.0 |
| rsvd | 1.0.5 |
| Rtsne | 0.17 |
| rversions | 2.1.2 |
| rvest | 1.0.4 |
| S4Arrays | 1.6.0 |
| S4Vectors | 0.44.0 |
| sass | 0.4.9 |
| ScaledMatrix | 1.10.0 |
| scales | 1.4.0 |
| scattermore | 1.2 |
| SCEVAN | 1.0.3 |
| scran | 1.30.2 |
| sctransform | 0.4.2 |
| scuttle | 1.12.0 |
| selectr | 0.4-2 |
| sessioninfo | 1.2.2 |
| Seurat | 5.3.0 |
| SeuratObject | 5.1.0 |
| shape | 1.4.6.1 |
| shiny | 1.9.1 |
| shinybusy | 0.3.3 |
| shinyWidgets | 0.8.7 |
| SingleCellExperiment | 1.28.1 |
| sitmo | 2.0.2 |
| snow | 0.4-4 |
| sodium | 1.3.2 |
| sourcetools | 0.1.7-1 |
| sp | 2.1-4 |
| spam | 2.11-1 |
| SparseArray | 1.6.2 |
| sparseMatrixStats | 1.18.0 |
| spatstat.data | 3.1-6 |
| spatstat.explore | 3.4-3 |
| spatstat.geom | 3.4-1 |
| spatstat.random | 3.4-1 |
| spatstat.sparse | 3.1-0 |
| spatstat.univar | 3.1-3 |
| spatstat.utils | 3.1-4 |
| SQUAREM | 2021.1 |
| statmod | 1.5.0 |
| stringi | 1.8.7 |
| stringr | 1.5.1 |
| SummarizedExperiment | 1.36.0 |
| svglite | 2.2.1 |
| swagger | 5.17.14.1 |
| sys | 3.4.3 |
| systemfonts | 1.2.3 |
| tensor | 1.5.1 |
| testthat | 3.2.1.1 |
| textshaping | 0.4.0 |
| tibble | 3.3.0 |
| tidyr | 1.3.1 |
| tidyselect | 1.2.1 |
| tidytree | 0.4.6 |
| tidyverse | 1.3.2 |
| timechange | 0.3.0 |
| timeDate | 4041.110 |
| tinytex | 0.53 |
| toastui | 0.3.4 |
| treeio | 1.26.0 |
| triebeard | 0.4.1 |
| tzdb | 0.4.0 |
| UCSC.utils | 1.2.0 |
| urlchecker | 1.0.1 |
| urltools | 1.7.3 |
| usethis | 2.1.6 |
| utf8 | 1.2.6 |
| uuid | 1.2-1 |
| uwot | 0.2.3 |
| vcd | 1.4-10 |
| vctrs | 0.6.5 |
| versions | 0.3 |
| vipor | 0.4.7 |
| viridis | 0.6.5 |
| viridisLite | 0.4.2 |
| visNetwork | 2.1.2 |
| vroom | 1.6.5 |
| waldo | 0.5.3 |
| webutils | 1.2.2 |
| whisker | 0.4.1 |
| withr | 3.0.2 |
| writexl | 1.5.1 |
| xfun | 0.48 |
| xml2 | 1.3.6 |
| xopen | 1.0.1 |
| xtable | 1.8-4 |
| XVector | 0.46.0 |
| yaGST | 2017.08.25 |
| yaml | 2.3.10 |
| yulab.utils | 0.2.0 |
| zeallot | 0.1.0 |
| zip | 2.3.1 |
| zlibbioc | 1.52.0 |
| zoo | 1.8-12 |
| base | 4.4.1 |
| boot | 1.3-30 |
| class | 7.3-22 |
| cluster | 2.1.6 |
| codetools | 0.2-19 |
| compiler | 4.4.1 |
| datasets | 4.4.1 |
| foreign | 0.8-86 |
| graphics | 4.4.1 |
| grDevices | 4.4.1 |
| grid | 4.4.1 |
| KernSmooth | 2.23-24 |
| lattice | 0.22-5 |
| MASS | 7.3-61 |
| Matrix | 1.6-5 |
| methods | 4.4.1 |
| mgcv | 1.9-1 |
| nlme | 3.1-165 |
| nnet | 7.3-19 |
| parallel | 4.4.1 |
| rpart | 4.1.23 |
| spatial | 7.3-15 |
| splines | 4.4.1 |
| stats | 4.4.1 |
| stats4 | 4.4.1 |
| survival | 3.7-0 |
| tcltk | 4.4.1 |
| tools | 4.4.1 |
| utils | 4.4.1 |

---
